# Decoding the Language of Chickens - An Innovative NLP Approach to Enhance Poultry Welfare

**DOI:** 10.1101/2024.04.29.591707

**Authors:** Suresh Neethirajan

## Abstract

This research investigates the utilization of the Natural Language Processing-based WHISPER model for decoding chicken vocalizations, with the goal of comprehending the semantics and emotions embedded in their vocal communications. By leveraging advanced acoustic analysis techniques, the study focuses on interpreting the syntax and temporal patterns inherent in the vocalizations to discern the underlying affective states of chickens. This approach facilitates a non-invasive method of monitoring poultry welfare, enhancing traditional animal welfare assessments which often rely on direct human observation and can induce stress in the animals. The principal results from the application of the WHISPER model demonstrate its efficacy in classifying various chicken vocalizations into distinct categories that reflect specific emotional states such as distress, contentment, and fear. This classification is achieved by analyzing the frequency, duration, and intensity of vocalizations, thus providing a detailed insight into the emotional well-being of the animals. Our findings indicate that real-time monitoring of chicken vocalizations using NLP techniques can significantly improve the responsiveness and precision of welfare interventions. This method reduces the need for human interaction, minimizes stress for the animals, and allows for the continuous assessment of their well-being in a farming environment. Furthermore, the research highlights the potential of NLP tools in recognizing and interpreting complex animal vocalizations, which could lead to advancements in automated animal welfare monitoring systems. This study underscores the transformative potential of integrating sophisticated computational models like the WHISPER NLP model into animal welfare practices. By providing a more humane and efficient approach to monitoring animal welfare, this research contributes to the broader field of precision livestock farming, suggesting a shift towards more scientifically informed and welfare-centric farming practices. The application of such technologies not only aids in the immediate improvement of animal welfare but also supports sustainable farming operations by promoting the health and productivity of poultry through enhanced welfare standards.

## 1. Introduction

Understanding the vocalizations of poultry, particularly chickens, is pivotal for several reasons. These vocalizations serve as primary communication channels among chickens, signaling a spectrum of states from contentment to distress. Such vocal cues are essential for the identification of immediate threats or long-term welfare concerns, including disease or social stress. The welfare implications of poultry vocalizations extend beyond ethical considerations, impacting productivity and quality in the farming industry significantly (Neethirajan, 2023; Marino, 2017; Hernandez et al., 2022; Rollin, 2021).

The practice of animal welfare monitoring has undergone significant evolution over the past decades. Initially, the assessment of animal welfare was predominantly based on traditional methodologies that include visual inspections, physiological measurements, and behavioral tests. These methods have provided foundational insights into the well-being of animals, facilitating the development of standards and practices aimed at enhancing animal care.

Visual inspections have been a cornerstone of welfare monitoring, relying on the observation of physical condition, behavior, and environment. However, this method’s effectiveness is contingent on the observer’s experience and can vary significantly, introducing subjective bias into welfare assessments. Additionally, the presence of human observers can alter the natural behavior of animals, potentially masking signs of distress or discomfort (Neethirajan, 2020).

Physiological indicators, such as cortisol levels, heart rate, and immune function, offer objective data on the animal’s stress and health status. Despite their objectivity, these methods often require invasive procedures to collect samples, which can itself induce stress in animals. Moreover, the interpretation of these measurements requires specialized knowledge, limiting their applicability in diverse farming contexts (Vasdal et al., 2022).

Behavioral tests aim to evaluate the affective states, preferences, and social interactions of animals. While these tests provide valuable insights into animal welfare, they are time-consuming and often necessitate specific environmental conditions to be effectively conducted. The subjective interpretation of behavioral outcomes presents an additional challenge, potentially leading to inconsistencies in welfare assessment (Neethirajan, 2022).

The introduction of acoustic analysis technologies marks a significant advancement in the field of animal welfare monitoring. This innovative approach leverages the vocalizations of animals, such as chickens, to gain insights into their affective states in a non-invasive manner.

Acoustic analysis enables the continuous monitoring of vocalizations, offering real-time insights into the welfare of animals. This technology can detect subtle changes in vocal patterns that may indicate stress, discomfort, or well-being, providing a dynamic and responsive tool for welfare assessment. Unlike traditional methods, acoustic analysis minimizes human-animal interaction, reducing the potential for stress-induced behavior changes and allowing for the observation of animals in their natural state (Neethirajan and Kemp, 2021).

One of the key advantages of acoustic analysis is its non-invasive nature. It does not require physical contact with the animals, thereby eliminating the stress associated with handling and sampling. This aspect is particularly beneficial in large-scale farming operations, where the welfare monitoring of individual animals using traditional methods is impractical (Pijpers et al., 2022).

Acoustic analysis goes beyond mere detection of vocalizations; it provides insights into the emotional lives of animals. By analyzing the frequency, duration, and intensity of vocalizations, researchers can infer the emotional (affective) states of chickens, distinguishing between contentment, fear, and distress. This level of understanding is critical for the development of welfare-enhancing interventions and management practices, contributing to the overall well-being of animals (Ferreira et al., 2021). By offering a sophisticated, non-invasive, and objective means to understand the affective states of animals, acoustic analysis stands as a valuable tool in the continuous effort to improve animal welfare. This advancement underscores the importance of adopting innovative technologies in the pursuit of humane and effective animal care practices.

## 2. The Language of Chickens

Chickens communicate through a complex system of vocalizations, which are integral to their social organization, environmental interactions, and welfare indicators. This communicative complexity is crucial for conveying information regarding threats, food sources, social hierarchy, and emotional well-being, necessitating a comprehensive understanding for effective poultry welfare and management strategies (Nicol, 2015).

The categorization of chicken vocalizations into alarm calls, food calls, comfort, and distress calls (Laurijs et al., 2021) has provided profound insights into their behavioral and welfare implications. Alarm calls are paramount for the survival of the flock, serving as an auditory signal to warn of predator presence (Zhang et al., 2023). The differentiation of these calls into aerial versus terrestrial predator-specific sounds reveals an advanced level of acoustic complexity and specificity (Burnham, 2023). Such vocal distinctions indicate chickens’ capacity for nuanced communication, enabling them to convey precise information about the nature of threats, thereby facilitating appropriate flock responses (Ferreira et al., (2024); Cartmill, 2023). This specificity underscores an evolved communicative precision, vital for the adaptive survival strategies within avian species.

Food calls represent a critical communication tool, mediating the balance between competition and cooperation within the flock’s social structure (Bessa Ferreira et al., 2022). These vocalizations not only signal the presence of a food source but also embody the communal aspect of foraging behavior (Marler, 2004). By attracting flock mates to food, these calls reinforce social bonds and cohesion, illustrating a sophisticated interplay between individual needs and group dynamics (Marler, 2004; Du et al., 2018). The dual role of food calls in facilitating communal foraging while maintaining social order highlights the complex social intelligence inherent in chickens (Burish et al., 2004).

The dichotomy between comfort and distress calls offers a window into the emotional states of chickens. Comfort calls, typically emitted during peaceful activities such as nesting or preening, reflect a state of contentment and security (Bestman, 2022). Conversely, distress calls signal discomfort, fear, or pain, serving as an immediate indicator of welfare concerns (Papageorgiou et al., 2023; Manteuffel et al., 2004). The ability to monitor and interpret these vocalizations provides invaluable insights into the health, environmental quality, and social well-being of the flock, allowing for timely interventions to mitigate stressors (Neethirajan, 2023).

Vocalizations are pivotal in the establishment and maintenance of social hierarchies, regulating access to resources and reducing conflict, while also maintaining group cohesion and facilitating social learning (Koshiba et al., 2013; Freire, 2020; Millman and Duncan, 2001; Fichtel and Manser, 2010; Vernes et al., 2021; Ginovart-Panisello et al., 2020).

Traditional approaches to animal welfare assessment have predominantly relied on visual inspections, physiological measurements, and behavioral tests. These methodologies, while foundational in the field of animal welfare science, present several limitations and challenges that can impact their effectiveness and reliability.

Traditional welfare assessment methods are often criticized for their labor-intensive nature and the potential stress they impose on animals. Visual inspections as mentioned above, for example, require continuous human presence, which can alter the natural behavior of chickens, leading to potential biases in welfare assessment. One of the primary challenges associated with traditional welfare assessment methods is their reactive nature. These methods often identify welfare issues only after they have manifested physically or behaviorally, which may be too late for effective intervention. Additionally, the need for specialized training to accurately interpret behavioral cues and physiological data can limit these methods’ applicability across different farming contexts.

The advent of technological advancements, particularly in acoustic analysis, offers promising opportunities to overcome many of these limitations. Acoustic analysis allows for continuous, non-invasive monitoring of chicken vocalizations, providing real-time insights into the welfare state of the flock without the need for direct human observation or invasive procedures. This approach not only reduces the potential stress on animals but also enables the early detection of distress signals, facilitating timely interventions to mitigate welfare issues.

The transition from manual observation to automated systems marks a significant evolution in the field of animal welfare monitoring. Technological advancements, particularly in the domain of acoustic analysis, have revolutionized our ability to assess and understand the affective states of chickens. Automated systems equipped with acoustic analysis capabilities can continuously monitor the vocalizations of chickens, decoding subtle changes in sound patterns that may indicate stress, discomfort, or disease. This method of welfare assessment is significantly less invasive than traditional approaches, allowing for the continuous monitoring of animal welfare without direct interference. Moreover, acoustic analysis can capture a wide range of vocalizations, from distress calls to contentment signals, providing a comprehensive picture of the flock’s welfare state (Neethirajan and Kemp, 2021; Pijpers et al., 2022).

The role of acoustic analysis extends beyond mere detection of vocalizations; it offers insights into the complex emotional lives of chickens. By analyzing the frequency, amplitude, and temporal patterns of vocalizations, researchers can infer the affective states of chickens, distinguishing between states of contentment, fear, stress, and social cohesion. This nuanced understanding of chicken vocalizations and their implications for welfare is a leap forward in animal welfare science, enabling more empathetic and effective management practices.

The integration of Artificial Intelligence (AI) and Machine Learning (ML) in bioacoustic research represents a cutting-edge approach to deciphering the complex vocalizations of chickens. These technologies hold the potential to transform poultry welfare monitoring by providing deep insights into the communicative behaviors of chickens. AI and ML algorithms can analyze vast datasets of chicken vocalizations, identifying patterns and correlations that may be imperceptible to the human ear. By training these algorithms on labeled datasets of known vocalizations, researchers can develop models capable of classifying sounds into categories such as alarm calls, food calls, and distress signals with high accuracy (Cai et al., 2023).

The application of AI and ML in interpreting chicken vocalizations offers a proactive approach to welfare monitoring. These technologies can detect subtle changes in vocalization patterns that signal the onset of stress or disease, often before visible signs emerge. Moreover, the ability of AI and ML to continuously learn and adapt to new data ensures that these systems become increasingly accurate and reliable over time, enhancing their utility in real-world farming operations. These technologies offer a more humane, efficient, and effective approach to understanding and improving the welfare of chickens, emphasizing the importance of continuous innovation in the pursuit of animal welfare excellence.

Table 1 presents a succinct overview of pivotal research outcomes, methodological frameworks, and consequential insights within the realm of poultry and birds bioacoustics analysis and species identification, underscored by the utilization of sophisticated computational methodologies. Each entry in the table captures the core findings and contributions of the corresponding study, underscoring the varied methodologies and their applications in bioacoustics and animal behavior studies. This compilation serves to underline the extensive diversity and innovative approaches permeating current research in these fields.

**Table 1.**
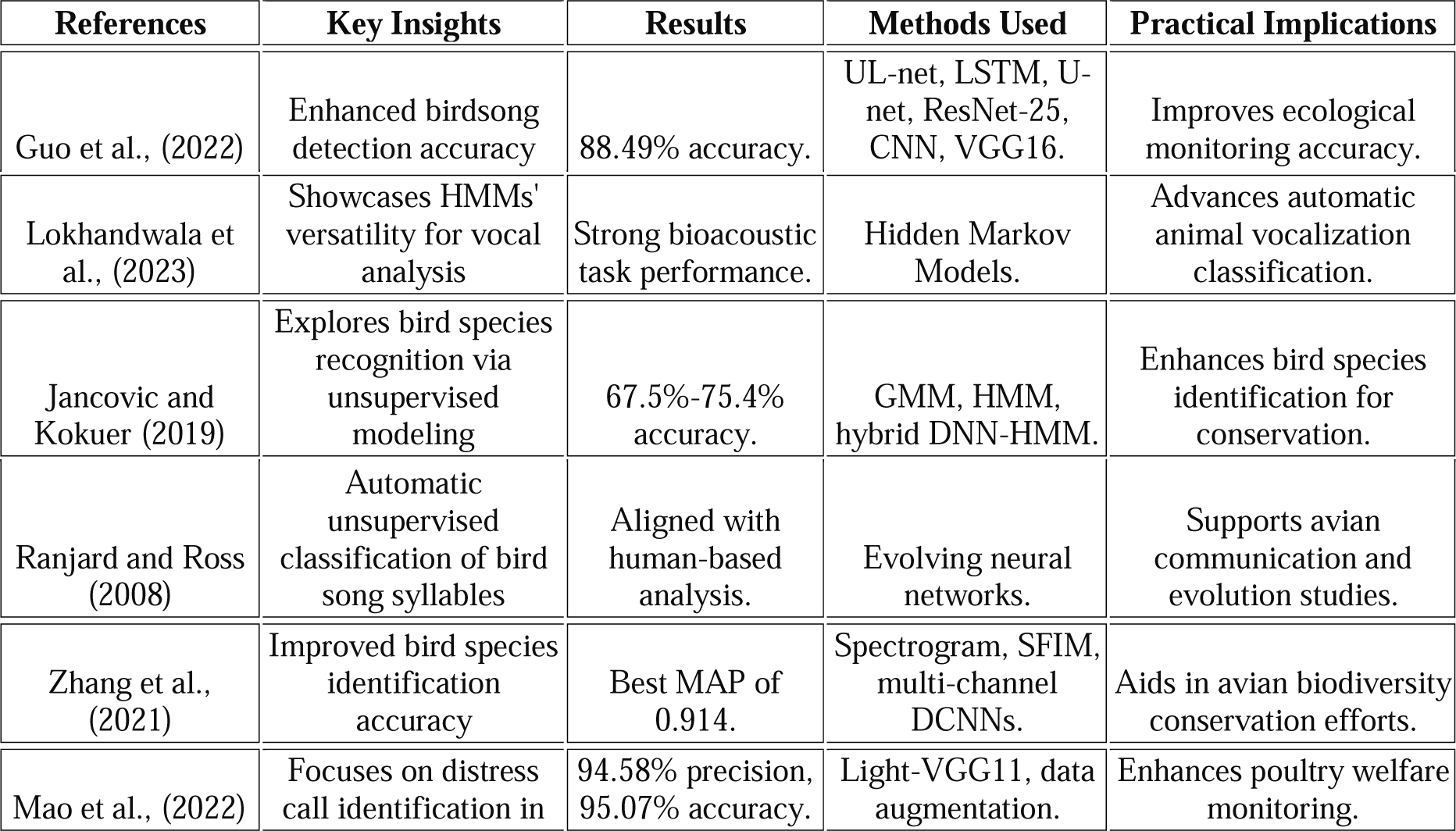
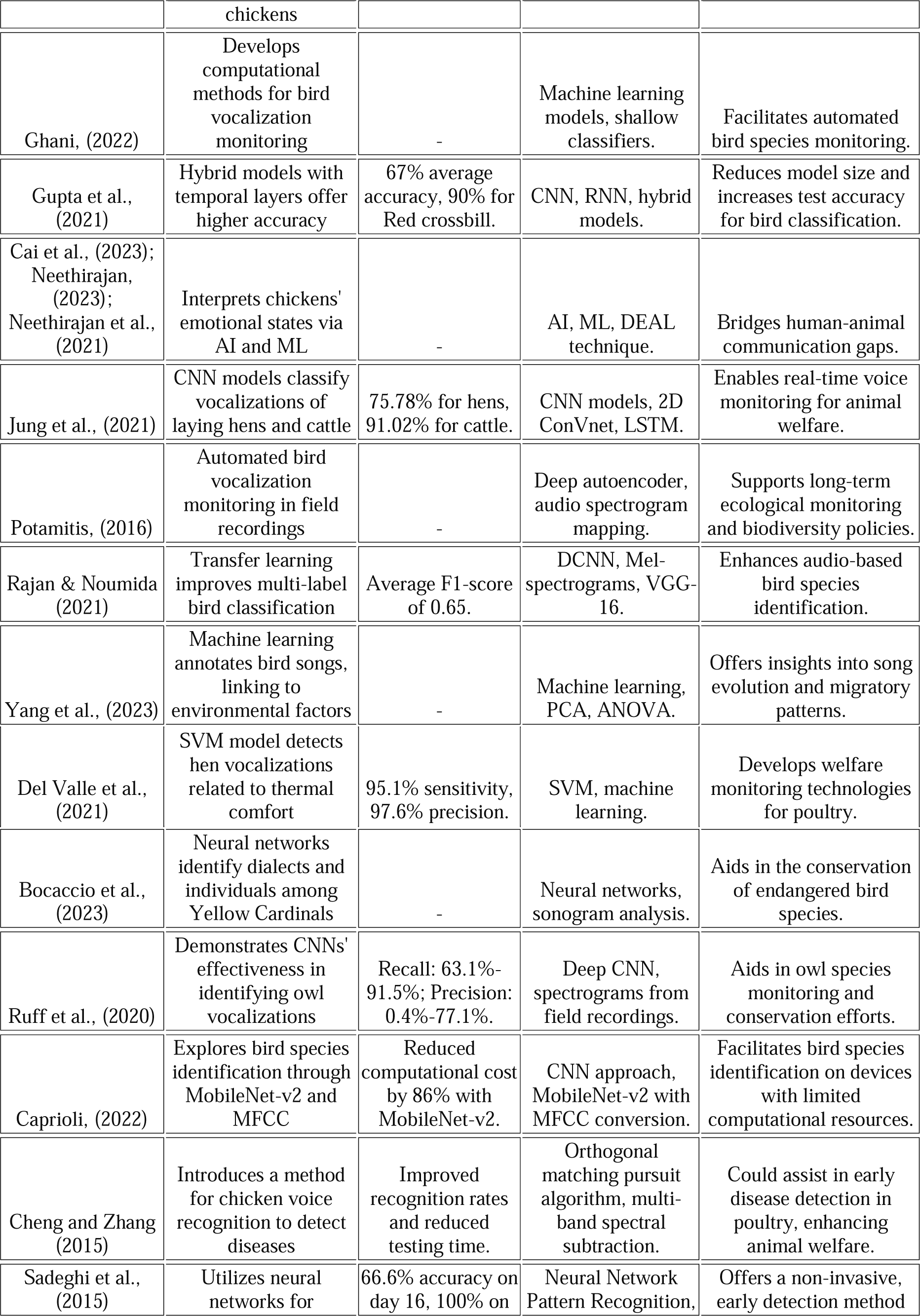
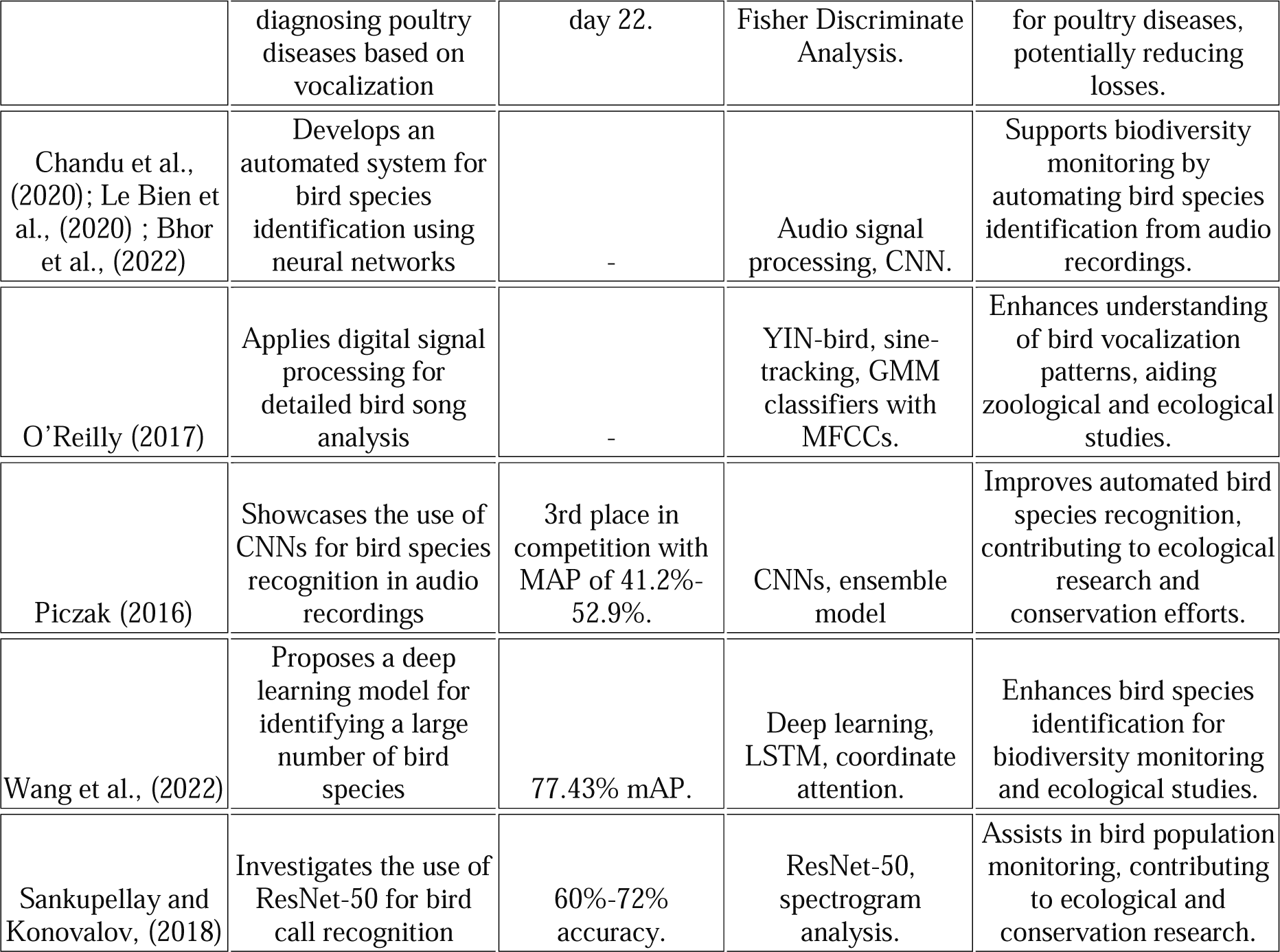
Overview of Avian Vocalization analysis and bird Species ID Research.

## 3. The Whisper NLP Model for Poultry Vocalization Analysis

### 3.1. Understanding the Whisper NLP Model

Whisper, an innovative Automated Speech Recognition (ASR) model released by OpenAI in 2022, has significantly advanced the field with its ability to accurately interpret speech across a multitude of languages and under various audio conditions. Utilizing a Transformer-based architecture, known for its effectiveness in sequential data processing and capturing complex dependencies, Whisper excels in ASR by adeptly managing diverse accents, dialects, and styles, even in noisy environments. Its design for multilingual support enhances its utility for global applications, making it invaluable for international platforms and content with diverse linguistic inputs. The integration of sophisticated neural network architectures allows Whisper to navigate the intricacies of language, demonstrating unparalleled proficiency in not just human speech recognition but also in translating nuanced linguistic patterns (Vachhani et al., 2023; Hasan et al., 2023; Goldstein et al., 2023; Xia et al., 2023; Yeo et al., 2023; Zhang et al., 2023). This adaptability suggests its potential for broader applications, including analyzing animal vocalizations, marking Whisper as a pinnacle of ASR technology that embodies OpenAI’s commitment to creating tools that offer precision and versatility across a spectrum of linguistic and auditory challenges.

### 3.2. Adapting NLP Models for Animal Vocalization Analysis

Extending the capabilities of NLP models like Whisper to the analysis of animal vocalizations represents an innovative cross-disciplinary application. The underlying neural networks that enable the model to process human speech can similarly be trained to recognize and interpret the vocal patterns of chickens and other animals. This approach necessitates a substantial dataset of annotated animal sounds, where each vocalization is linked to specific behaviors, emotional states, or environmental conditions (Stowell et al., 2019). By leveraging these datasets, the model could learn to identify distinct vocal signatures associated with stress, contentment, or social interactions among poultry, akin to the processing of human emotions and sentiments through speech.

### 3.3. Potential for ‘Translating’ Chicken Vocalizations

The application of the Whisper NLP model to chicken vocalizations opens the door to ‘translating’ these sounds into a format comprehensible to humans. This capability could revolutionize computational bioacoustics and animal welfare science by providing a novel method for assessing the emotional and psychological states of chickens. Integrating machine learning, audio processing, and sentiment analysis, this approach could identify specific vocal patterns indicative of distress, well-being, or social harmony (Dogan and Akbulut, 2023). Such insights would not only enhance our understanding of poultry behavior but also inform more empathetic and responsive animal management practices.

### 3.4. NLP Models and Their Potential Applications in Animal Behavior Studies

The prospective application of NLP models like Whisper in animal behavior studies extends beyond mere translation. These models could facilitate real-time monitoring and analysis of vocalizations, enabling prompt detection and response to welfare issues. For instance, changes in the frequency or intensity of distress calls could trigger automatic alerts, prompting immediate intervention by animal caretakers or farmers (Webster and Margerison, 2022). Moreover, the adaptability of NLP models to different ‘dialects’ or vocal patterns across chicken breeds highlights the potential for breed-specific welfare assessments. Such precision could lead to more tailored and effective management strategies, optimizing environmental conditions and social arrangements to suit the vocal communication styles of specific flocks.

The Whisper model, initially designed for human speech recognition, harnesses deep neural networks to understand and translate languages across various dialects with remarkable accuracy. Its application to animal sounds involves training the model on datasets of animal vocalizations annotated with behaviors, emotional states, or environmental contexts. This process enables Whisper to recognize and categorize distinct vocal patterns, similar to how it processes human languages.

The novelty in applying Whisper to animal vocalizations lies in its ability to provide detailed insights into the affective states and social dynamics of animals, potentially transforming areas like wildlife conservation, animal welfare monitoring, and the study of animal behavior. The methodological approach includes recording animal sounds, annotating them with expert knowledge, and then using these annotated datasets to fine-tune Whisper’s algorithms for specific animal species.

This application represents a significant advancement in computational bioacoustics, showcasing the model’s versatility and the potential for cross-disciplinary innovation. The uniqueness of this approach is in its ability to extend state-of-the-art NLP techniques, traditionally used for human language processing, to the realm of animal communication, offering a novel perspective on understanding non-human languages and contributing to our knowledge of animal cognition and social structures. Table 2 provides a synthesis of research integrating the Whisper model alongside various machine learning approaches for audio-to-text conversion and advanced speech recognition tasks. Highlighting key methodologies, breakthrough insights, and technical innovations, the Table 2 showcases the broad utility and impact of these technologies in understanding human and animal communications.

**Table 2.** Diverse Applications of the Whisper Model and Machine Learning in Audio Processing.

| Focus | Methods / Approach | Key Insights | Novelty / Advancement | Technical Info | Species | Reference |
| --- | --- | --- | --- | --- | --- | --- |
| Audio-to-text conversion using Whisper model | Integration of Whisper model into an audio-to-text software tool | Selection of Whisper models based on transcription speed and accuracy | Methodology for easier utilization of Whisper models for audio transcription | User interface and experimental results | Humans | Haz et al., (2023) |
| Investigating hallucination phenomena in machine learning outputs | Analysis of causes and mitigation strategies for hallucination in ML models | Identification of factors leading to hallucination and proposed mitigation techniques | New insights into the nature of hallucinations in ML models | Techniques for detecting and mitigating hallucinations | Humans | Mittal et al. (2024) |
| Transcribing emergency calls to text | Use of ASR with speaker diarization; integration of Whisper and pyannote.audio models | Implementation of a system for transcribing emergency calls with high accuracy | Development of a web application for transcribing emergency calls within a secure environment | Architecture and implementation details | Humans | Thuestad, (2023) |
| Detecting fish calls using AI technologies | Application of machine learning models (not Whisper) to detect fish calls in aquatic environments | Demonstrated high accuracy of AI models in detecting specific fish calls | Advancement in the use of AI for monitoring aquatic ecosystems | Details on AI model development and deployment | Fish | Waddell et al., (2021) |
| General audio understanding model with in-context learning (ICL) | Few-shot learning via ICL and retrieval-augmented generation (RAG) | First audio understanding model demonstrating strong audio understanding ability | Introduction of Audio Flamingo with an architecture combining audio feature extraction, | Neural architecture details, including the use of ClapCap as the audio feature | Humans | Kong et al., (2024) |
|  |  |  | transformation<br>,<br>and<br>conditioning<br>on audio<br>inputs | extractor |  |  |
| Bird species<br>classification<br>using audio | Application of<br>attention<br>mechanism<br>and deep<br>learning for<br>bioacoustic<br>classification | High<br>classification<br>accuracy with<br>attention-<br>based deep<br>CNN<br>Inception-v4 | Use of multi-channel fusion<br>methods and<br>attention<br>mechanism<br>for enhanced<br>accuracy | Attention-VGG-16 model<br>and confusion<br>matrix for<br>classification | Birds | Noumida<br>and Rajan,<br>(2022) |
| Cetacean<br>communication | Use of<br>machine<br>learning for<br>analyzing<br>whale<br>vocalizations | Insights into<br>how increased<br>anthropogenic<br>sounds impact<br>whale<br>communication | Proposal for a<br>global library<br>of underwater<br>biological<br>sounds | Discussion on<br>the need for<br>interdisciplinary<br>research for<br>understanding<br>whale<br>communications | Sperm<br>Whales | Andreas et<br>al., (2021) |
| Automatic<br>speech<br>recognition<br>(ASR) | Comparison<br>of deep<br>learning<br>techniques | Detailed<br>comparison of<br>feature<br>extraction<br>methods and<br>state-of-the-art<br>classification<br>models in<br>ASR | Comprehensive<br>overview of<br>deep<br>learning's<br>impact on<br>ASR<br>performance | Discussion on<br>speech datasets<br>and online<br>toolkits | Humans | Malik et al.,<br>(2021) |
| Bat call<br>classification | Evaluation of<br>Transformer<br>network for<br>single and<br>mixed species<br>bat call<br>classification | The<br>Transformer<br>approach<br>shows better<br>performance<br>for both single<br>and mixed<br>species<br>recordings | Use of<br>Transformer<br>network for<br>bat call<br>classification<br>with fewer<br>parameters<br>and improved<br>accuracy | Comparison of<br>architectures<br>and<br>performance<br>metrics | Bats | Fundel et<br>al., (2023) |
| Decoding<br>animal<br>communication | Analyzing<br>vocalizations<br>to understand<br>communication<br>patterns | Insights into<br>syntactical and<br>temporal<br>patterns in<br>animal sounds | Novel<br>methodologies<br>for analyzing<br>animal<br>communication | Techniques for<br>deciphering<br>animal<br>vocalizations | Various | Assom<br>(2023) |
| Study of<br>sperm whale<br>vocalizations | Analysis of<br>vocal patterns<br>to understand<br>social<br>structures | Insights into<br>sperm whale<br>communication<br>and social<br>behavior | Advancement<br>in cetacean<br>bioacoustics<br>research | Techniques for<br>analyzing and<br>categorizing<br>whale<br>vocalizations | Sperm<br>Whales | Kather<br>(2023) |
| Linking sound<br>perception to<br>semantic<br>understanding | Comparative<br>study of<br>behavioral and<br>neural<br>responses to<br>sounds | Understanding<br>of how the<br>brain<br>processes and<br>interprets<br>sounds | Insights into<br>the neural<br>basis of sound<br>perception and<br>its semantic<br>interpretation | Analysis of<br>brain activity<br>in response to<br>sound stimuli | Humans | Huang et al.,<br>(2024) |

## 4. Objectives

The primary objective of this study is to apply the Whisper NLP model to analyze chicken vocalizations, with the goal of improving welfare monitoring in poultry farming. By using a tool initially created for human language, we aim to explore whether the complexities of chicken vocalizations can be effectively interpreted to provide meaningful insights into their well-being.

The hypotheses of the current study are,

1. The Whisper NLP model can be successfully adapted to classify chicken vocalizations into meaningful categories related to welfare conditions.
2. The sentiment analysis through the Whisper NLP model will reveal distinct patterns corresponding to the stress levels and health status of chickens.
3. The transcription and sentiment analysis of chicken vocalizations will provide a more accurate and objective assessment of welfare compared to traditional observation methods.

This study seeks to bridge the gap between advanced computational models and practical animal welfare assessment, with the potential to transform current practices in poultry farming. By validating these hypotheses, the research could lay the groundwork for broader applications of NLP technology in animal welfare science.

## 5. Materials and Methods

This study employed three distinct datasets to investigate the semantics of avian (poultry chicken) communications through sentiment analysis. The overarching goal was to discern patterns that might indicate stress or distress in poultry vocalizations, leveraging advanced machine learning techniques and audio processing models.

### 5.1. Data Collection

Dataset 1 was curated from our research group’s prior work and partially presented in Neethirajan, 2023. It encompasses audio recordings from the CARUS animal experimental facility at Wageningen University, Netherlands featuring fifty-two Super Nick chickens across three cages under varying conditions to simulate commercial poultry environments. Stress was induced using an umbrella and loud dog barking sounds to elicit flight responses, with recordings taken before and after these events to capture the range of vocalizations. The dataset is accessible at Zenodo.

Dataset 2, as detailed in Adebayo et al., 2023, comprises 346 audio signal files collected at the poultry research farm at Bowen University, Nigeria. These files are systematically categorized into three folders based on the health status of the birds and environmental conditions: healthy, noisy, and unhealthy. Each category is designed to reflect distinct aspects of the birds’ welfare and the ambient environment. The healthy folder contains 139 files featuring typical vocalizations of chickens under normal health conditions. The noisy folder includes 86 files with background noises such as moving vehicles and human conversations, in addition to sounds related to poultry behaviors like feeding and pecking. The unhealthy folder encompasses 121 files, selectively comprising sounds associated with potential health issues, such as chicken coughs, snores, and rales. The duration of the audio files varies from 5 to 60 seconds, offering a diverse range of sound environments and vocalizations for analysis. The methodology used for recording and categorizing these vocalizations is comprehensively described in the publication. This dataset is accessible for download at Mendeley Data, facilitating further research and analysis.

### 5.2. Data Preparation and Analysis Pipeline

The analysis pipeline began with the installation of critical software and libraries, including FFmpeg for audio file processing and Python libraries such as Librosa, NumPy, Pandas, and Scikit-learn for data manipulation, feature extraction, and machine learning tasks. Audio Pre-processing involved normalization, noise reduction, and segmentation to prepare the files for further analysis. The Whisper model from OpenAI, alongside custom scripts in Python, facilitated the transcription of audio signals into text, which were subsequently analyzed for sentiment using the Natural Language Toolkit (NLTK).

### 5.3. Feature Representation and Machine Learning Models

N-gram Models and Count Vectorization were employed to transform the transcribed text data into numerical features suitable for machine learning analysis. N-grams of varying lengths (unigrams, bigrams, trigrams) were generated and transformed using count vectorization to create a frequency matrix of these n-grams for each vocalization.

The processed data then served as input for Machine Learning Models, including Naive Bayes, Random Forest, and Logistic Regression classifiers, to categorize the vocalizations into predefined categories (healthy, noise, unhealthy). This classification aimed to identify patterns correlating with the birds’ health status and environmental conditions.

### 5.4. Sentiment Analysis

In our analysis, the SentimentIntensityAnalyzer from the NLTK library played a pivotal role, analyzing the emotional content of the vocalizations transcribed from audio to text. This tool provided sentiment scores, which were then classified into positive, negative, or neutral categories based on polarity scores, offering a clear measure of the emotional undertones present in the data.

We employed a structured approach to organize these sentiment scores into an Excel sheet, enabling an effective comparison and visualization of sentiment variations across the studied datasets and conditions. This methodical compilation aids in understanding the emotional impacts reflected in poultry vocalizations under various scenarios.

### 5.5. Ethical Considerations

All procedures involving animal subjects were conducted in strict accordance with ethical guidelines and approved by the appropriate institutional review boards. Efforts were made to minimize stress and discomfort to the animals throughout the study.

## 6. Results and Discussions

### 6.1. Results from Dataset 1

#### 6.1.1. Examination of Character Occurrences

The analysis of character frequency across different texts (Figure 1) offers a nuanced perspective on the linguistic composition inherent within our datasets. Comprising 39 distinct entries, this overview encapsulates both the ubiquity and scarcity of specific characters. Predominantly, the character ‘i’ emerges as the most prevalent, recorded 68,387 times, closely followed by ‘a’ and ‘o’, with 30,359 and 45,021 occurrences, respectively. This analysis extends beyond the realm of common English alphabets to encompass punctuation marks (’.’, ‘,’, ‘-’) and lesser-seen characters (’æ’, ‘′’), alongside numerals (’1’, ‘2’, ‘3’, ‘4’, ‘5’). Intriguingly, the presence of non-English characters (’諷’, ‘•’, ‘怎’) hints at either the dataset’s multilingual nature or the inclusion of data anomalies. Certain characters, such as ‘j’ and ‘y’, manifest less frequently, offering a window into the character distribution’s diversity and aiding in further linguistic analysis.

**Figure 1.**
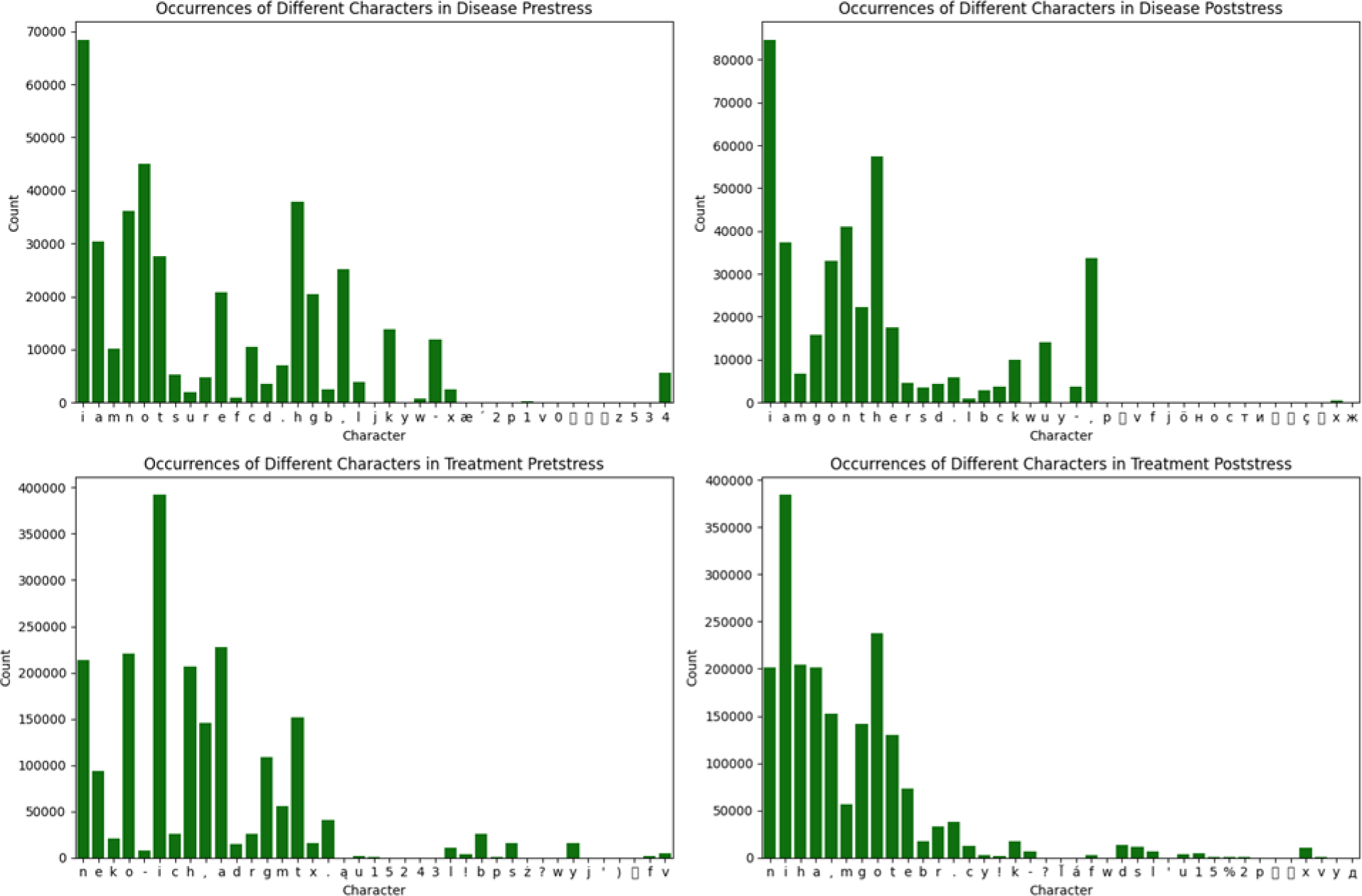
Character Frequency Analysis Across Four Phases – (A) Disease Prestress, (B) Disease Poststress, (C)Treatment Prestress, and (D) Treatment Poststress.

This intricate character mapping extends to the disease post-stress dataset, wherein ‘i’ notably dominates once again, with 84,597 appearances, underscoring its linguistic significance. This dataset also documents the inclusion of punctuation and numerals, alongside less common and non-English characters (’시’, ‘ö’, ‘н’, ‘아’), suggesting a linguistic breadth that spans beyond monolingual confines. Such character diversity is crucial for understanding the linguistic intricacies and potential multilingual dimensions embedded within the dataset.

In examining the treatment prestress dataset, ‘n’ notably prevails with 213,382 occurrences, closely followed by ‘e’ and ‘o’. The incorporation of punctuation and numerals alongside infrequent non-English characters (’ą’, ‘ż’, ‘た’) indicates a rich linguistic diversity and possibly specific terminologies pertinent to the treatment prestress context. This character assortment, including both common and rare occurrences, reflects the text’s linguistic diversity and complexity, enriching the analysis of treatment-related discourse.

Similarly, the treatment post-stress dataset reveals ‘i’ as the most recurring character, with ‘n’ and ‘h’ also prominently featured. The dataset is characterized by a broad array of English characters, punctuation marks, and numerals, suggesting a comprehensive linguistic composition. Noteworthy is the inclusion of non-English characters (’l’, ‘á’), indicating a multilingual aspect or specific terminologies relevant to the treatment post-stress phase. This diverse character usage underscores the complexity and richness of the linguistic content, offering insights into the textual elements within the dataset.

Overall, the character frequency analysis across these datasets illuminates the linguistic richness and diversity present within the texts. It highlights the prevalence of certain characters, the incorporation of punctuation and numerals, and the intriguing presence of non-English characters, suggesting a multilayered linguistic composition. This analysis not only contributes to our understanding of the datasets’ linguistic properties but also facilitates a deeper exploration of the texts’ linguistic and multilingual nuances.

#### 6.1.2. Analytical Insights into Textual Composition: Evaluating Linguistic Metrics Across Different Stress Phases

The analysis of text metrics (Figure 2) illuminates the structural and linguistic attributes of the disease prestress dataset. The “text_length” measure, quantifying at 307,281, marks the aggregate character count of the text, providing a fundamental perspective on its volumetric composition. The “num_of_vowels” metric, tabulated at 120,258, enumerates the vowels within the text, highlighting their pivotal role in the phonetic framework and articulation nuances. Equally, the “num_of_consonants” figure, standing at 133,753, catalogs the consonants, underscoring their indispensable function in crafting linguistic sounds and patterns, thereby enriching the text’ phonological and articulatory nuances. The “num_of_stopwords” count, recorded at 8,687, identifies the frequency of grammatically essential yet semantically sparse words, revealing insights into the syntactic organization, linguistic practices, and stylistic nuances of the text. This evaluation of stopwords aids in the dissection of textual content, fostering sentiment analysis and thematic extraction, thus advancing the scope of text analysis and comprehension of natural language.

**Figure 2.**
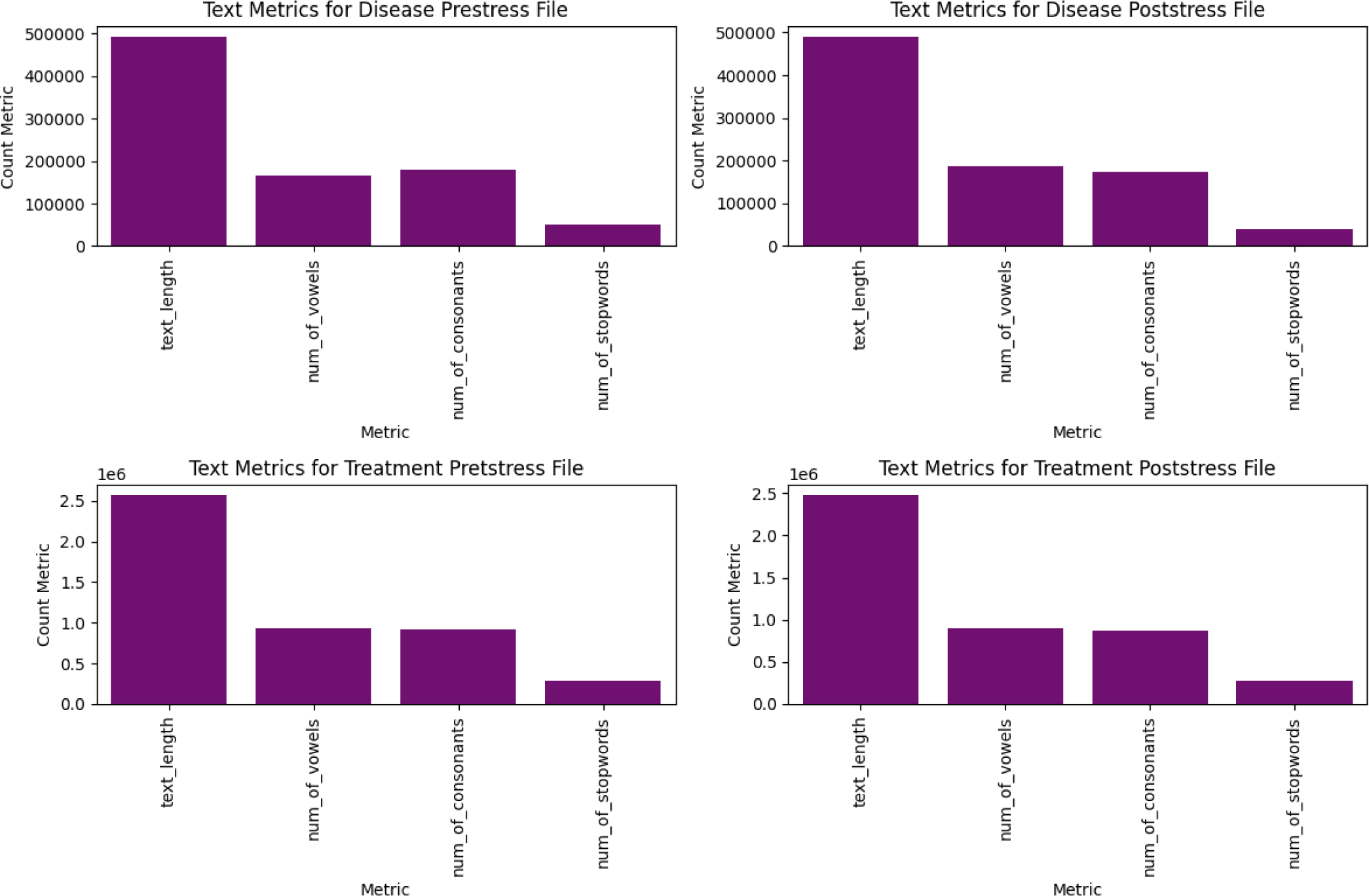
Comparative Analysis of Text Metrics: Text Length, Number of Vowels, Number of Consonants, and Number of Stopwords Across Disease Prestress, Disease Poststress, Treatment Prestress, and Treatment Poststress.

Similarly, the post-stress dataset’s text metrics reveal distinctive characteristics. The “text_length” at 315,039, “num_of_vowels” at 137,436, and “num_of_consonants” at 124,032, delineate the textual landscape, emphasizing the critical roles of vowels and consonants in the phonetic and phonological construct. The “num_of_stopwords” metric, at 7,115, further dissects the textual structure, underscoring the prevalence of functional yet minimally meaningful words, which enriches the analytical framework for content classification, sentiment examination, and thematic delineation, thereby enhancing linguistic and narrative understanding.

In examining the treatment prestress dataset, the metrics such as “text_length” at 1,598,079, “num_of_vowels” at 674,507, and “num_of_consonants” at 652,825, shed light on the voluminous and intricate linguistic architecture. These metrics not only chart the text’ dimensional and phonetic landscape but also accentuate the stylistic and syntactical fabric through the “num_of_stopwords” metric at 47,082. This granular analysis facilitates a deeper exploration of textual substance, supporting advanced interpretative tasks such as thematic identification, sentiment analysis, and linguistic profiling.

For the treatment post-stress, metrics like “text_length” at 1,487,076, “num_of_vowels” at 642,086, and “num_of_consonants” at 595,772, offer a comprehensive overview of the text’ scale and phonological complexity. The “num_of_stopwords” at 30,139 further elucidates the textual syntax and stylistic attributes, enriching the analytical capacity for textual categorization, emotional analysis, and thematic investigation, thereby broadening the avenues for textual understanding and natural language analysis.

#### 6.1.3. Phonetic Composition Analysis - Dissecting the Roles of Vowels and Consonants Across Different Stages of Stress and Treatment

By dissecting the roles of the phonetic components, we can gain invaluable insights into the linguistic underpinnings that define each stage, offering a deeper understanding of the textual dynamics and the potential implications for further linguistic and thematic exploration.

##### 6.1.3.1. Vowel Distribution and Impact

Our analysis (Figure 3) uncovers a significant variation in vowel usage across the datasets, reflecting the nuanced phonetic structures that underpin the linguistic texture of each stage. The character ‘i’, consistently the most frequent vowel across all stages, highlights its fundamental role in contributing to the phonetic diversity and complexity of the text. Notably, in the treatment post-stress phase, ‘i’ ascends to a staggering 384,029 occurrences, underscoring its prominence and the phonetic richness it brings to the text. The analysis further reveals a compelling dynamic in the frequency of ‘a’ and ‘e’, where ‘a’ exhibits a marked increase in prevalence in the treatment prestress dataset, indicating a shift in linguistic patterns that could correlate with specific thematic focuses or stylistic changes in the text. Conversely, ‘e’ maintains a relatively stable presence, reinforcing its essential role in the phonetic composition of the corpus.

**Figure 3.**
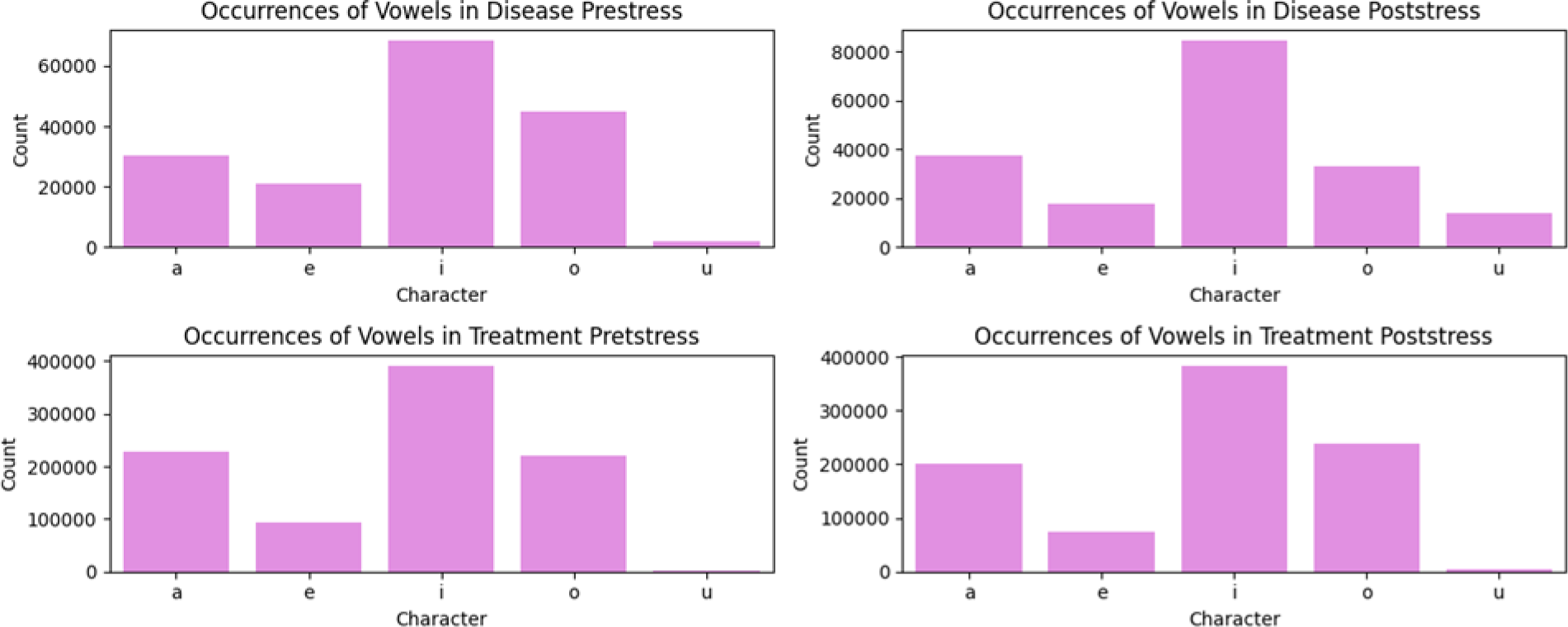
Distribution of Vowel Frequencies Across Different Stages of Stress and Treatment.

##### 6.1.3.2. Consonant Dynamics and Linguistic Texture

Turning to consonants (Figure 4), the analysis illuminates the foundational roles of ‘n’, ‘h’, and ‘g’ in sculpting the corpus’s phonological properties. The pronounced increase in the occurrence of ‘n’ in the treatment prestress phase, reaching 213,382 instances, suggests a linguistic adaptation possibly tied to the articulation of specific concepts or terminologies inherent to this stage. Moreover, the consonant ‘h’ demonstrates a substantial presence, particularly in the treatment post-stress phase, with 204,586 occurrences, indicating its significance in enhancing the text’ phonological depth.

**Figure 4.**
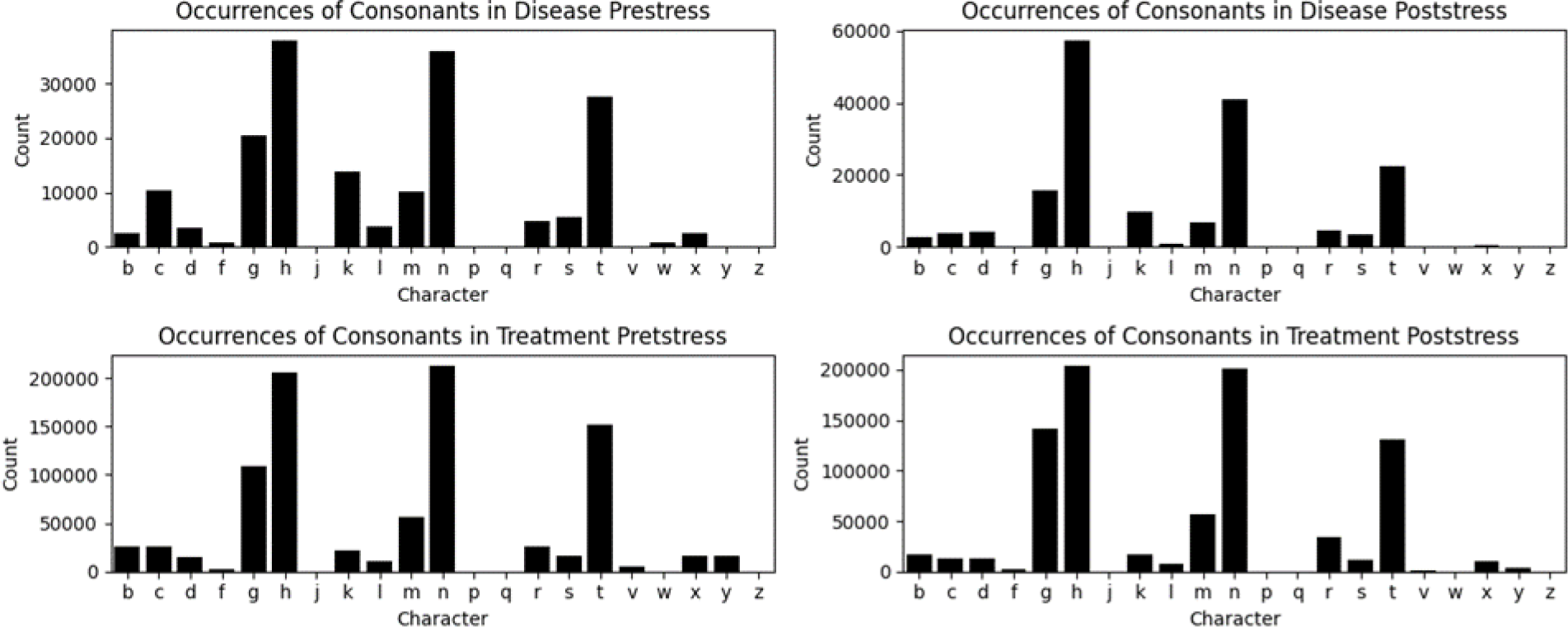
Consonant Frequency Analysis in Various Phases of Stress and Treatment.

**Figure 5A.**
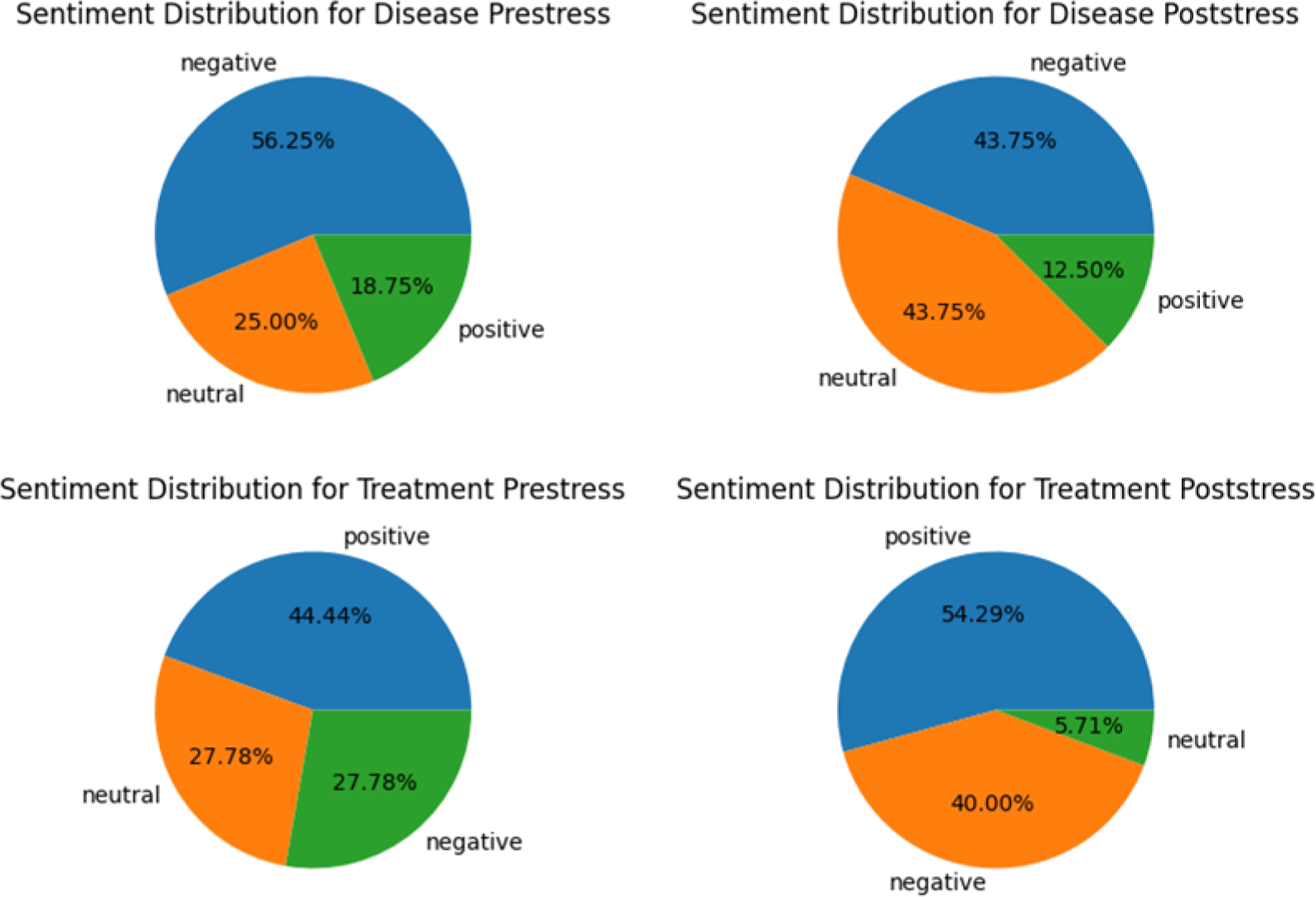
Comparative Sentiment Distribution Across Disease and Treatment Phases

**Figure 5B.**
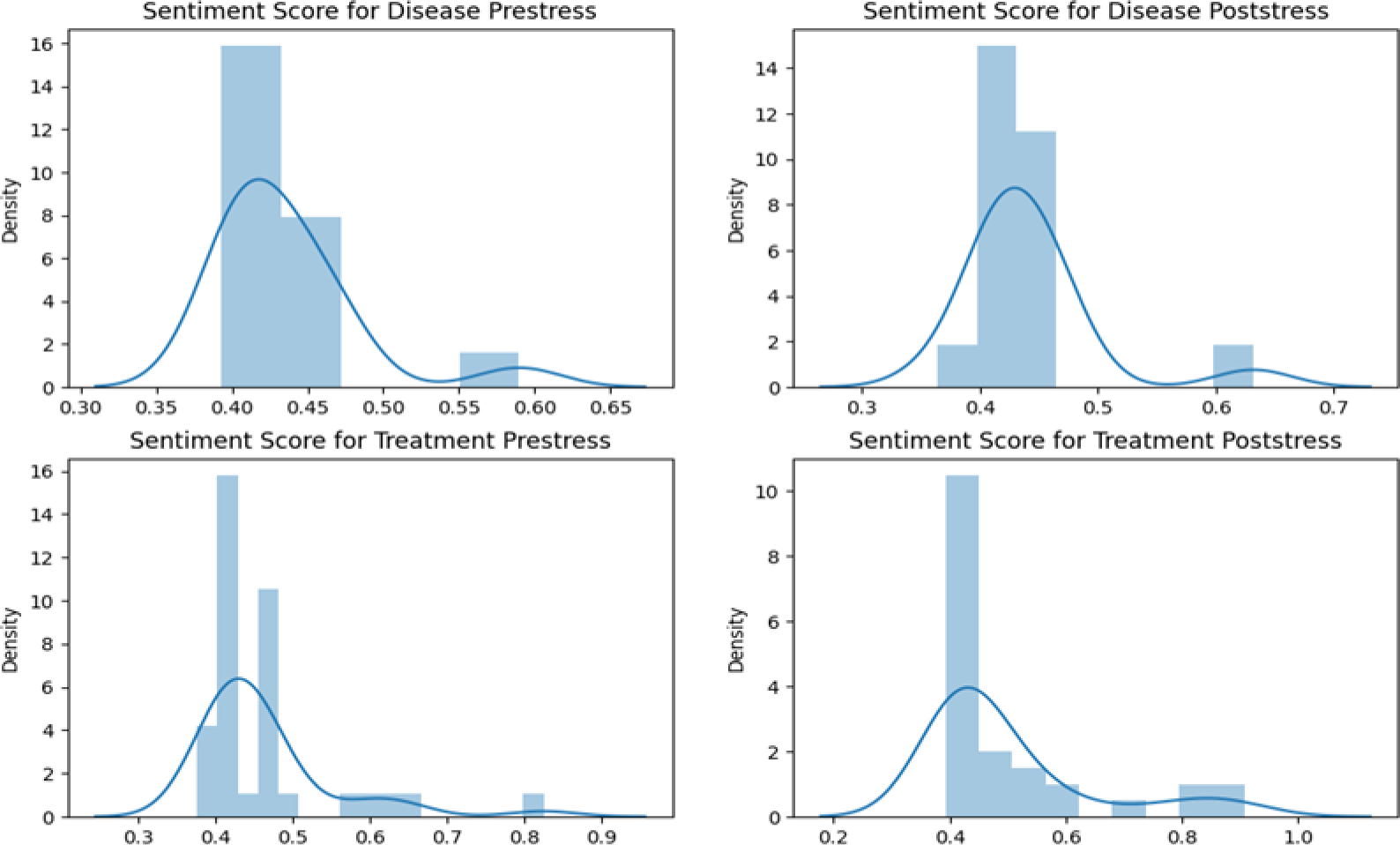
Distribution of Sentiment Scores Across Disease and Treatment Phases - An Analysis Using Displots for Disease Prestress, Disease Poststress, Treatment Prestress, and Treatment Poststress

**Figure 5C.**
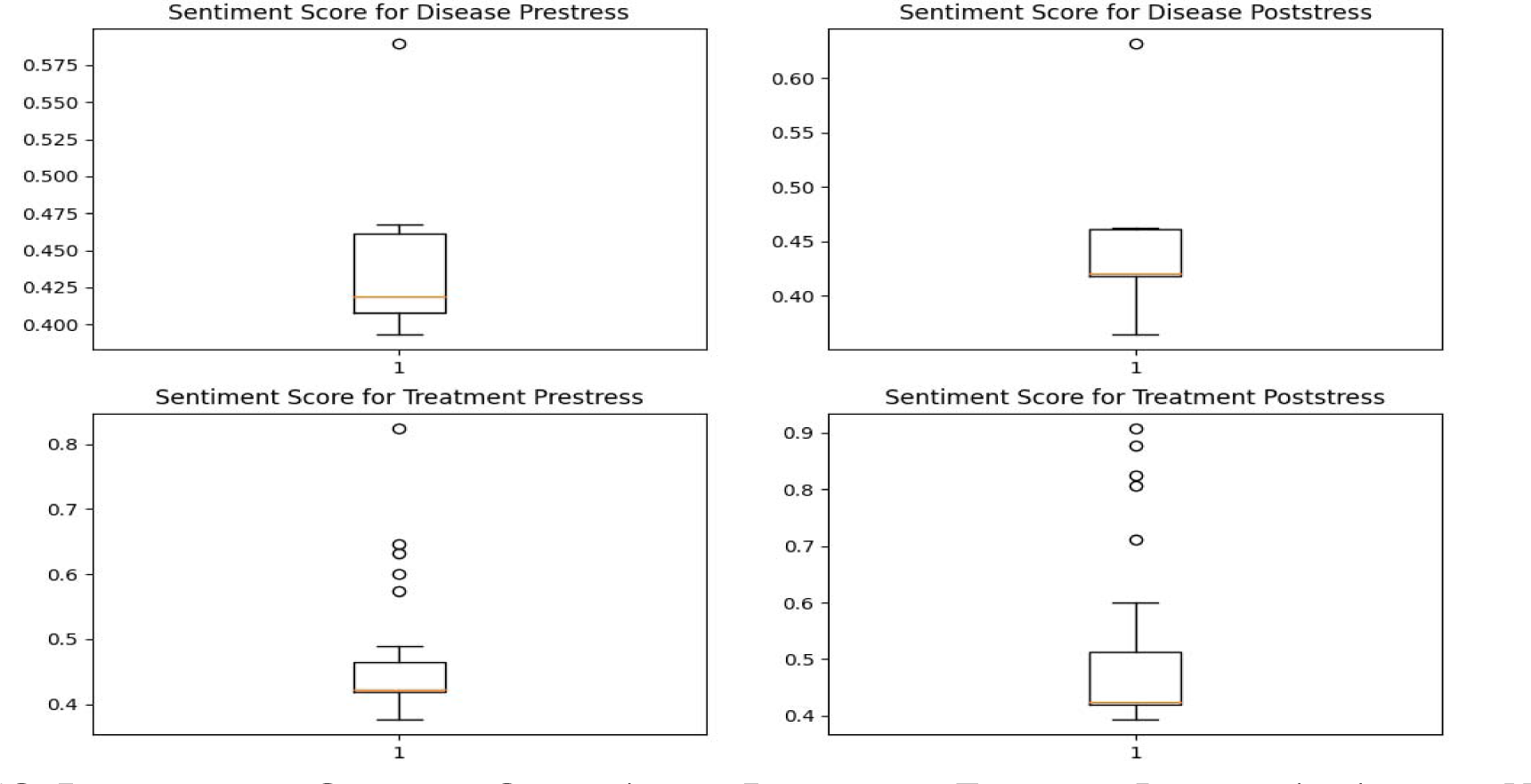
Distribution of Sentiment Scores Across Disease and Treatment Phases - An Analysis Using Boxplots for Disease Prestress, Disease Poststress, Treatment Prestress, and Treatment Poststress

**Figure 5D.**
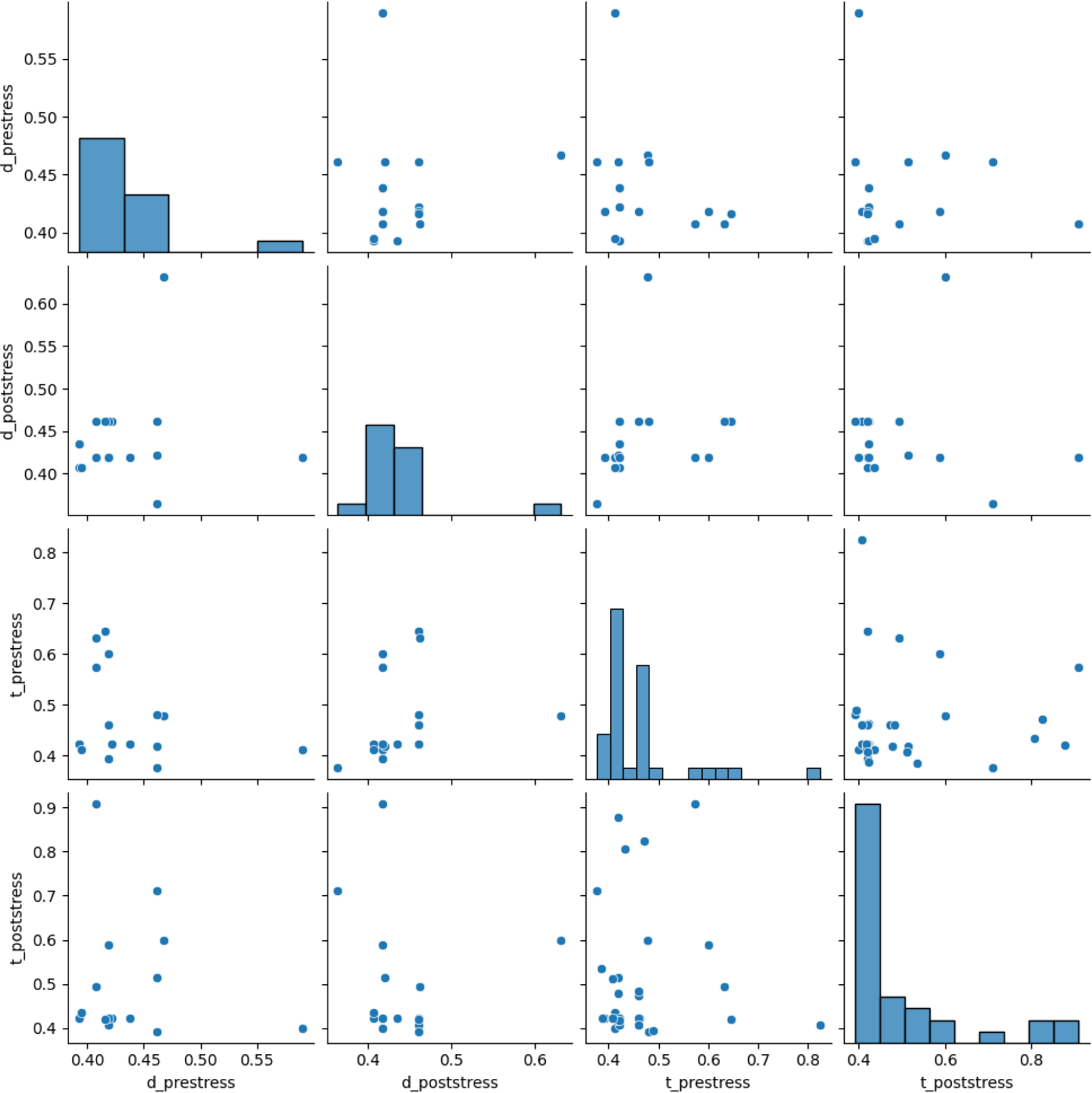
Analysis of Sentiment Correlation Among Vocalizations of Laying Hens Across Different Stress Phases. This graph illustrates the lack of correlation between sentiment scores across four distinct phases: disease prestress (d_prestress), disease poststress (d_poststress), treatment prestress (t_prestress), and treatment poststress (t_poststress). The Y-axis represents sentiment scores in the treatment poststress phase, while the X-axis categorizes the sentiment scores from the other three phases, showcasing a random distribution of data points without discernible patterns or trends.

Furthermore, the consonant ‘g’ emerges as a key player in the treatment prestress dataset, with its frequency soaring to 77,893 occurrences. This suggests a pivotal role in conveying specific phonetic nuances or contributing to the overall articulatory characteristics of the text, potentially tied to the expression of complex concepts or terminologies relevant to the treatment phase.

##### 6.1.3.3. Phonetic Richness and Linguistic Diversity

The analysis also highlights the occurrence of less common vowels and consonants, alongside instances of non-English characters, reflecting the corpus’s linguistic diversity and the incorporation of multilingual elements or specialized terminologies. Such phonetic richness not only contributes to the corpus’s complexity but also hints at the underlying thematic variances and contextual specificities of each dataset.

#### 6.1.4. Emotional Dynamics Across Disease and Treatment Phases - A Sentiment Analysis

The analysis of sentiment distribution by category across different stages related to disease and treatment reveals intricate emotional landscapes. During the disease prestress phase, a predominance of negative sentiment, exceeding half of the analyzed discourse, underscores a widespread atmosphere of apprehension and concern. This phase is marked by a notable prevalence of anxiety, fear, and worry, reflecting the psychological impact before any medical intervention. Approximately one-fourth of the sentiment distribution is characterized as neutral, signifying a substantial segment of the dialogue that does not distinctly align with either positive or negative emotions. Positive sentiments are markedly less frequent, comprising less than a quarter of the discourse, suggesting that instances of optimism or hope are significantly outweighed by negative emotions.

Upon transitioning to the poststress phase of the disease, a notable shift in sentiment distribution is observed. Negative and neutral sentiments approach parity, illustrating a nuanced equilibrium between ongoing concerns and a newfound sense of acceptance or stability following intervention. This phase is characterized by the persistence of negative sentiments, highlighting enduring challenges or adverse outcomes associated with the disease, despite the implementation of treatment strategies. Concurrently, the pronounced representation of neutral sentiments may reflect a pragmatic or resigned perspective among individuals. Positive sentiments remain minimal, indicating cautious optimism or restrained acknowledgment of positive developments amid prevailing difficulties.

In the treatment prestress phase, the sentiment landscape is distinctly optimistic. Positive sentiments predominate, reflecting widespread anticipation and confidence in the efficacy of forthcoming treatment options. Negative and neutral sentiments are equally represented, suggesting a balanced acknowledgment of potential uncertainties and the optimistic outlook towards treatment outcomes. This phase emphasizes the collective focus on the anticipated benefits and success of the proposed interventions, relegating concerns and uncertainties to the background.

The treatment poststress phase marks a significant evolution towards positivity. Positive sentiments constitute the majority, indicating the overall perceived success or effectiveness of the treatment in ameliorating the condition. The presence of negative sentiments diminishes substantially from the prestress phase, denoting a decrease in adverse concerns or outcomes following treatment. Neutral sentiments become scarce, underscoring a dialogue dominated by clear emotional valences, either positive or negative, with minimal expressions of ambivalence or neutrality.

#### 6.1.5. Analyzing Sentiment Dynamics Across Disease and Treatment Phases - A Comparative Study Using Displots and Boxplots

In the Disease Prestress phase, sentiment scores predominantly oscillate between 0.35 and 0.60, with a significant concentration in the range of 0.39 to 0.47. This distribution suggests a moderate sentiment intensity, with neither pronounced positivity nor negativity dominating the discourse. Similarly, the Disease Poststress phase exhibits a comparable sentiment score range from 0.35 to 0.65, with a central clustering around the 0.5 mark, indicating a prevailing neutrality or moderate sentiment intensity. As we progress to the Treatment Prestress phase, the sentiment scores extend from 0.35 to 0.85, yet the bulk of the scores remains around the moderate median of 0.5. Contrastingly, the Treatment Poststress phase reveals a shift with scores ranging from 0.40 to 1.0 and a notable concentration of scores between 0.4 and 0.6, suggesting an elevation in sentiment intensity and a skew towards more positive emotional expressions.

The boxplot analysis further delineates the sentiment score dynamics, presenting a detailed view of their distribution. In the Disease Prestress phase, the scores span from approximately 0.38 to 0.474, with outliers indicating extreme sentiments. A slight shift is observed in the Disease Poststress phase, with a narrowing of the interquartile range and an outlier marking a significant deviation. The Treatment Prestress phase mirrors this pattern, albeit with a notable outlier indicating a departure from the median sentiment score. The Treatment Poststress phase, however, showcases a broader range of sentiment scores with multiple outliers, highlighting a diversification in emotional responses and suggesting a more pronounced positive sentiment post-treatment.

Both the Dispolot and Boxplot analyses collectively showcase the nuanced sentiment landscapes associated with each phase of disease and treatments in the laying hens. They reveal a trajectory from moderate and balanced emotional expressions in the prestress phases towards more pronounced positivity in the post-treatment phase, reflecting the potential emotional impacts of stress in the chicken birds.

#### 6.1.6. An Emotion Score Correlation Analysis

The investigation into the correlation among emotion scores derived from laying hen vocalization datasets unveils a notable absence of discernible correlations. Analyzing the emotion correlation graph, a random distribution of data points is observed, devoid of any patterns or trends that might suggest a systematic relationship. In instances where emotion scores exhibit correlation, one would anticipate a structured arrangement within the scatter plot, characterized by data points aligning along a trend line or aggregating into distinct clusters. Contrarily, the data points in this study are dispersed randomly across the graph, signaling no evident connection between the emotion scores across different files. This observation implies that the emotional content captured within each dataset stands autonomously, uninfluenced by and not influencing the emotions documented in other datasets. The lack of correlation underscores the diversity and heterogeneity present in the emotional expressions of the vocalizations analyzed, suggesting each dataset embodies a distinct emotional landscape. Consequently, this analysis conclusively determines the absence of a recognizable correlation among the emotion scores, with the scatter plot elucidating a pattern of randomness as opposed to identifiable systematic relationships. This insight accentuates the nuanced and varied nature of emotional expressions among laying hens, highlighting the complexity of their emotional communication.

#### 6.1.7. Emotional Trajectories in Avian Vocalizations

##### 6.1.7.1. Emotional Intensity and Distribution Across Treatment Phases

The emotional landscape of laying hens, as inferred from their vocalizations across different stress and treatment phases, presents a nuanced picture of their psychological states. Initially, during the Disease Prestress phase, emotional expressions predominantly hover within the neutral range, suggesting an equilibrium in emotional states prior to any specific interventions. This equilibrium shifts subtly (Figure 6A) in the Disease Poststress phase, where a slight increase in expressions of sadness becomes evident, hinting at the nuanced impact of disease and subsequent interventions on avian emotional states.

**Figure 6A.**
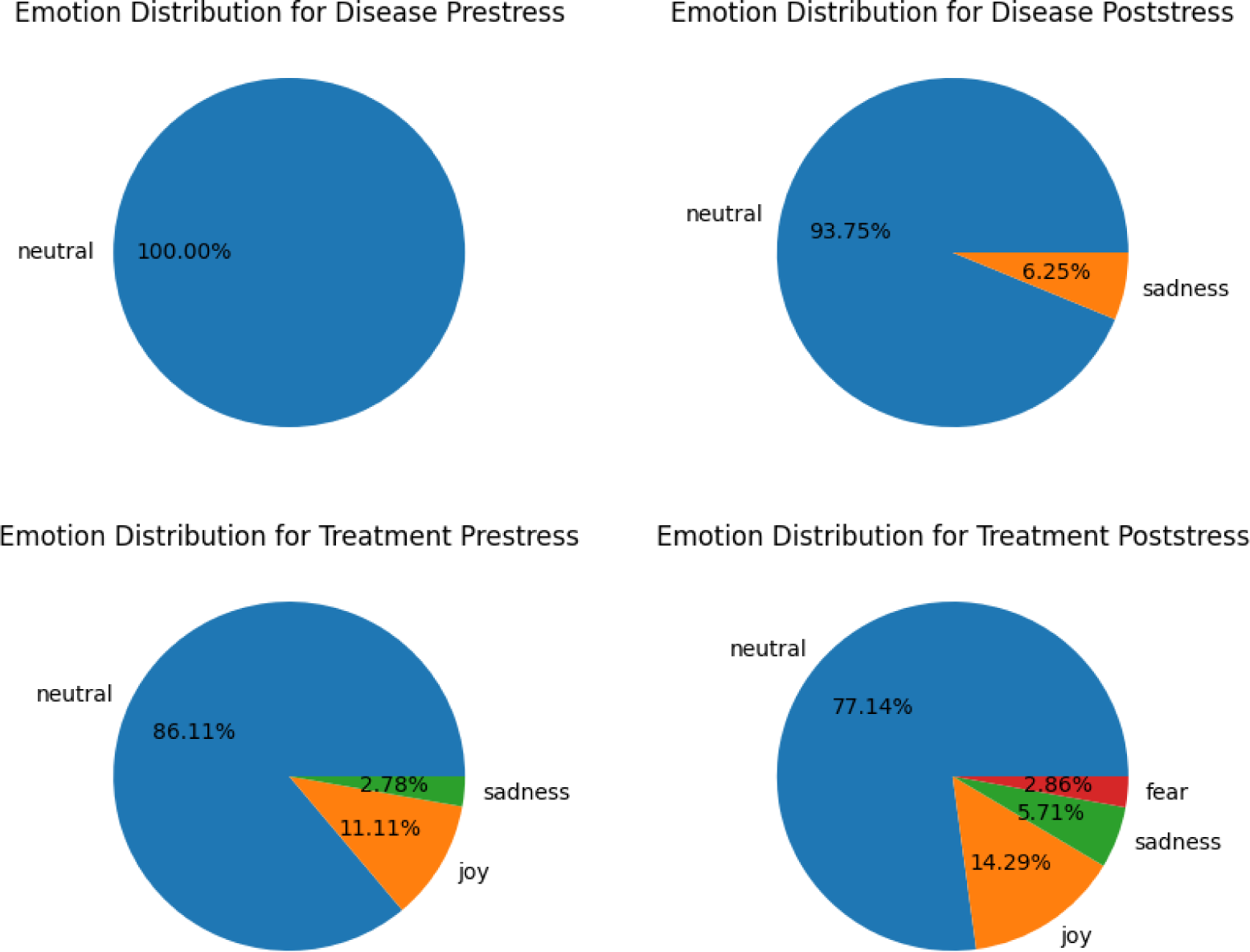
Emotional Landscape of Laying Hens. A Comparative Pie Chart Analysis across Phases. This series of pie charts illustrates the distribution of emotions—neutral, fear, sadness, and joy—in laying hens during disease prestress, poststress, and pre and post treatment phases, providing a visual representation of emotional shifts and the prevalence of each emotion in response to disease and treatment.

**Figure 6B.**
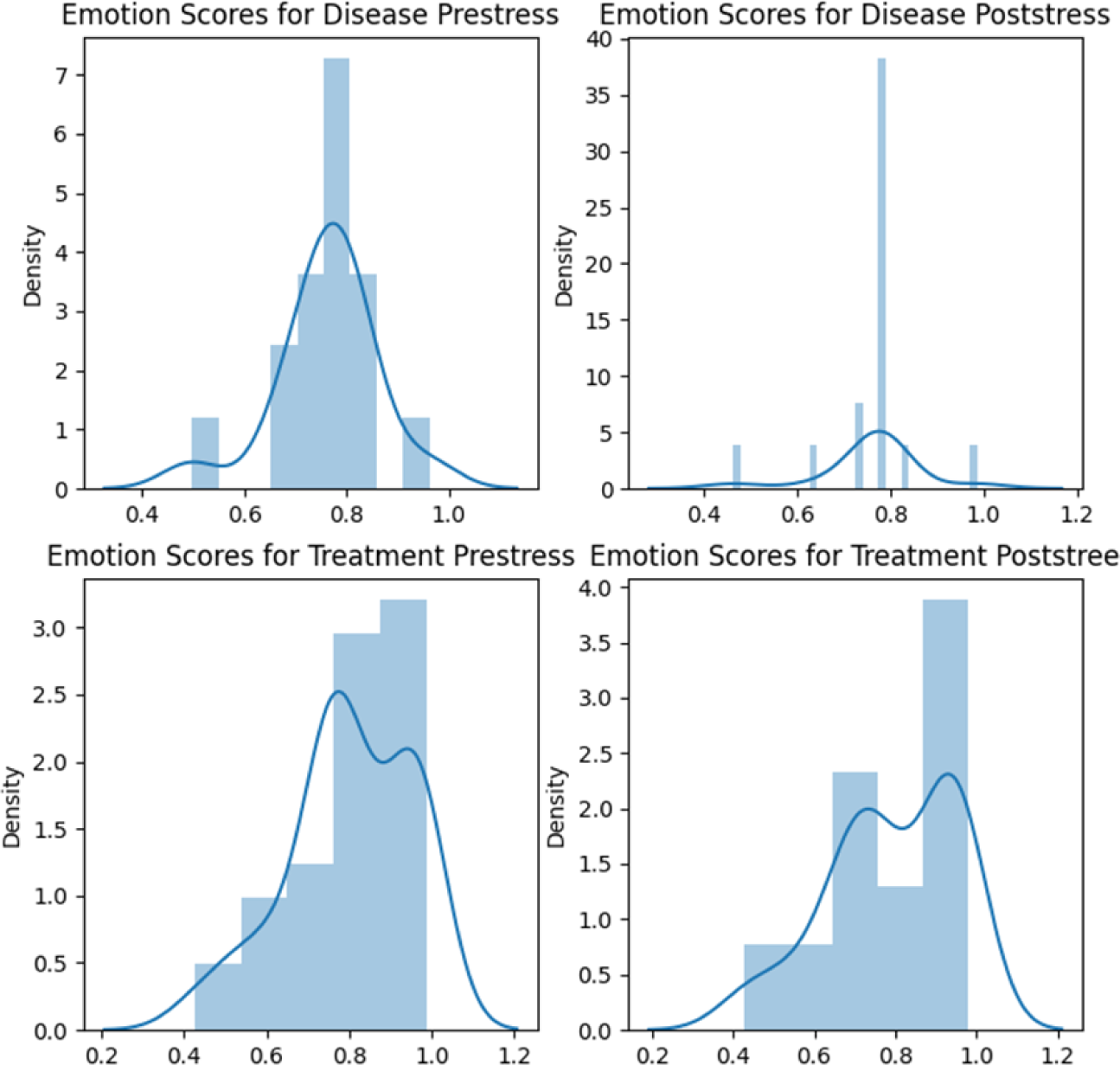
Density Distribution of Emotion Scores Across Treatment Phases in Laying Hens. This displot showcases the frequency distribution of emotion scores, ranging from 0.5 to 1.0, during disease prestress, disease poststress, treatment prestress, and treatment poststress phases, emphasizing the density and spread of emotional intensities experienced by laying hens.

**Figure 6C.**
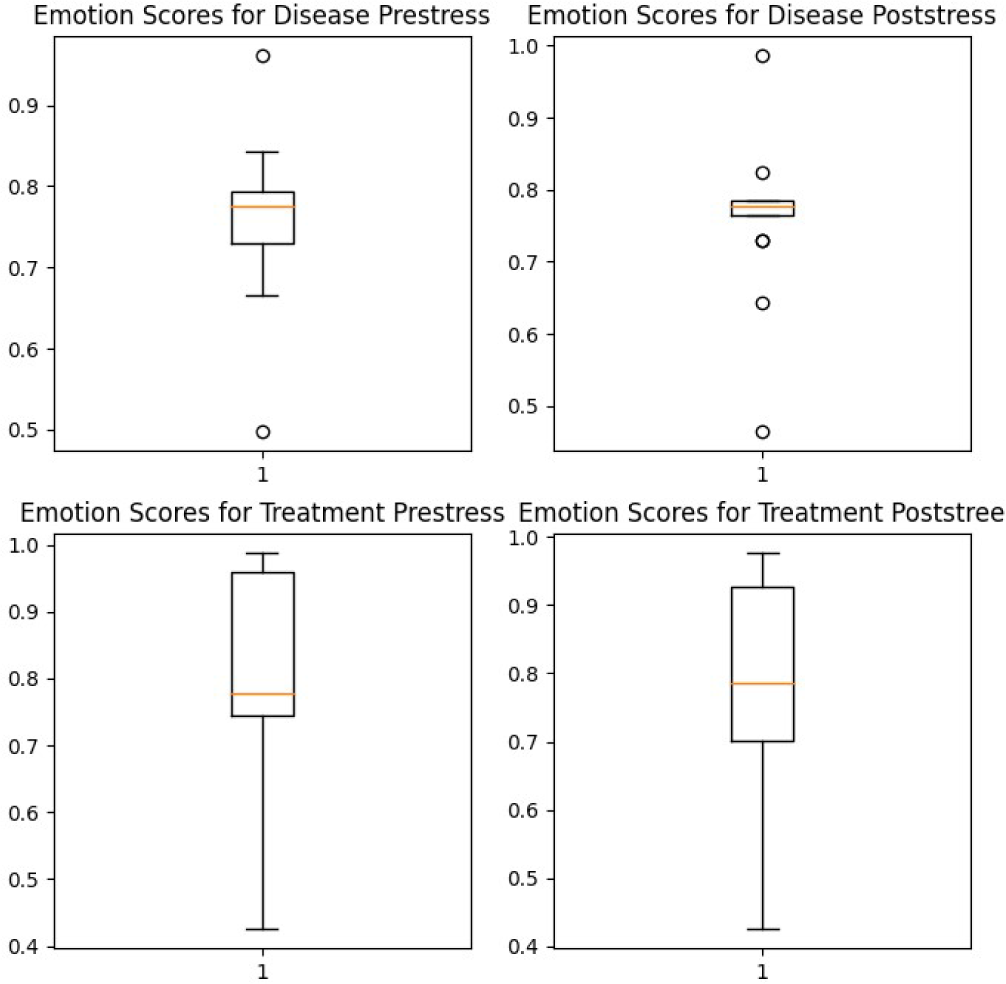
Boxplot Visualization of Emotion Scores in Laying Hens Across Treatment Phases. This box plot displays the range and distribution of emotion scores from 0.5 to 1.0 for laying hens during disease prestress, disease poststress, treatment prestress, and treatment poststress phases, highlighting median values, interquartile ranges, and outliers in emotional expression intensity.

**Figure 6D.**
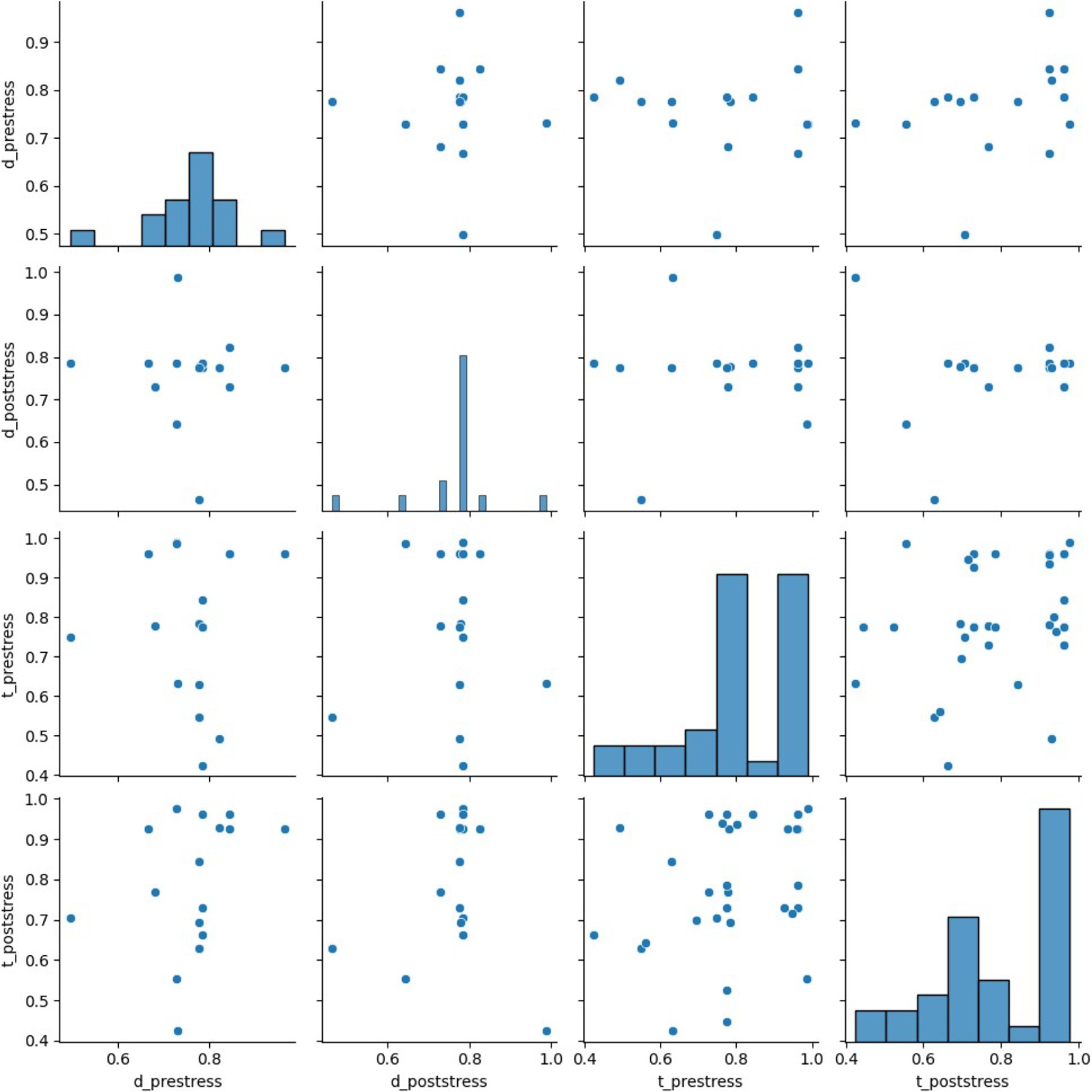
Scatter Plot Analysis of Emotion Score Correlation Among Treatment Phases for Laying Hens. This scatter plot examines the correlation, or lack thereof, among emotion scores across disease prestress, disease poststress, treatment prestress, and treatment poststress phases, depicting a non-linear distribution that suggests individual variability in emotional response patterns among laying hens.

As we transition into the Treatment Prestress phase, there is a noticeable predominance of neutral emotions, interspersed with instances of joy and sadness, reflecting the anticipatory and varied emotional responses to impending treatment. Interestingly, the post-treatment phase unveils a richer tapestry of emotional expressions, with increased instances of joy amidst continuing neutrality, sadness, and fear. This diversification post-treatment suggests a complex emotional response to the outcomes of medical interventions, illuminating the psychological rebound and adjustments experienced by the birds.

##### 6.1.7.2. Dissection of Emotion Scores - Boxplot and Displot Insights

The quantitative analysis of emotion scores through boxplot and displot methodologies further elucidates the emotional responses of laying hens. The boxplot analysis across phases reveals a significant span of emotional intensities, with notable outliers indicating instances of extreme emotional states. This variability underscores the depth of emotional experiences and potential stressors impacting avian welfare.

Similarly, displot analysis showcases a strong clustering of emotion scores towards higher intensities, particularly in the post-treatment phase, indicating a pronounced emotional response to the interventions undertaken. These findings suggest that while treatment may ameliorate physical symptoms, the emotional ramifications warrant careful consideration, emphasizing the need for holistic approaches to avian healthcare that address both physical and psychological well-being.

##### 6.1.7.3. Emotion Score Correlation - A Scatter Plot Analysis

The investigation into the correlation of emotion scores among the vocalizations across phases reveals a striking absence of systematic patterns. The scatter plot analysis indicates a random distribution of data points, signifying a lack of correlation between the emotional expressions across different phases. This randomness highlights the individualized nature of emotional responses among laying hens, challenging the notion of uniform emotional reactions to disease and treatment. The heterogeneity observed underscores the complexity of avian emotional states, suggesting that each bird may experience and express emotions in a uniquely personal manner.

#### 6.1.8. Textual Analysis of Data Similarity Across Treatment and Disease Phases

The analytical exploration of data files using the cosine similarity metric has unveiled a pronounced degree of concordance amongst the datasets across various disease and treatment stages. This high similarity, consistently exceeding 90%, signifies uniform data patterns, elucidating a seamless narrative of health-related experiences and interventions.

In the phase of Disease Prestress, an impressive similarity index of over 90% is observed when compared to Disease Poststress, Treatment Prestress, and Treatment Poststress phases. This remarkable congruence underscores a unified data pattern that transcends the temporal phases of disease progression and intervention, showcasing a persistent thread of health-related phenomena.

The Disease Poststress phase, likewise, displays an eminent similarity exceeding 90% with the remaining datasets, notably achieving a 94% similarity with both the Treatment Prestress and Treatment Poststress phases. This accentuates the data’s uniformity, highlighting a cohesive pattern that bridges the pre- and post-treatment experiences.

Transitioning to the Treatment Prestress phase, a conspicuous similarity of more than 90% is evident with all other datasets, particularly exhibiting a 95% similarity with the Treatment Poststress phase and a 94% similarity with the Disease Poststress phase. This underscores the intrinsic coherence among the datasets, reflecting a consistent pattern that navigates through the nuances of treatment preparation and aftermath.

Similarly, the Treatment Poststress phase maintains this trend of high similarity, with over 90% correlation observed with all datasets, especially noting a 95% similarity with the Treatment Prestress phase and a 94% similarity with the Disease Poststress phase. This reaffirms the datasets’ uniformity, further emphasizing the integrated narrative of disease and treatment progression. The pronounced similarity across all datasets highlights a remarkable continuity and coherence in the patterns of data, reflective of the interconnected nature of disease and treatment phases.

#### 6.1.9. Lexical Analysis of Vocalization Data Across Different Phases of Chicken Birds’ Stress and Treatment

##### 6.1.9.1. Word Frequency Analysis in Disease and Treatment Contexts - Disease Prestress Phase - Lexical Insights

The analysis of word frequencies within the disease prestress phase reveals predominant terms such as “nihai” with 17,241 mentions, signifying a possible keyword in the dataset. Other significant terms include “going,” “kichi,” and “neko,” suggesting themes of action and possibly social interactions among chicken birds. Spatial terms like “next” and “room,” and perspective-oriented words “other” and “side,” offer clues to the environmental and social contexts influencing the birds’ stress levels. The recurrence of specific terms demands an in-depth contextual understanding to elucidate their relevance to the birds’ prestress conditions.

##### 6.1.9.2. Disease Poststress Phase - Linguistic Patterns

Post-stress analysis indicates a continuous prominence of “nihai” alongside “going,” “kichi,” and “neko,” underscoring sustained themes or concerns following stress episodes. Increased occurrences of “other” and “side” suggest a possibly altered perspective or environment post-stress. The introduction of terms like “hikki” and “able,” and the maintained presence of “bathroom,” point towards changes in the birds’ capabilities or behaviors post-stress. This phase’s lexical landscape necessitates further investigation into the evolving or persistent stressors and their linguistic markers.

##### 6.1.9.3. Treatment Prestress Phase - Preparatory Linguistic Landscape

The term “nihai” dominates the prestress treatment phase, with an overwhelming 127,570 mentions, alongside terms denoting planning and spatial orientation like “going,” “next,” and “room.” The appearance of “nihaiya” introduces a variant or related concept, hinting at the complexity of the treatment anticipation phase. Words indicating alternative perspectives or preparations, such as “back,” “other,” and “side,” coupled with “kichi” and “school,” reflect the multifaceted preparations and considerations prior to treatment interventions. These findings suggest an intricate web of factors and anticipatory behaviors warranting comprehensive analysis.

##### 6.1.9.4. Treatment Poststress Phase - Post-Treatment Lexical Dynamics

In the aftermath of treatment interventions, “nihai” remains a central term, suggesting its critical relevance to the treatment discourse. Terms indicating transition and planning, such as “going, “room,” and “next,” along with “bathroom,” point towards ongoing adjustments or continued care following treatment. The persistence of “kichi” and “neko” implies lasting themes or concerns, while “back” suggests a return to normative behaviors or environments. This phase’s word frequencies uncover the sustained or new focal points post-treatment, highlighting th nuanced recovery or adaptation processes.

#### 6.1.10. Lexical Dynamics and Thematic Resonance in Chicken Vocalizations - A Comprehensive Unigram, Bigram, Trigram, and Visualization Analysis

##### 6.1.10.1. Unigram Analysis and Visualization Insights

The analysis of unigrams, encapsulated in word clouds, prominently highlights “nihai” as a pivotal term, commanding attention across all analyzed stages of chicken vocalizations. This term, along with “room,” “going,” “neck,” and “kich,” forms a core lexical set that recurrently appears, indicating these words’ fundamental importance in the discourse on chicken vocalizations during different stress and treatment conditions. Intriguingly, the analysis reveals a richer lexicon in the phases of disease prestress and treatment prestress, suggesting these periods provoke a broader range of vocal expressions. The consistent presence of these terms, albeit with varying frequencies across the phases, marks them as key elements within the discourse, suggesting their significance in the context of chicken vocalizations and their reactions to disease and treatment processes.

##### 6.1.10.2. Bigram Exploration and Patterns

Delving into bigrams unveils a sophisticated linguistic structure, with “nihai nihai” markedly repeated, signifying a thematic focus or emphasis pertinent across the disease and treatment narratives. The pairs “neko kichi” and “kichi neko” emerge with frequency, implying a significant thematic or conceptual linkage between these terms. Sequential and spatial narratives are illuminated through bigrams such as “going next,” “next room,” and “room going,” reflecting narratives of progression, movement, or transition. Moreover, the associative combinations involving “other,” “side,” and “going” in various permutations suggest thematic explorations of perspective, dimensionality, or relational dynamics within the discourse. These insights illuminate the rich and interconnected linguistic landscape of chicken vocalizations, revealing the depth of thematic engagement and narrative construction across the stages of disease and treatment.

##### 6.1.10.3. Trigram Analysis and Thematic Insights

The examination of trigrams offers an enriched perspective on the thematic and linguistic intricacies within chicken vocalizations. The dominant recurrence of “nihai nihai nihai” underscores its thematic prominence, echoing through the dataset as a central motif. Trigrams like “neko kichi neko” and “kichi neko kichi” denote a complex thematic interplay, suggesting intricate relationships or thematic continuities. Sequential trigrams such as “going next room” and “next room going” depict clear narrative progressions, indicating actions or transitions emblematic of the different phases explored. The presence of trigrams involving “other,” “side,” and “going” in various configurations points to a nuanced exploration of spatial and relational themes, enriching our understanding of the discourse. Through trigram analysis, we delve deeper into the layered linguistic and thematic fabric of chicken vocalizations, highlighting the nuanced expressions and thematic diversity present across states.

##### 6.1.10.4. Visual Representation of Lexical Patterns

The word clouds created for unigrams, bigrams, and trigrams (Figure 9A, 9B, 9C) serve as visual manifestations of the study’s findings, illustrating the prominence and frequency of linguistic patterns and thematic elements. These visualizations effectively highlight the recurrent linguistic structures and thematic motifs, providing a compelling overview of the core elements that characterize the discourse on chicken vocalizations across various conditions. The visual analysi brings to light the thematic coherence and linguistic richness of the corpus, offering a vivid portrayal of the underlying patterns that shape chicken vocalizations through the phases of disease and treatment.

**Figure 7.**
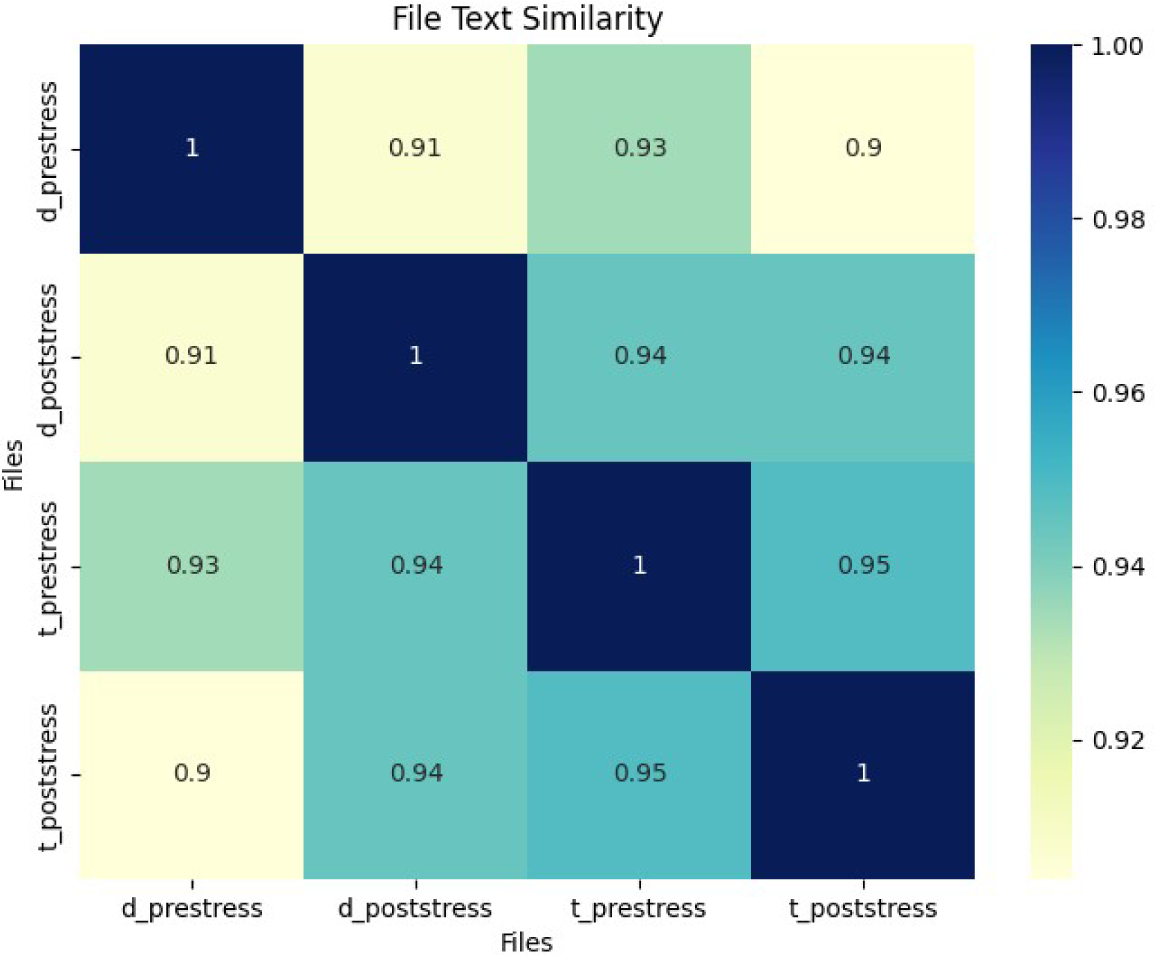
Cosine Similarity Matrix Map depicting Textual Similarity among Disease and Treatment Phases. The Y-axis enumerates the files under comparison: Treatment Poststress (t_poststress), Treatment Prestress (prestress), Disease Poststress (d_poststress), and Disease Prestress (d_prestress). The X-axis mirrors these files, facilitating a comprehensive visual analysis of the degrees of similarity across different stages of disease and treatment, quantified through cosine similarity metrics.

**Figure 8.**
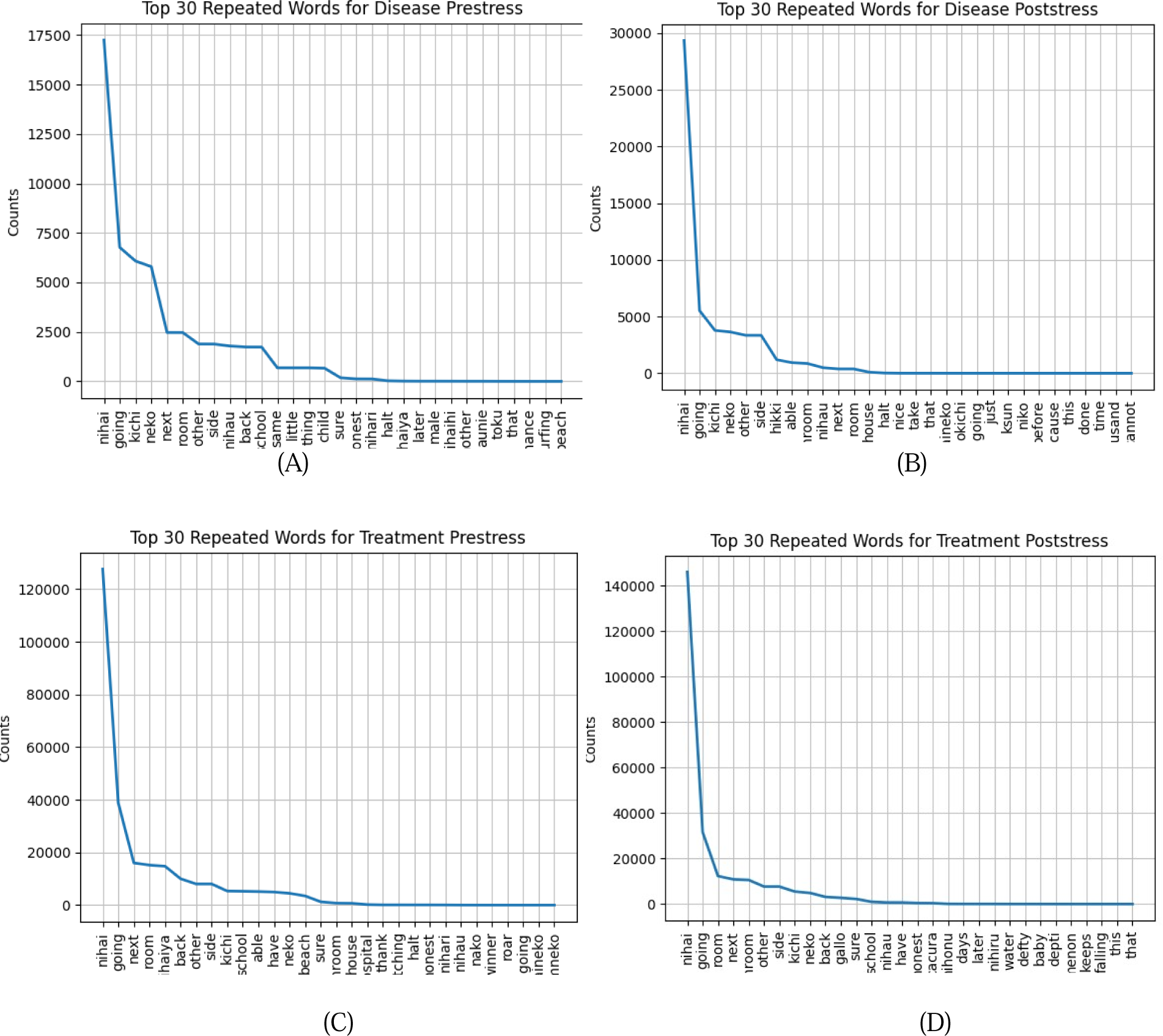
Comparative Analysis of the Top 30 Most Frequently Occurring Words Across Four Phases: Disease Prestress (A), Disease Poststress (B), Treatment Prestress (C), and Treatment Poststress (D). Thi consolidated figure juxtaposes the counts (Y-axis) of the top 30 words identified in each phase (X-axis) within chicken birds’ vocalizations, offering a comprehensive overview of lexical variations and consistencies through the disease and treatment cycles.

**Figure 9A.**
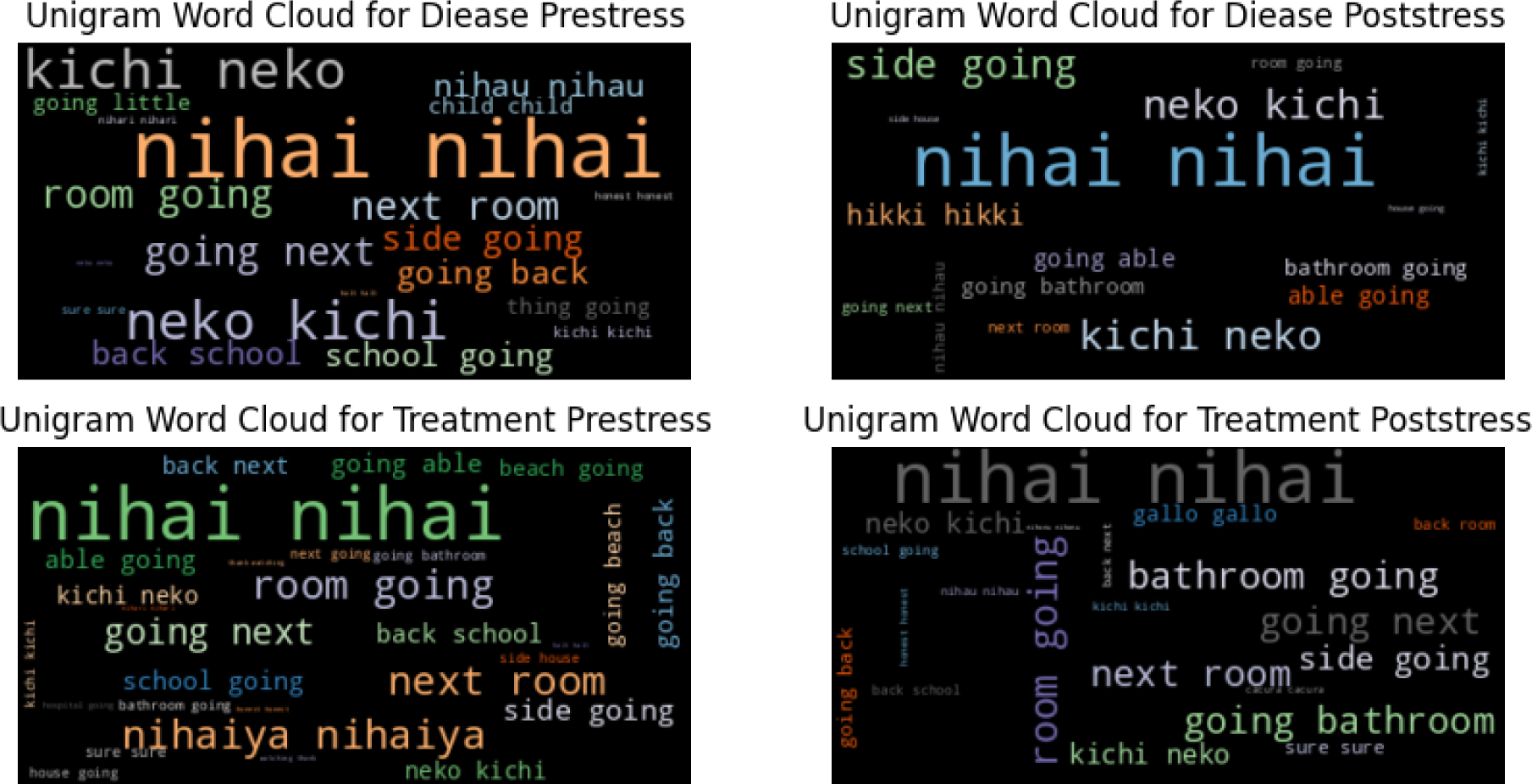
Unigram Word Cloud for Chicken Vocalizations Across Phases – This visualization encapsulates the unigram analysis of chicken vocalizations through different stages of disease and treatment, spotlighting dominant terms like “nihai” and their prevalent use across contexts.

**Figure 9B.**
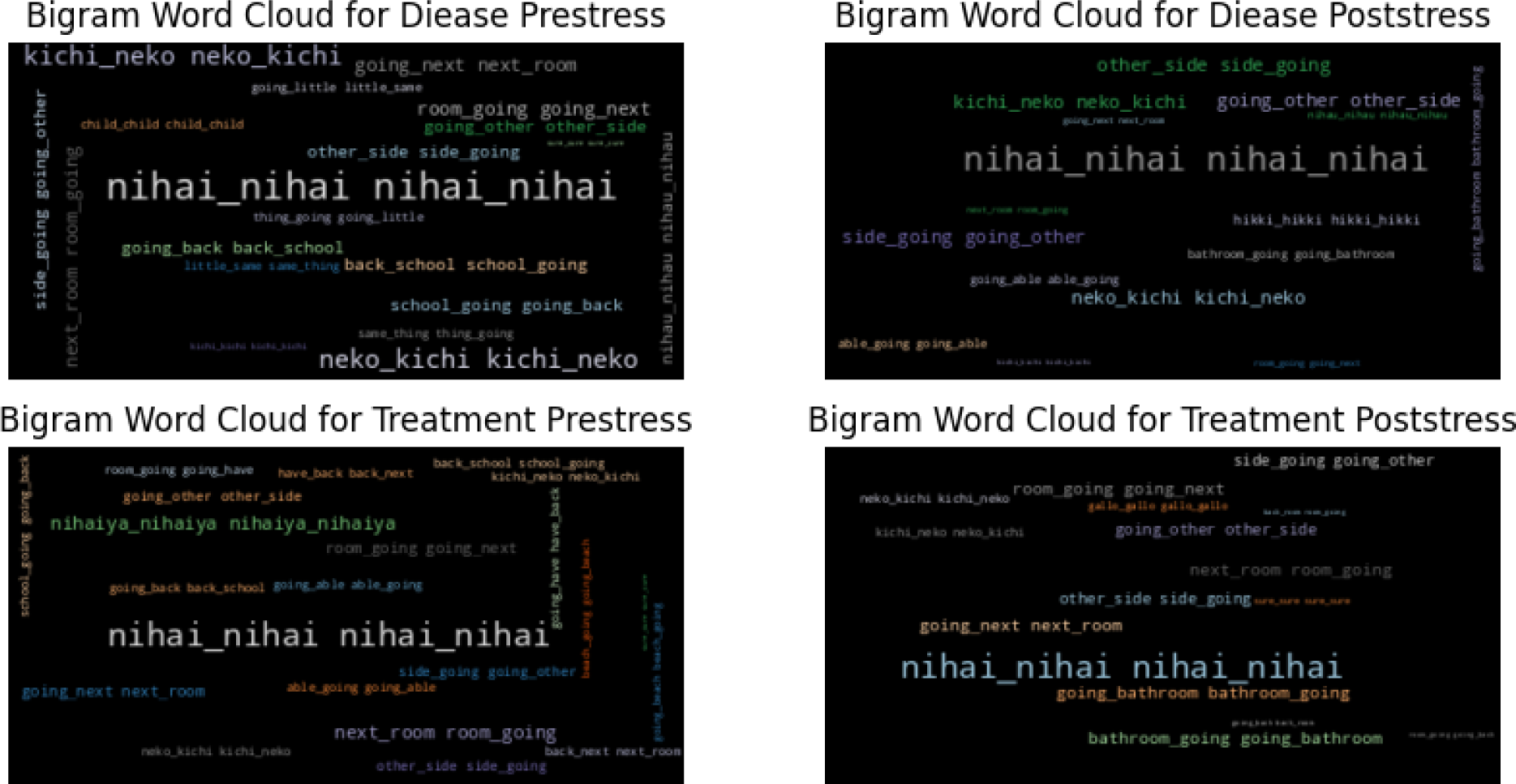
Bigram Word Cloud for Chicken Vocalizations Across Conditions – This word cloud delves into the bigram analysis, shedding light on the pivotal pairings of terms. It visually articulates the significant thematic links and linguistic connections that manifest through varying stress and treatment scenarios in chicken vocalizations.

**Figure 9C.**
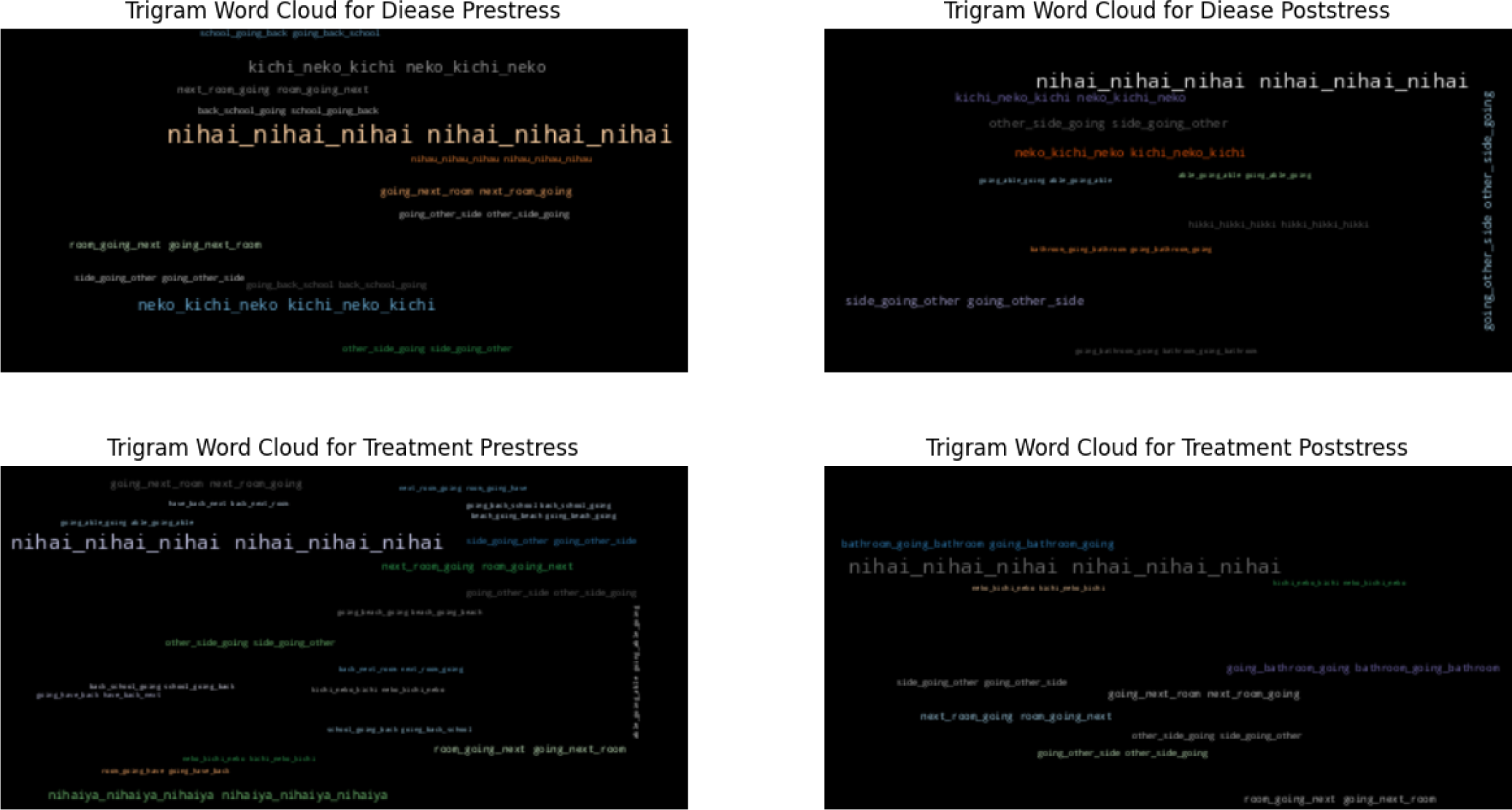
Trigram Word Cloud for Chicken Vocalizations Across Stages – Offering a deep dive into trigram analysis, this visualization uncovers the intricate linguistic frameworks and thematic continuities that characterize chicken vocalizations. It highlights the complexity and thematic depth within the dataset, showcasing prevalent trigrams through diverse phases of disease and treatment.

#### 6.1.11 Linguistic Characteristics Across Treatment and Disease Phases - An Analysis of Parts of Speech Frequencies

##### 6.1.11.1. Disease Pre-stress Analysis

The linguistic structure during the disease pre-stress phase is characterized by a diverse array of parts of speech (Figure 10). Adjectives and present participle verbs are most prevalent, with 6,607 and 6,782 instances respectively, highlighting the descriptive and ongoing nature of the discourse. Nouns are also prominent, with 8,412 appearances, providing a solid foundation for the narrative. Adverbs, with 1,756 instances, enrich descriptions of actions and qualities. Foreign words are particularly notable, appearing 29,609 times, indicating a rich integration of diverse linguistic elements. Less frequent parts of speech, such as comparative adverbs and conjunctions, appear sparingly, reflecting their minimal role in this context. This linguistic pattern offer detailed insights into the narrative styles and syntactic preferences that dominate discussions about disease pre-stress, with a balanced use of descriptive and dynamic language.

**Figure 10.**
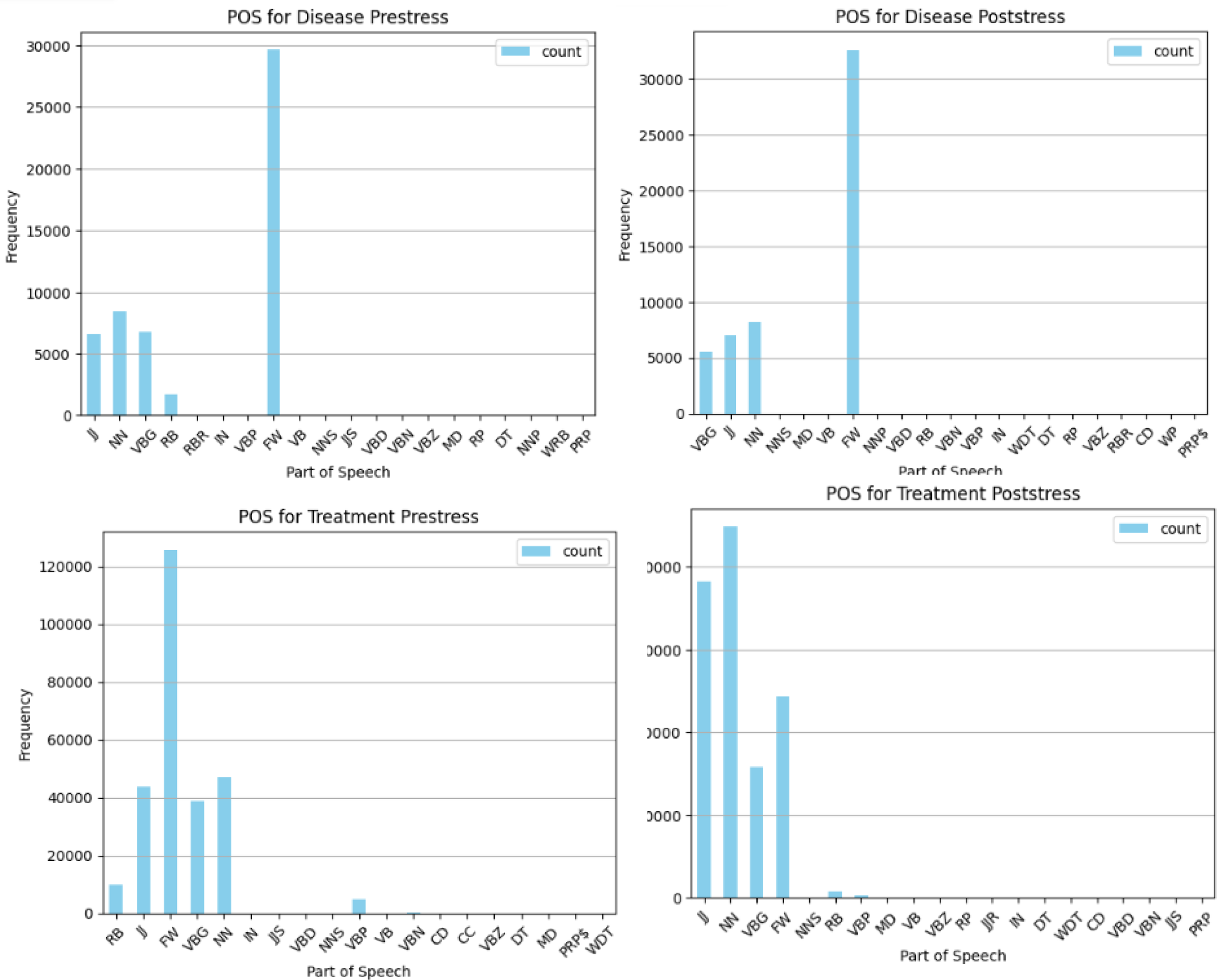
A-D: Comparative Distribution of Parts of Speech in Chicken Vocalizations Across Disease and Treatment Phases. These graphs illustrate the frequency distribution of parts of speech for Disease Prestress (A), Disease Poststress (B), Treatment Prestress (C), and Treatment Poststress (D) in chicken vocalizations, obtained via Whisper model analysis. The graphs highlight variations in linguistic composition across different phases.

##### 6.1.11.2. Disease Post-stress Composition

In the disease post-stress phase, the linguistic composition evolves subtly. Present participle verbs, adjectives, and nouns continue to be highly frequent, supporting ongoing actions and detailed descriptions within the discourse. The count of foreign words increases to 32,616, further emphasizing the linguistic diversity. The occurrence of modal and base verbs, although less frequent, introduces variations in verbal forms used to express conditionalities or hypothetical scenarios. Less common parts of speech, such as cardinal numbers and possessive pronouns, are used selectively, enhancing a focused narrative tailored to post-stress conditions. This phase maintains a rich tapestry of linguistic elements, continuing a detailed and dynamic narrative style that reflects the complexities encountered after disease.

##### 6.1.11.3. Treatment Pre-stress Linguistics

The treatment pre-stress phase shows a marked increase in the frequency of all parts of speech, particularly adjectives and nouns with 43,816 and 47,048 occurrences respectively. This indicates a heightened emphasis on descriptive and nominal elements, likely reflecting thorough preparations and discussions related to impending treatments. Present participle verbs also feature strongly, emphasizing ongoing processes or actions. An extraordinary presence of foreign words, totaling 125,635 occurrences, underscores an expanded diversity in language usage, possibly including medical or technical terminology. Less frequent parts of speech, such as prepositions and modal verbs, although less common, contribute to a rich linguistic profile that supports a nuanced exploration of anticipatory narratives regarding treatment.

##### 6.1.11.4. Treatment Post-stress Linguistic Trends

In the treatment post-stress phase, the linguistic landscape is richly populated with descriptive language, evidenced by 76,527 occurrences of adjectives. Nouns and present participle verbs continue to dominate, underscoring a focus on identifying entities and detailing ongoing actions related to post-treatment experiences. The substantial count of foreign words, 48,638, continues to reflect a broad linguistic spectrum, likely encompassing various medical and recovery-related terms. The infrequent use of modal verbs and particles, along with minimal use of conjunctions and determiners, indicates a selective syntactic style that prioritizes specific narrative elements. This phase showcases a sophisticated use of language to articulate the complexities and outcomes of post-treatment scenarios, encapsulating the diverse aspects of recovery and ongoing care.

#### 6.1.12 Topic Modeling Analysis Across Disease and Treatment Phases in Chicken Vocalizations

##### 6.1.12.1. Topics for Disease Prestress

The topic modeling for disease prestress reveals a diverse set of themes, where each topic encapsulates a specific aspect of the discourse (Figure S1). Topics involving family and relational terms (Topic 0, Topic 5) highlight familial roles and dynamics, evidenced by terms like “father,” “child,” and “nihari,” pointing to the emotional and relational discussions prevalent during the prestress phase. Spatial and action-oriented topics (Topic 1, Topic 6) include words like “side,” “back,” and “going,” suggesting a narrative focus on movement and spatial orientation. Themes of assurance and affirmation (Topic 2, Topic 7) are marked by words such as “sure,” “honest,” and “nihai,” indicating discussions centered around certainty and reassurance within the context. Mixed personal and spatial references (Topic 3, Topic 4) merge personal interactions with physical settings, as seen with terms like “room,” “next,” and family-related words, suggesting a blend of personal and locational narratives. The diverse topics underline the complexity of the linguistic landscape in the disease prestress phase, showing a nuanced interplay of themes from familial interactions to descriptions of space and actions.

Topic 0 through Topic 9 reveal central themes with dominant terms like “nihai” and “going” indicating key focus areas in discussions. These terms are surrounded by less frequent but contextually significant words, which add depth and specificity to each topic. For example, “school” and “father” appear across topics at varying frequencies, emphasizing their relevance in different thematic contexts. Inter-topic connections are evident where similar terms appear across multiple topics, linking themes such as movement and family dynamics, which helps in understanding the broader narrative connections within the prestress discussions.

##### 6.1.12.2. Topics for Disease Poststress

Poststress topics transition slightly in thematic focus while retaining some continuity from the prestress phase (Figure S2). Continued focus on family and relations (Topic 0, Topic 5) with terms like “father” and “child” persists, illustrating ongoing familial discussions post-stress. Introduction of terms related to personal care and hygiene (Topic 1, Topic 8) such as “bathroom,” suggest a shift towards more practical and recovery-oriented discussions. Emphasis on capability and action (Topic 2, Topic 7) as indicated by terms like “able” and “going,” points to a narrative of recovery and progression. The transition from prestress to poststress topics reflects a shift in narrative focus from immediate emotional responses to recovery and practical issues, maintaining a detailed and dynamic discourse that mirrors the evolving contexts of disease progression and recovery.

##### 6.1.12.3. Topics for Treatment Prestress

High frequency of descriptive and nominal elements (Topic 0, Topic 5) like “nihai” and “school” underscore a focus on detailed descriptive narratives and specific planning discussions related to upcoming treatments (Figure S3). Significant mentions of process and action terms (Topic 3, Topic 6) such as “next” and “going,” highlight ongoing actions and processes pertinent to treatment preparation. This phase shows an enriched linguistic usage that reflects intensive preparation and anticipatory discussions, characterized by a broad and complex use of language that spans descriptive, nominal, and action-oriented discourse.

##### 6.1.12.4. Topics for Treatment Poststress

Continued emphasis on descriptive terms (Topic 0, Topic 9) with “nihai” and “water” suggesting a focus on outcomes and ongoing conditions post-treatment (Figure S4). Terms indicating recovery and practical adjustments (Topic 4, Topic 8) such as “bathroom” and “back,” which are indicative of discussions around recovery and returning to normalcy. This final phase demonstrates a sophisticated narrative that incorporates a wide array of descriptive and practical terms, reflecting the complexities of post-treatment experiences and ongoing care discussions, effectively capturing the diverse linguistic elements used to convey these nuanced aspects.

#### 6.1.13. Inter-Topic Distance Mapping of Chicken Vocalizations - A Multidimensional Scaling Analysis

##### 6.1.13.1. Disease Prestress

The multidimensional scaling analysis for disease pre-stress vocalizations delineates twenty distinct topics. Topics 1, 2, and 3 display significant thematic divergence from each other, with the bulk of the topics—seventeen in total—forming a closely positioned cluster within the plot. Topic 3 is uniquely closer to this large cluster, indicating subtle thematic connections or overlaps with the general discourse observed in the pre-stress phase.

##### 6.1.13.2. Disease Poststress

During the poststress phase, the inter-topic distance map reveals noticeable variability among the analyzed topics. Topic 1 is particularly distinct, markedly separated from the others, which highlights unique thematic concerns following disease-induced stress. A cluster comprising Topics 4, 5, 7, 11, 15, and 17 is tightly grouped, reflecting close thematic similarities. The rest of the topics largely merge into a significant overlapping cluster, indicating a high degree of thematic resonance across these discussions.

##### 6.1.13.3. Treatment Prestress

For treatment pre-stress, the analysis charts twenty topics, with Topics 1 and 2 showing clear thematic distinctions from each other. The other seventeen topics display extensive thematic overlap, merging visually into a single coherent area on the map. This suggests a high degree of uniformity in the discussion themes related to treatment anticipation, mirrored in their proximal representation on the plot.

##### 6.1.13.4. Treatment Poststress

The post-treatment analysis also maps twenty topics, where Topics 1 and 2 again show distinct thematic elements. The other seventeen topics, while closely overlapping, are positioned diametrically opposite to their configuration in the pre-stress phase on the plot. This inversion may reflect a shift in narrative focus or thematic concerns following the treatment, as indicated by the significant repositioning of these topics. Figure S5 underscore the complex interplay of themes and discussions across various stages of disease and treatment in chicken vocalizations, offering a nuanced understanding of the shifts and continuities in the narrative through advanced modeling techniques.

### 6.2. Results from Dataset 2

#### 6.2.1. Examination of Character Occurrences

In the “healthy” category, the comma (“,”) is the most frequent character, appearing approximately 14,000 times, closely followed by the characters “5” and “o,” each occurring around 12,000 times. Characters such as “f,” “w,” “j,” and “2” are notably less common. For the “noisy” files, the character “o” leads with an impressive 25,000 occurrences. It is followed closely by “i,” “t,” and “,”, each with about 20,000 occurrences. Conversely, “j” and “0” are among the least occurring characters in this category. In the “unhealthy” category, “o” once again predominates with over 25,000 appearances. Other frequently occurring characters include “1,” “.”, “5,” “a,” “g,” and “e,” each nearing 15,000 occurrences. Remarkably, “j” remains one of the least frequent characters even in this set. A consistent pattern of character dominance and rarity is observed across the three categories—healthy, noisy, and unhealthy—highlighting uniform character usage despite the varied acoustic environments. This consistency (Figure 11) suggests a distinctive textual or meta-data characteristic shared across the files, reflecting uniform tagging or encoding practices irrespective of the sound type or recording conditions.

**Figure 11.**
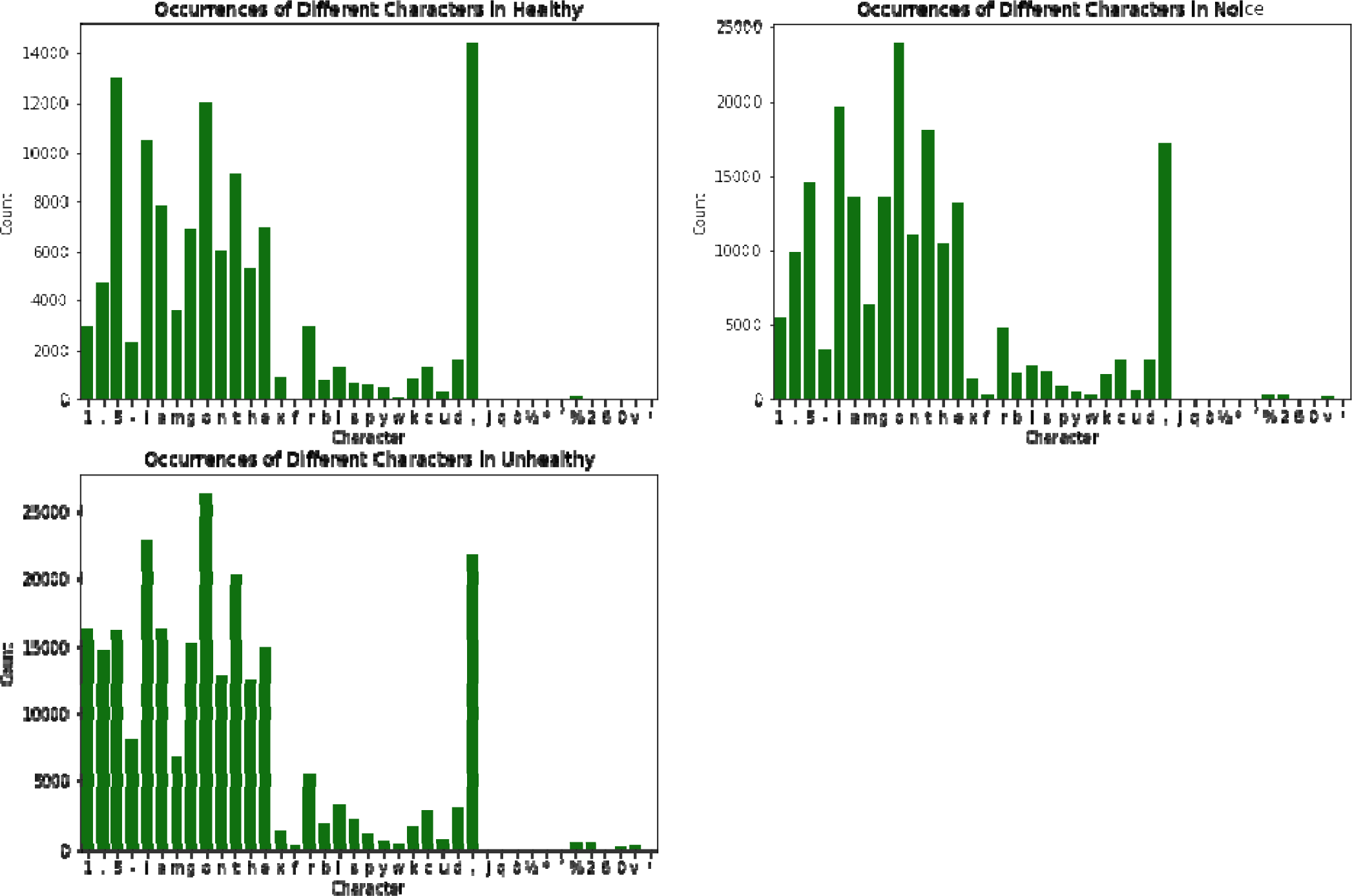
Character Frequency Analysis Across Three Phases – (A) Healthy, (B) Noise and (C) Unhealthy

#### 6.2.2. Analytical Insights into Textual Composition - Evaluating Linguistic Metrics Across Healthy, Noisy, and Unhealthy Phases of the Chicken Vocalization Dataset

In the “healthy” phase of the dataset, the textual composition encompasses approximately 150,000 characters, with an evenly distributed linguistic structure consisting of about 50,000 vowels and 50,000 consonants. Additionally, this phase includes around 10,000 stopwords, illustrating a baseline complexity in the text’s composition.

Transitioning to the “noise” phase, the dataset expands significantly, containing roughly 250,000 characters. Despite the increase in total character count, the proportion of vowels and consonants remains consistent with the “healthy” dataset, each maintaining a count of about 50,000. However, there is a substantial increase in the number of stopwords, which surge to approximately 100,000, indicating a higher density of common, less informative words that may reflect the chaotic nature of the background noises captured in this phase. In the “unhealthy” phase, the dataset further enlarges to encompass around 300,000 characters. This phase sees doubling in the number of vowels and consonants, totaling approximately 100,000 each. The stopword count also increases but remains considerably lower than the “noise” phase, tallying around 20,000 stopwords. This suggests a denser and potentially more complex textual structure, reflecting the severity of the conditions under which these vocalizations were recorded.

Across all three phases—healthy, noise, and unhealthy—a consistent pattern in the growth of textual elements is evident (Figure 12). However, the “unhealthy” dataset is distinguished by a marked increase in both vowels and consonants compared to the “noise” dataset, highlighting an amplification in the complexity and diversity of textual elements. This escalation may correlate with the increased expressiveness or distress in the vocalizations associated with the unhealthy conditions of the subjects.

**Figure 12.**
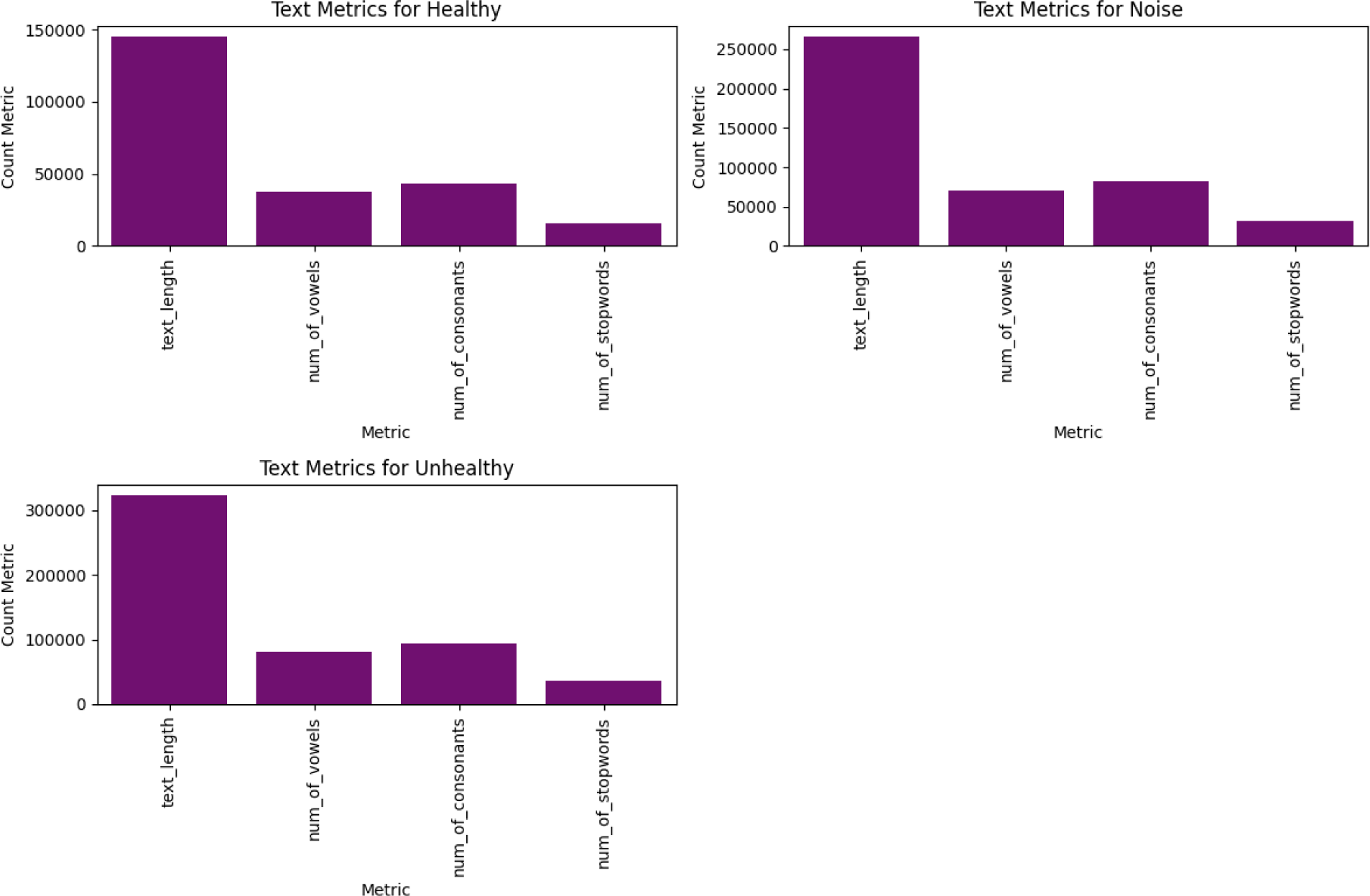
Comparative Analysis of Text Metrics - Text Length, Number of Vowels, Number of Consonants, and Number of Stopwords Across Healthy, Noise and Unhealthy phase.

#### 6.2.3. Phonetic Composition Analysis: Dissecting the Roles of Vowels and Consonants Across Healthy, Noise, and Unhealthy Phases of Chicken Vocalizations

##### 6.2.3.1. Vowel Distribution

In the “healthy” dataset, the analysis of vowel occurrences (Figure 13) reveals that the vowels “a” and “e” are present between 5,000 and 10,000 times each, whereas “i” and “o” appear approximately 10,000 times, with “o” being the most frequently occurring vowel. Notably, “u” shows the lowest frequency among the vowels. Transitioning to the “noise” dataset, the occurrences of “a” and “e” remain stable at around 10,000 instances each. However, “i” and “o” show a significant increase, with each appearing around 20,000 times. In this dataset, “o” notably surpasses all other vowels, reaching over 20,000 occurrences, while “u” continues to be the least frequent.

**Figure 13.**
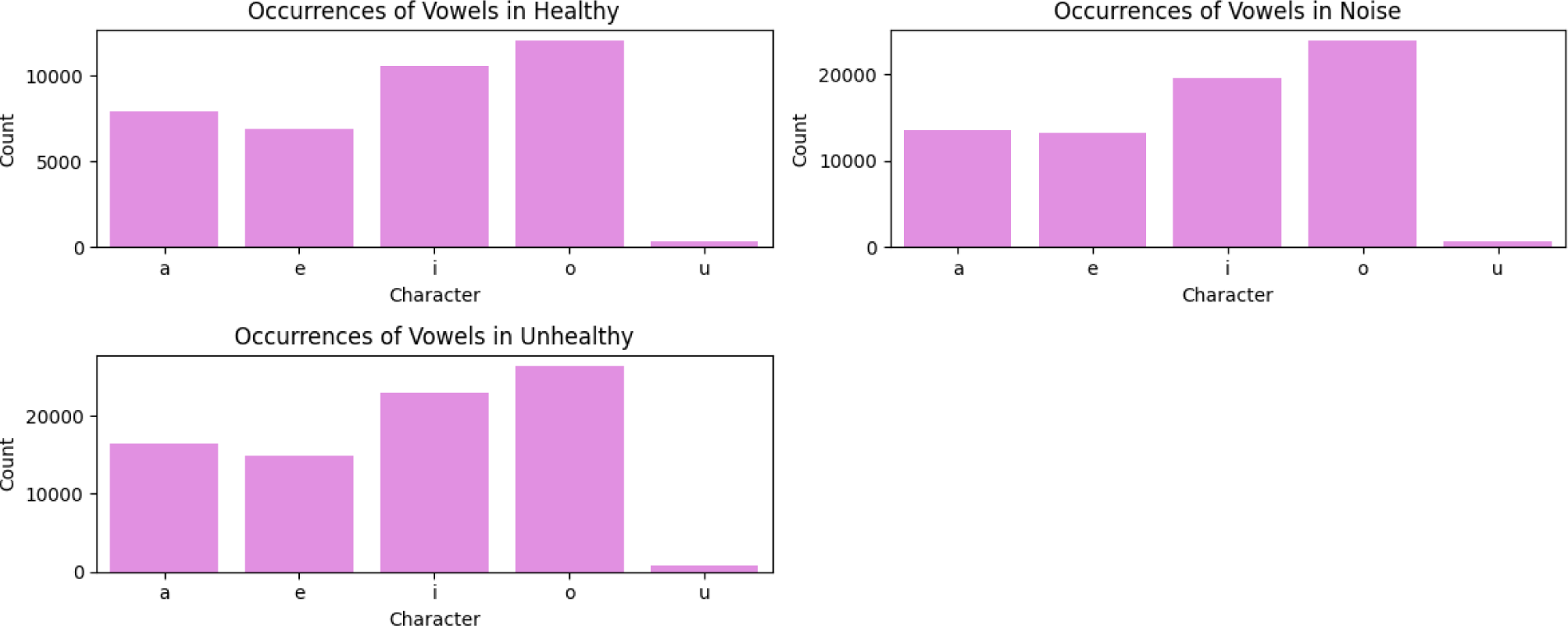
Distribution of Vowel Frequencies Across Different Stages of Chicken Vocalization - Healthy, Noise, and Unhealthy Dataset Analysis

In the “unhealthy” dataset, the frequency of vowels “a” and “e” rises to between 10,000 and 20,000, indicating an increase in vocal activity or stress. Both “i” and “o” exceed 20,000 occurrences, with “o” consistently maintaining its dominance as the most frequent vowel. The trend continues with “u” remaining the least occurring vowel across all datasets. This consistent pattern across the datasets highlights the distinctive phonetic characteristics preserved despite the variations in environmental and health conditions.

##### 6.2.3.2. Consonant Distribution

In the “healthy” dataset, consonants display varied frequencies (Figure 14), with “g,” “h,” and “n” each recorded around 6,000 times, whereas “m” and “r” are noted approximately 4,000 times each. The consonant “t” stands out significantly, being the most frequent with over 8,000 occurrences, while “w” ranks as the least frequent.

**Figure 14.**
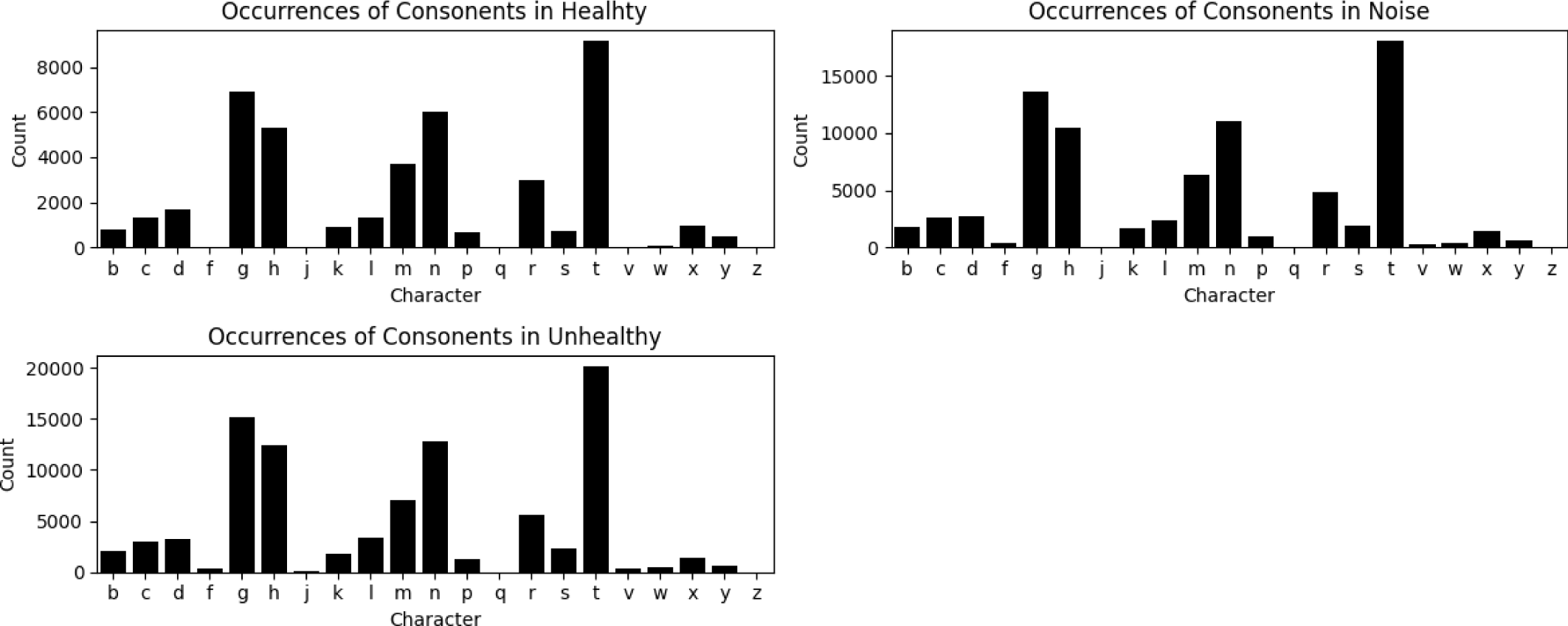
Consonant Frequency Analysis in Various Phases of Chicken Vocalizations - Healthy, Noise, and Unhealthy Dataset Comparisons

Shifting to the “noise” dataset, both “h” and “n” see an increase in occurrences to about 10,000 each, while “m” and “r” display a slight increase to around 5,000 occurrences each. Notably, “t” remains the most dominant consonant, escalating to over 15,000 repetitions, and “v” emerges as the least frequent consonant.

In the “unhealthy” dataset, “g” leads with 15,000 occurrences, while “h” and “n” range between 10,000 and 15,000. Once again, “t” captures the spotlight with approximately 20,000 occurrences, underlining its persistent dominance across all phases. Conversely, “j” is the least prevalent consonant.

Across all three datasets—healthy, noise, and unhealthy—a consistent pattern is evident, with “t” maintaining its position as the most frequently occurring consonant. This consistency highlights a remarkable stability in the use of consonants despite the variations in dataset characteristics, suggesting a foundational structure in the phonetic composition of the chicken vocalizations across different environmental and health contexts.

#### 6.2.4. Emotional Dynamics Across Healthy, Noise and Unhealthy Phases - A Sentiment Analysis

In the “healthy” dataset, sentiments predominantly lean towards positivity, comprising 82% positive sentiments, complemented by 16% negative sentiments and a minimal presence of neutral sentiments. This trend of positivity is mirrored in the “noise” dataset, where positive sentiments dominate at 77%, followed by 22% negative sentiments, with only a few neutral instances. The “unhealthy” dataset shows a similar pattern, with 78% positive sentiments, 21% negative sentiments, and an insignificant amount of neutral sentiments. Across all three datasets (Figure 15)—healthy, noise, and unhealthy—a consistent pattern is observed: positive sentiments are predominant, neutral sentiments are scarce, and negative sentiments are less frequent. This consistent trend of positivity highlights a remarkable uniformity in sentiment distribution, regardless of the varying conditions of the dataset contexts.

**Figure 15.**
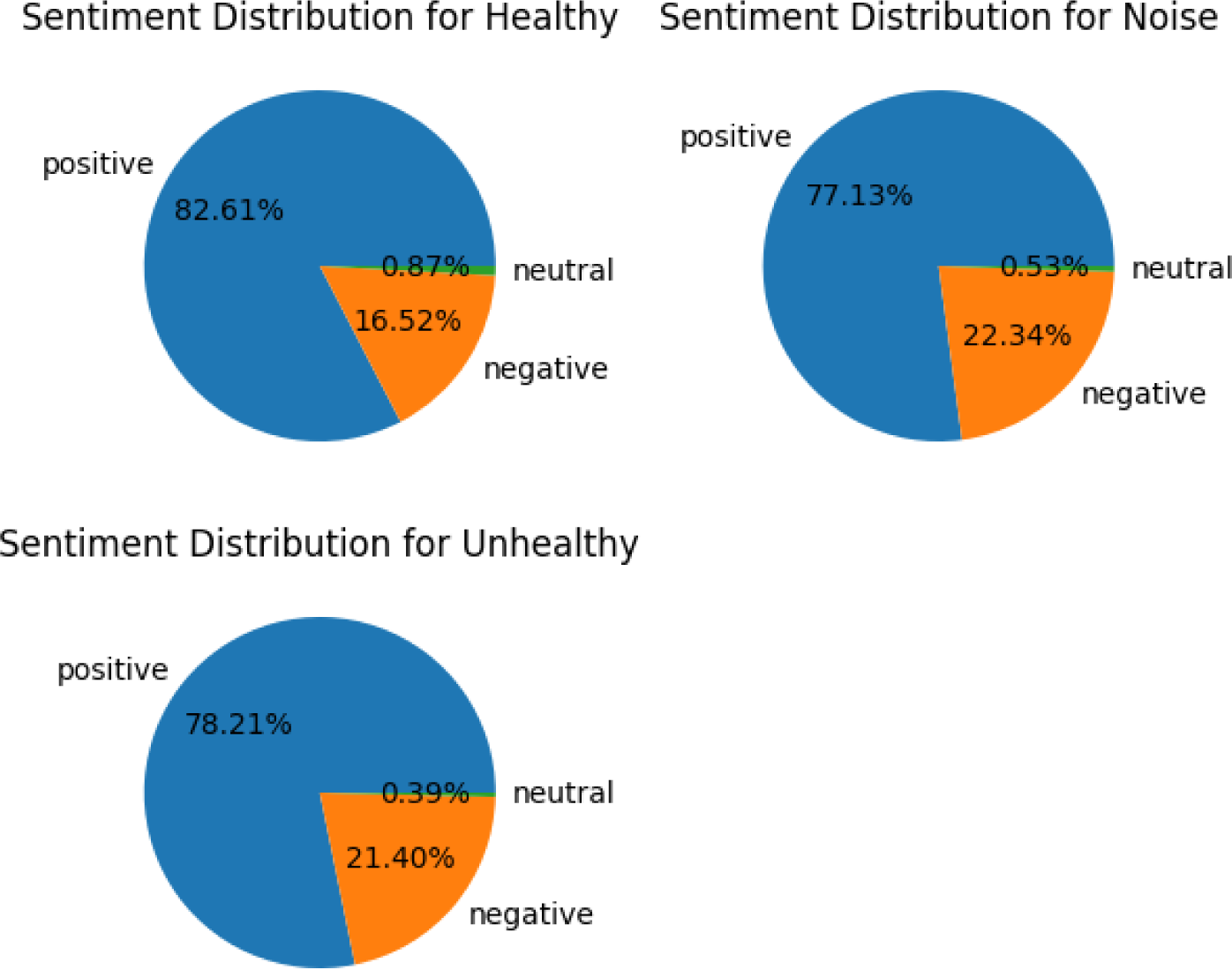
Comparative Sentiment Distribution Across Healthy, Noise, and Unhealthy Phases of the Chicken Vocalization Dataset

#### 6.2.5. Analyzing Sentiment Dynamics Across Healthy, Noise, and Unhealthy Phases: A Comparative Study Using Displots and Boxplots

##### 6.2.5.1. Sentiment Score Distribution Displot Analysis

The distribution of sentiment scores for healthy dataset ranges from 0.37 to 0.8, with the bulk of the scores falling between 0.37 and 0.60, indicating a prevalent positivity within the dataset. The less frequent occurrence of scores between 0.7 and 0.8 suggests fewer instances of highly positive sentiments. Scores for the Noise dataset again range from 0.37 to 0.8. Like in the healthy dataset, the majority of scores are concentrated between 0.37 and 0.60. There is a noted decrease in scores above 0.70, indicating a drop in extremely positive sentiments, which aligns with the potentially disruptive nature of the noise context. For the Unhealthy Dataset,scores span a similar range of 0.37 to 0.8, with the highest density around 0.40, suggesting the presence of predominantly moderately positive sentiments. This distribution mirrors the other datasets, emphasizing a consistent sentiment trend across varying health conditions. Overall, across all three datasets, the most frequent sentiment scores are noted within the 0.4 to 0.5 range, highlighting a universal trend of positive sentiments regardless of the dataset’s health or environmental context. Figure 16A illustrates the distribution of sentiment scores within the healthy, noise, and unhealthy phases of the dataset. Displots reveal the concentration and spread of sentiment values, highlighting the prevalence of moderately positive sentiments across the different contexts and showcasing the variations in sentiment intensity among the datasets.

**Figure 16A:**
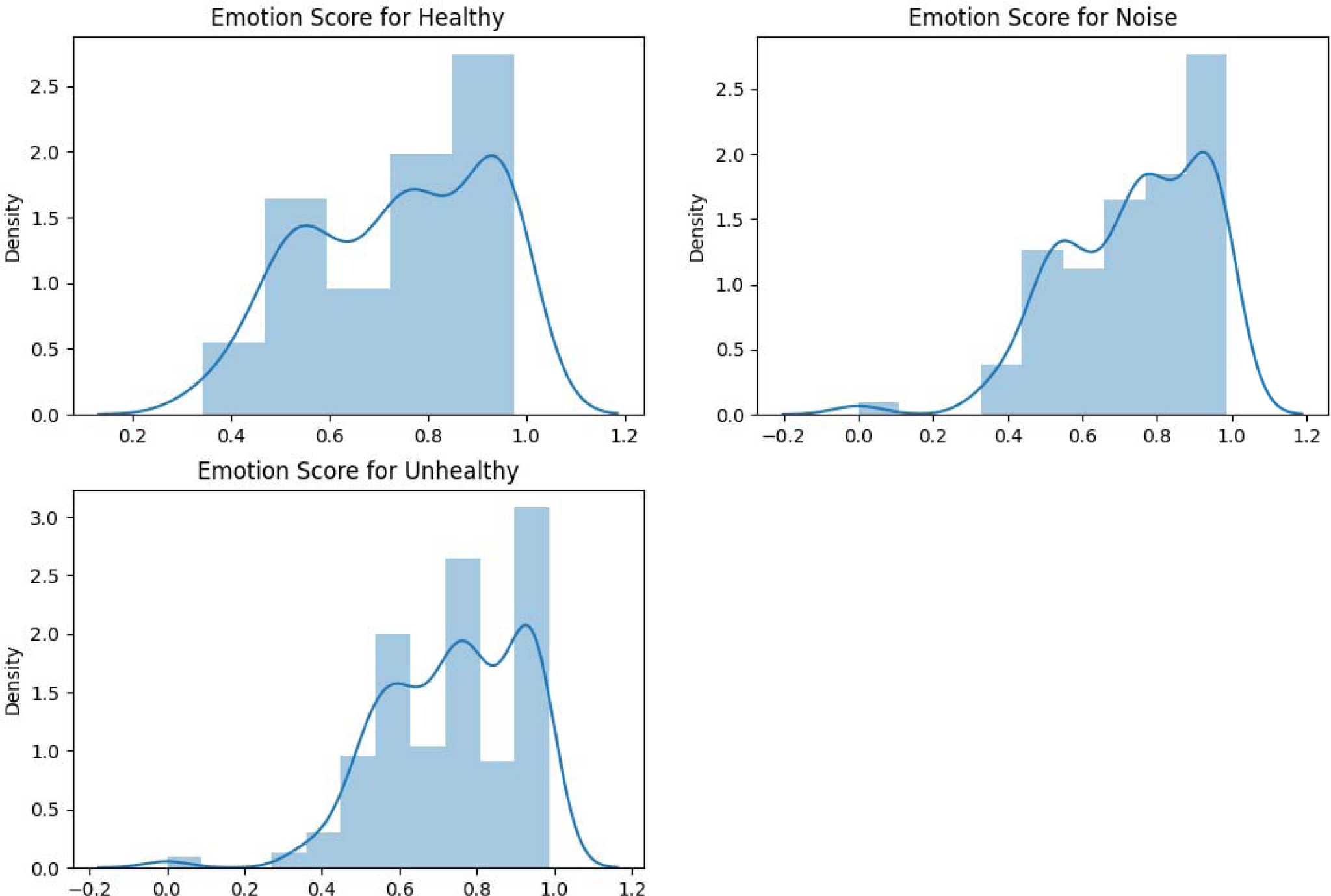
Distribution of Sentiment Scores Across Healthy, Noise, and Unhealthy Phases - An Analysis Using Displots

##### 6.2.5.2. Sentiment Score Distribution Using Boxplot Analysis

The boxplot for healthy dataset reveals a distribution of sentiment scores from 0.3 to 0.7, with half of the data falling between 0.3 and 0.5. The median sentiment score stands at 0.5, indicating that the typical sentiment is moderately positive. An outlier at 0.8 points to occasional highly positive sentiments. For the Noise Dataset, the spread also ranges from 0.3 to 0.7, with 50% of the scores below 0.50 and the median again at 0.5. A segment of the dataset, about 25%, records scores below 0.42, highlighting a variance in sentiment intensity. An outlier at 0.8 reflects extreme positive sentiments, similar to the healthy dataset. While for the Unhealthy Dataset, the range extends from slightly less than 0.4 to 0.7. With half of the data below a sentiment score of 0.50 and a consistent median of 0.5, the pattern seen in previous datasets persists, suggesting stability in sentiment distribution across different contexts. The presence of an outlier at 0.8 is also noted here. The consistent distribution pattern, median values, and presence of outliers across all datasets suggest a stable trend in sentiment dynamics, highlighting how sentiments, while predominantly positive, show slight variations reflective of the different dataset conditions. This uniformity in sentiment distribution across diverse contexts underscores a commonality in emotional expression among the datasets. Figure 16B presents the spread of sentiment scores through boxplots for the healthy, noise, and unhealthy phases of the dataset. The boxplots elucidate the median, quartiles, and outliers within each dataset phase, illustrating the consistency and variance in sentiment distributions across different environmental and health conditions.

**Figure 16B.**
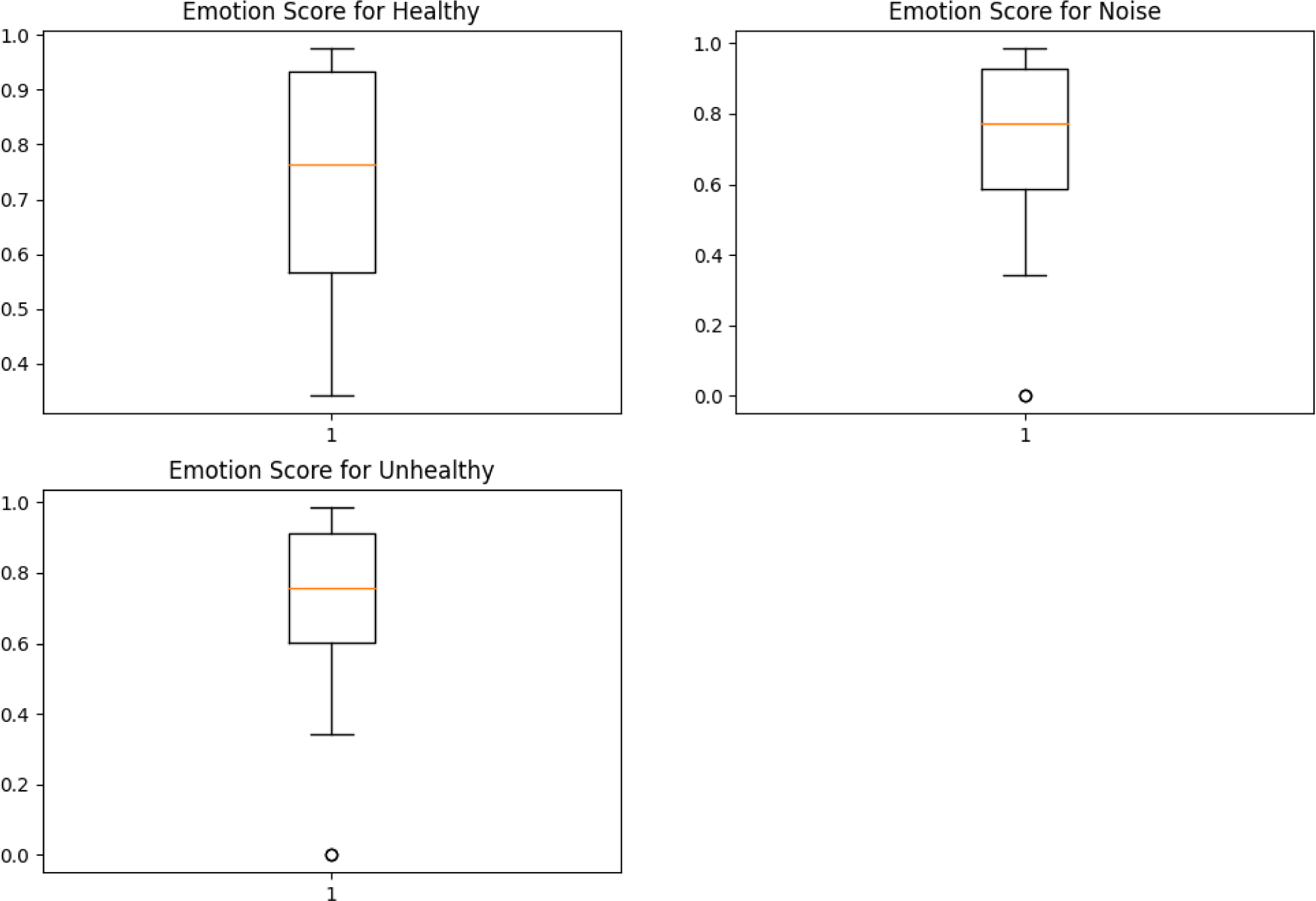
Distribution of Sentiment Scores Across Healthy, Noise, and Unhealthy Phases - An Analysis Using Boxplots

#### 6.2.6. Analytical Insights into Sentiment and Emotion Dynamics Across Healthy, Noise, and Unhealthy Phases of Chicken Vocalization Dataset

##### 6.2.6.1. Sentiment Correlation

There is a discernible positive correlation among the sentiment scores across all datasets derived from the analysis of chicken vocalizations. This correlation is evident from the diagonal trends observed in the correlation matrices, which indicate a robust positive correlation between the sentiment scores of different contexts. Specifically, a positive correlation is noted between the “healthy” and “noise” datasets, as well as between the “healthy” and “unhealthy” datasets. This suggests that as sentiment scores increase in one dataset, they tend to similarly increase in another, reflecting a consistent pattern (Figure 17A) of sentiment distribution across different conditions of chicken vocalizations. This uniform sentiment expression across various datasets underscores common emotional responses despite differences in the health status or environmental background of the chickens.

**Figure 17A.**
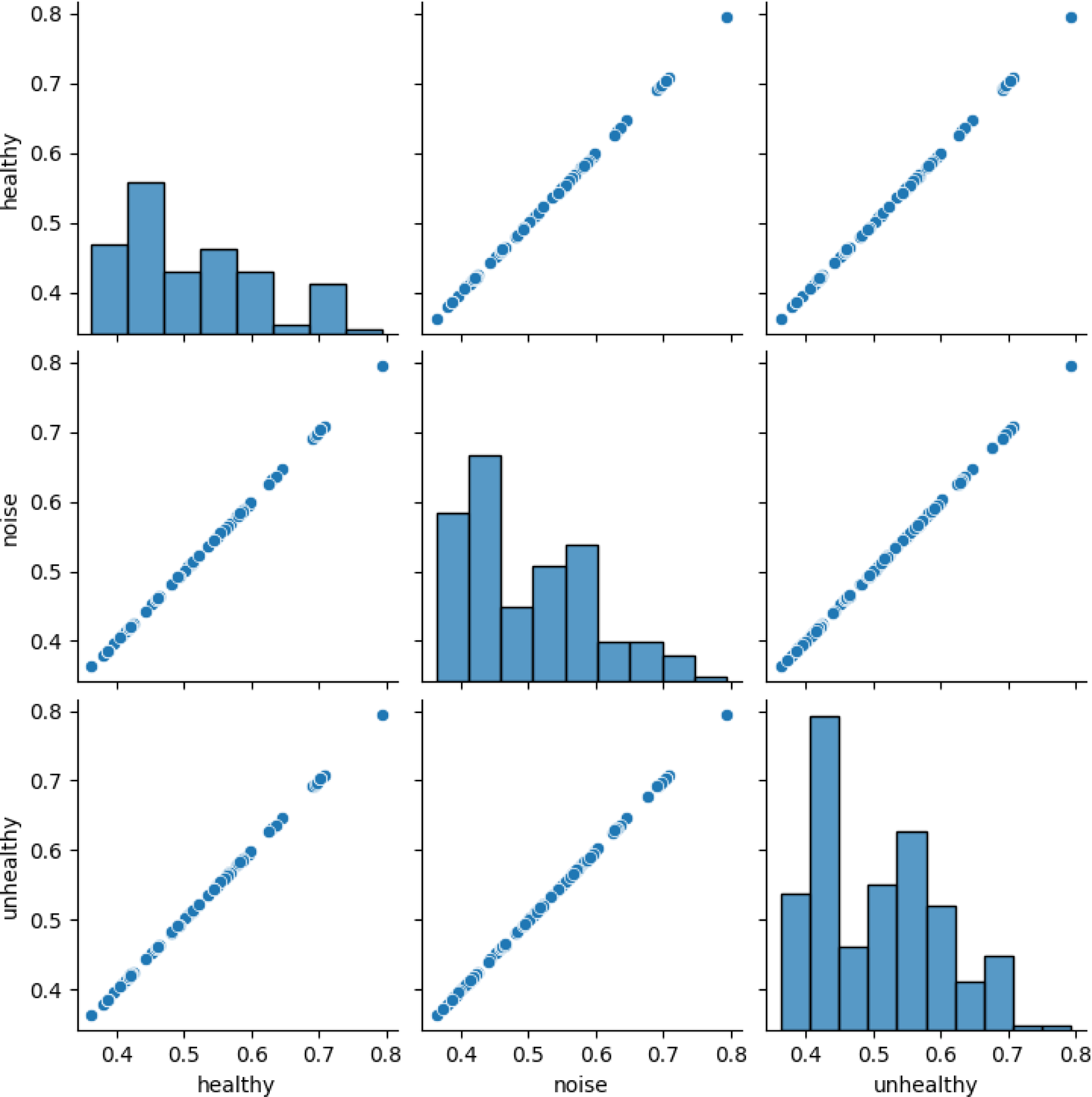
Sentiment Correlation in Chicken Vocalizations Across Various Phases. This graph details the correlation, or lack thereof, between sentiment scores across three distinct phases - healthy, noise, and unhealthy, illustrating how sentiments align or diverge under different environmental or health conditions.

##### 6.2.6.2. Emotion Distribution by Category

In the “Healthy” dataset, the emotion distribution predominantly skews towards neutrality, with almost 90% of the emotions categorized as neutral, interspersed with minimal negative sentiments and occasional joyful expressions. This suggests a baseline emotional content (Figure 17B) in typical, undisturbed chicken vocalizations.

**Figure 17B.**
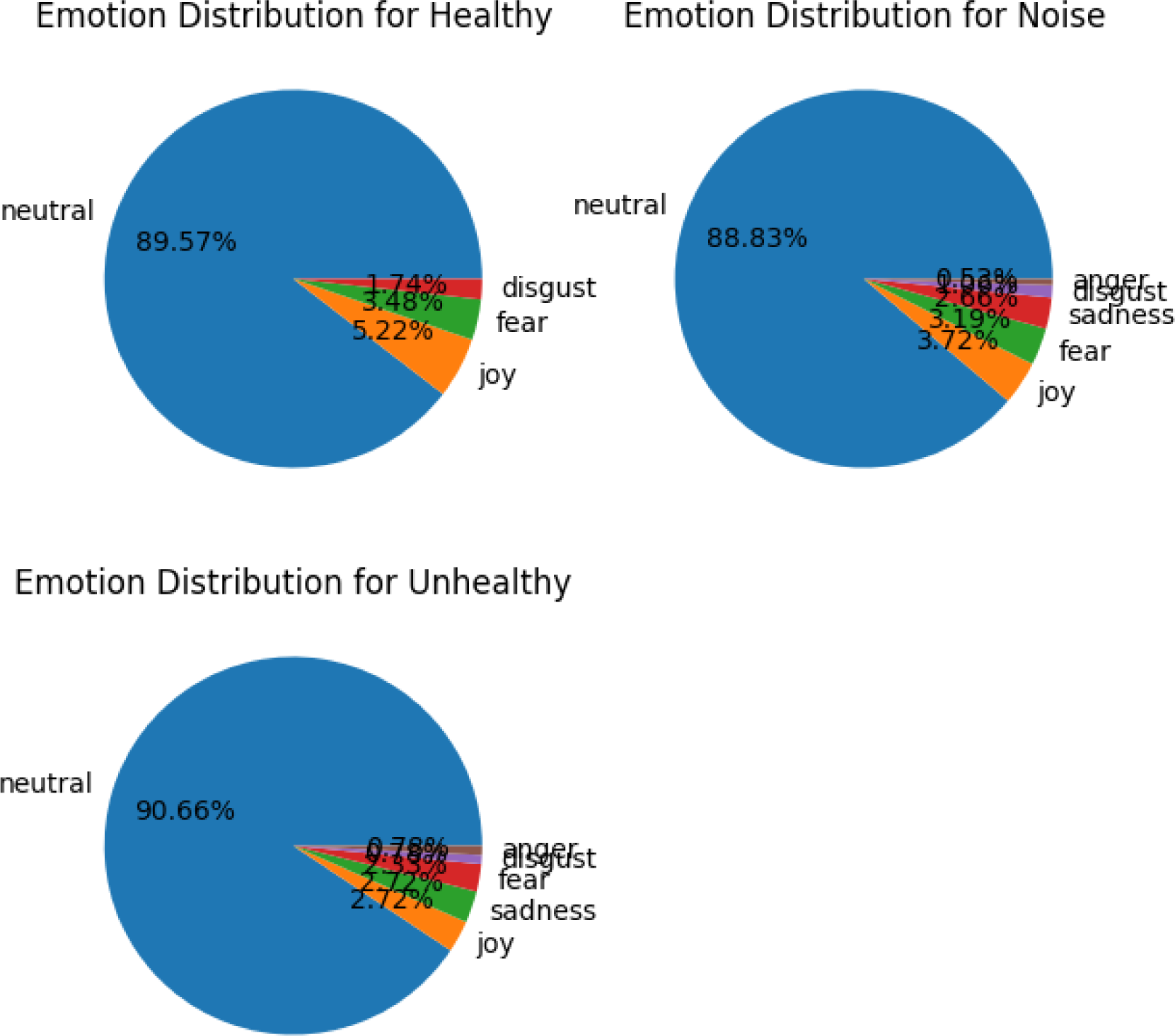
Emotional Profiles of Laying Hens Across Different Phases. This series of pie charts delineates the distribution of emotions—neutral, fear, sadness, and joy—in chickens during healthy, noise, and unhealthy phases. It provides a visual summary of emotional shifts and the prevalence of specific emotions in response to environmental noise and health challenges.

Transitioning to the “Noise” dataset, while the emotional tone remains largely neutral at approximately 90%, there is a notable presence of both negative and positive emotions, with around 4% of sentences exhibiting joy and 3% showing signs of fear. This indicates the impact of environmental noise on the emotional expression of chickens, introducing slight stress or disturbance. In the “Unhealthy” dataset, the emotional tone shifts slightly towards positivity, with around 90% positivity which includes a small but significant presence of negative emotions alongside traces of joy and sadness, each around 3%. This could reflect varying degrees of discomfort or stress due to health issues yet a predominance of baseline neutral to positive expressions.

Across all conditions—healthy, noise, and unhealthy—the overarching pattern remains one of neutrality with nuances of positive and negative emotions reflecting the complex interplay of health and environmental factors on chicken vocalizations.

##### 6.2.6.3. Emotion Score Analysis

Analysis of emotion scores across the healthy, noise, and unhealthy datasets reveals distinct patterns. In the healthy dataset, emotion scores range from 0.37 to 1.0, predominantly clustering between 0.8 and 1.0, indicating intense emotional expressions typically associated with stable, undisturbed conditions (Figure 17C). The noise dataset shows a wide distribution of scores from 0 to 1.0, with a concentration around 1.0, suggesting strong emotional responses possibly triggered by environmental stressors. In the unhealthy dataset, scores vary from 0 to 1.0, with peaks at 0.6, 0.8, and 1.0, highlighting emotional fluctuations possibly due to health-related distress.

**Figure 17C.**
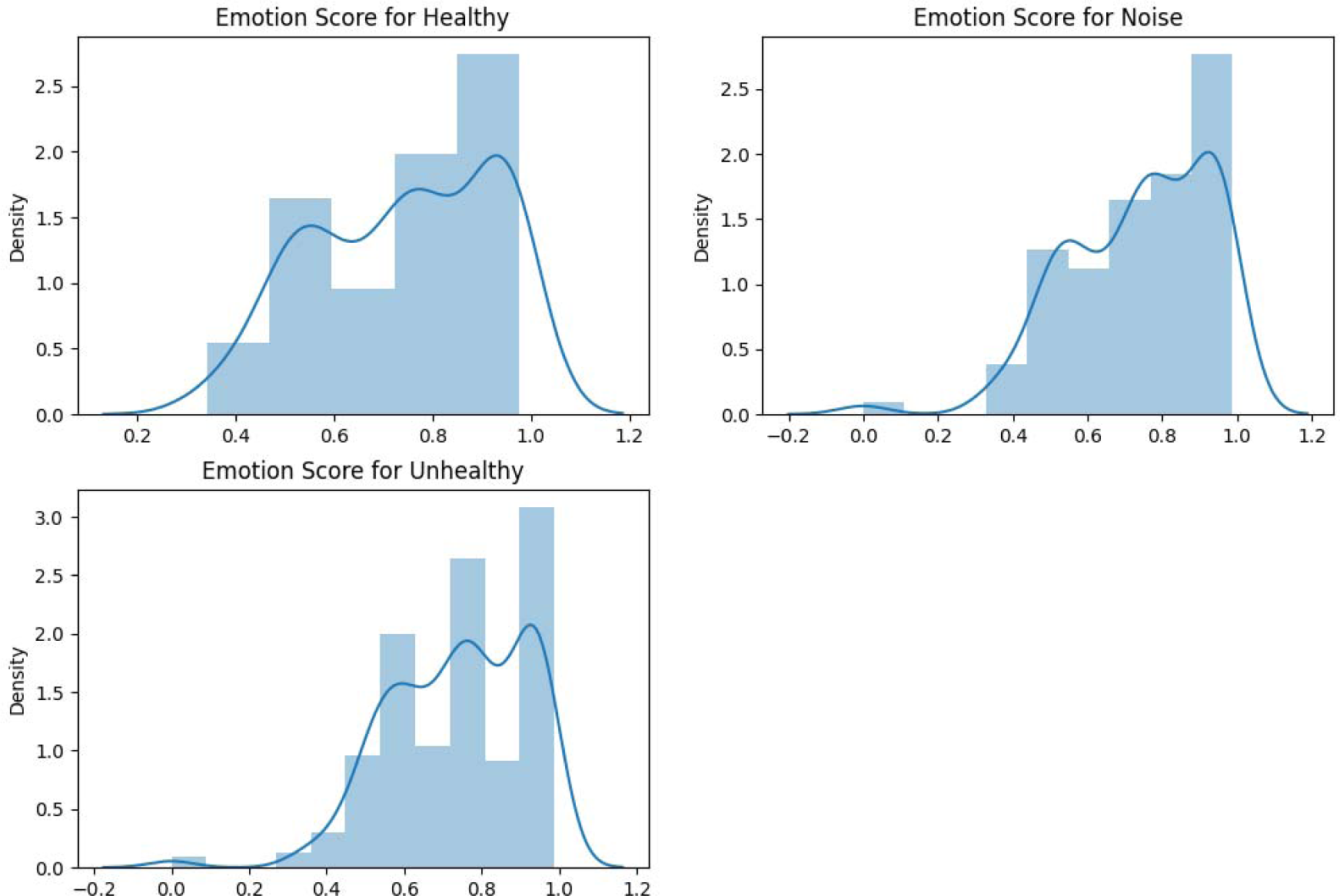
Emotion Score Density Distribution in Chickens Across Health Phases. This Displot presents the frequency distribution of emotion scores, highlighting variations in emotional intensity experienced by chickens during healthy, noise, and unhealthy conditions, with scores ranging from 0.5 to 1.0.

##### 6.2.6.4. Box Plot Analysis of Emotion Scores

In the healthy dataset, the box plot (Figure 17D) spans from approximately 0.10 to 1.0, with a median around 0.8, suggesting strong emotions dominate. This distribution is consistent in the noise dataset, with a median also at 0.8 but a slightly wider spread from 0.39 to nearly 1.0, reflecting the variability introduced by noise. The unhealthy dataset shows a distribution starting from 0.421 to 0.8, with a similar median, indicating that while the emotional intensity is comparable, the range is slightly narrower, possibly reflecting a different emotional response to health stressors.

**Figure 17D.**
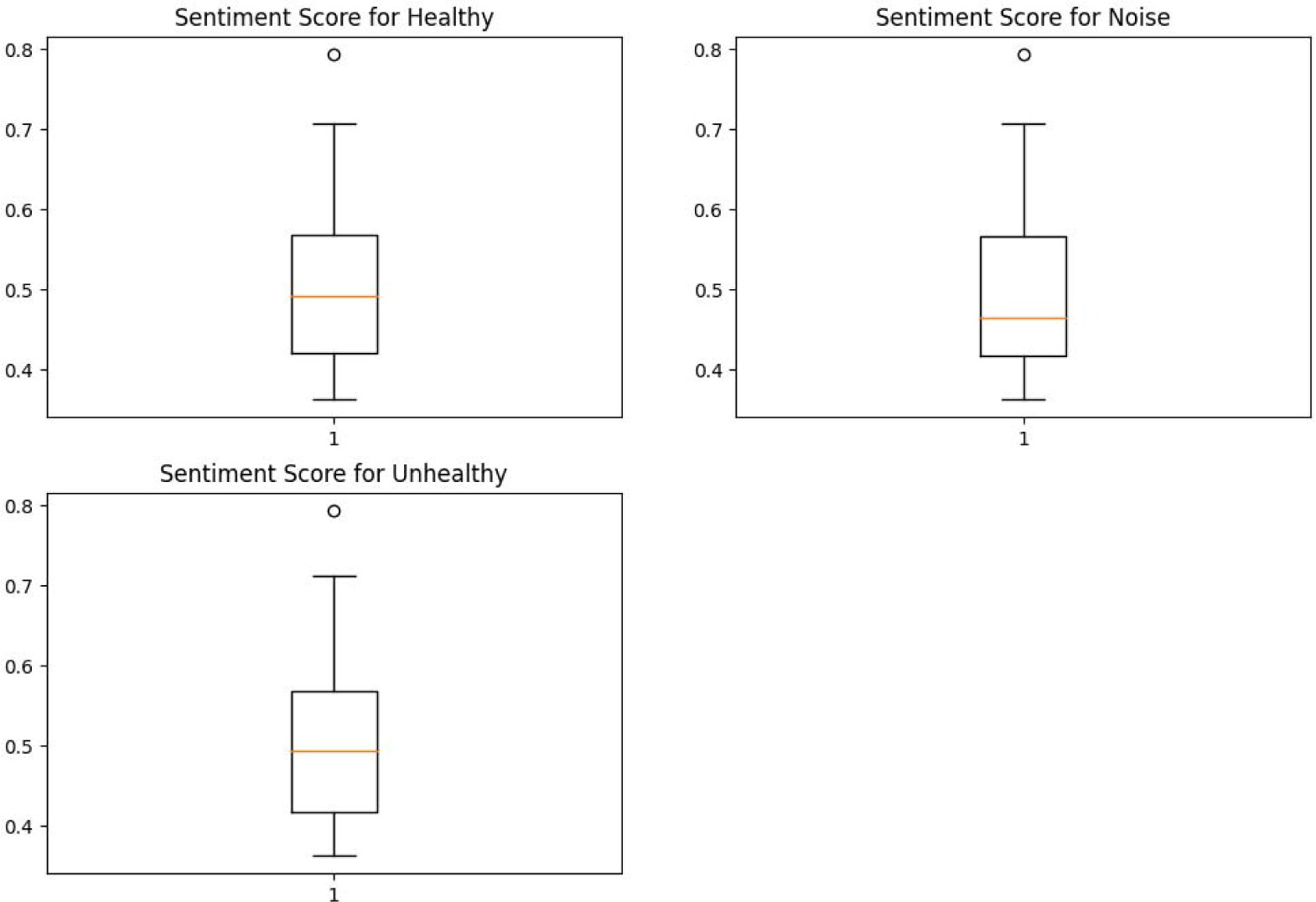
Boxplot Analysis of Emotion Scores Across Chicken Health Phases. This boxplot illustrates the range and distribution of emotion scores from 0.5 to 1.0 among chickens during healthy, noisy, and unhealthy phases. The plot emphasizes median values, interquartile ranges, and outliers, providing insights into the intensity of emotional expressions under different conditions.

In our analysis, a significant correlation in emotion scores across all datasets was observed, suggesting that emotional responses are consistently mirrored across different conditions. This phenomenon indicates that when emotion scores increase in one dataset, they tend to increase correspondingly in the others. Notably, the correlation coefficients start from approximately 0.2, underlining a measurable positive relationship (Figure 17E).

**Figure 17E.**
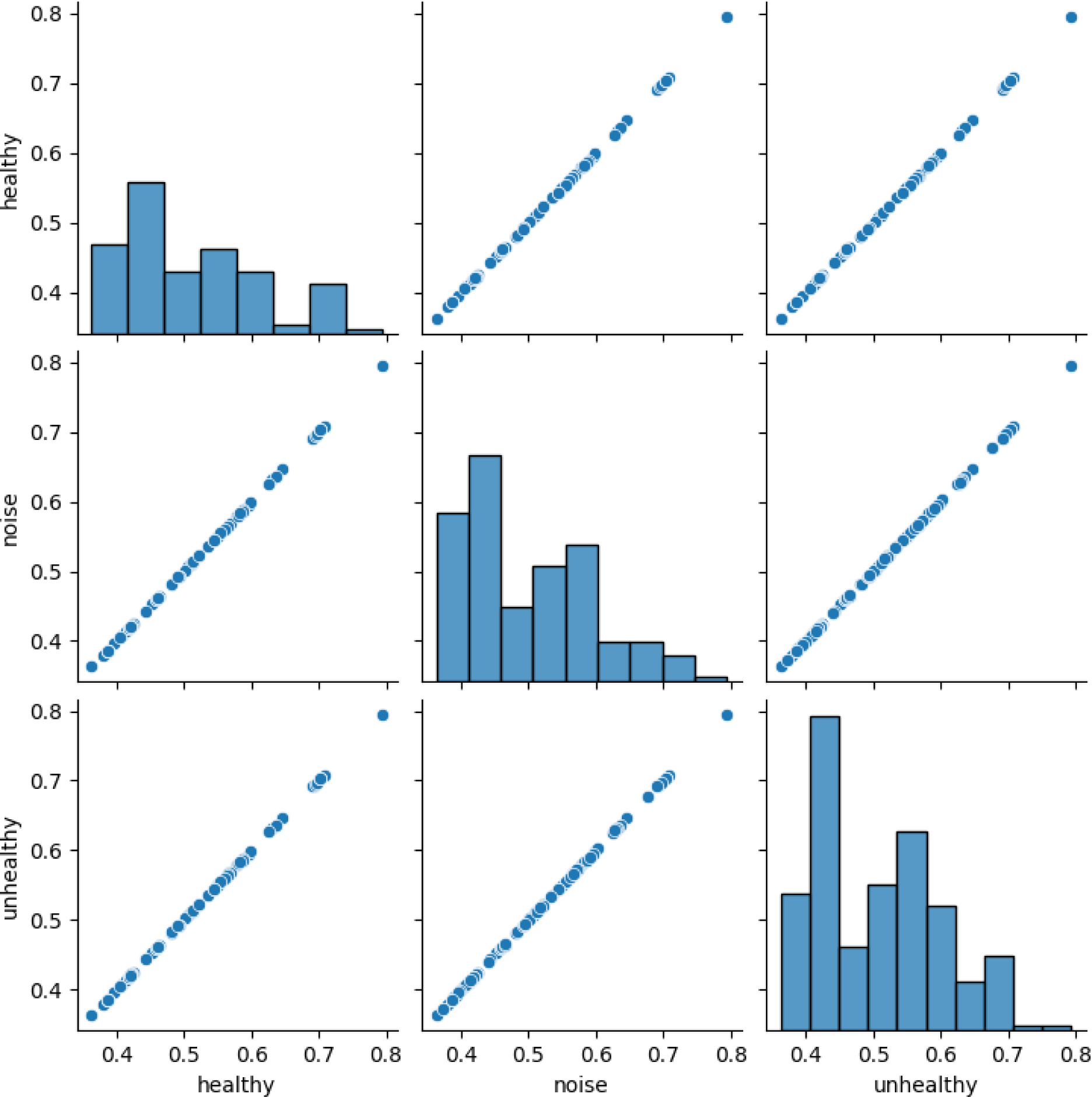
Correlation Matrix of Emotion Scores Across Chicken Vocalization Phases. This figure illustrates the correlation between emotion scores across the healthy, noise, and unhealthy datasets.

**Figure 18.**
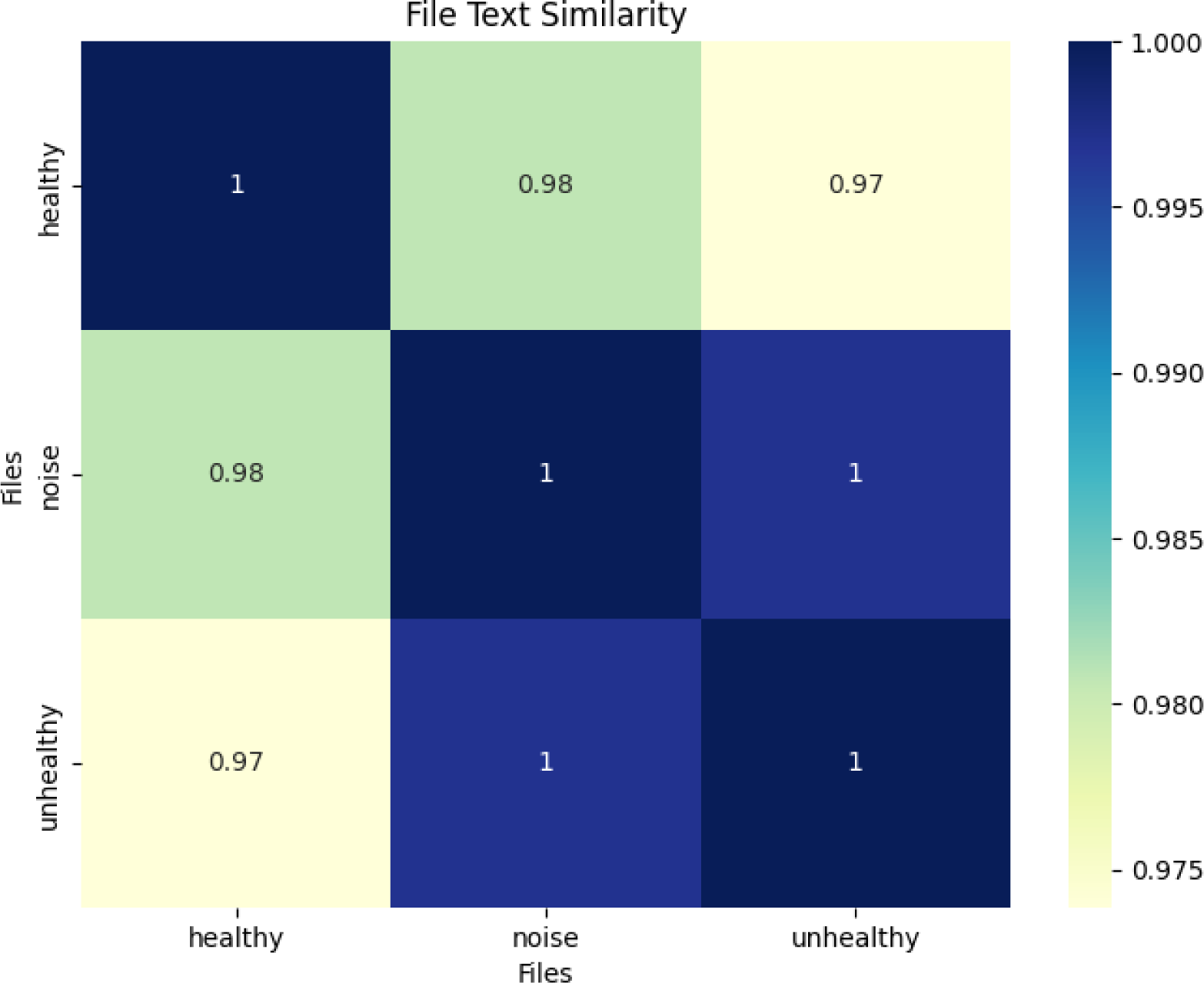
Cosine Similarity Matrix Map illustrating textual similarities among the ‘healthy’, ‘noise’, and ‘unhealthy’ datasets. The Y-axis lists the files being compared, and the X-axis corresponds to these files, providing a detailed visual analysis of the degrees of textual similarity across different stages—healthy, noise, and unhealthy. The matrix quantifies these relationships using cosine similarity metrics, highlighting significant overlaps in content and structure.

Among the dataset comparisons, the correlation between the healthy and unhealthy datasets, as well as between the healthy and noise datasets, was particularly robust. These strong correlations might reflect underlying similarities in emotional responses of chickens under varying environmental conditions or health states. The graphical representation of these correlations shows a remarkably strong alignment, especially between the noise and unhealthy datasets, which displayed one of the strongest correlations, suggesting similar emotional patterns despite the ostensibly different environmental conditions.

Furthermore, the relationship between the noise and healthy datasets depicted a perfect horizontal line in the correlation graph, highlighting a uniquely consistent and robust relationship across these conditions. Although this correlation appeared slightly weaker when compared to the other dataset pairs, it still represented a significant association. This consistency across different phases implies that there are common emotional responses triggered by both background noise and general health conditions in chickens. The results reveal a distinct pattern of emotional correlation across the datasets, with varying strengths of correlation suggesting that while the conditions differ, the emotional responses exhibit certain universal traits. This consistent emotional correlation underscores the interconnectedness of emotional responses in chickens across different health and environmental scenarios, which could have implications for understanding stress and well-being in poultry environments.

#### 6.2.7. Text File Similarity Analysis

In this study, we employed a graph-based method to evaluate the extent of textual similarity among the datasets. The analysis revealed remarkably high similarity scores, indicating significant overlap in content across different conditions. Specifically, the similarity between the “healthy” and “noise” datasets was scored at an impressive 0.98. Additionally, the “unhealthy” dataset demonstrated nearly identical content with both the “healthy” and “noise” datasets, achieving similarity scores of 0.97 respectively.

Most notably, the “noise” dataset exhibited complete congruence (100%) with the “unhealthy dataset, and a very high similarity score of 98% with the “healthy” dataset. This perfect score indicates that the texts from the “noise” and “unhealthy” conditions are virtually indistinguishable in terms of their content and structure. Similarly, the “unhealthy” dataset showed complete similarity (100%) with both the “noise” and “healthy” datasets, further emphasizing the significant textual overlap.

These findings highlight a profound level of textual similarity across the datasets, suggesting that the content, likely including descriptions of environmental and health conditions, is highly consistent across different files. This high degree of similarity may reflect standardized recording and notation practices or common thematic elements prevalent across the different environmental and health conditions of the datasets.

#### 6.2.8. Frequency Analysis Results

In the “healthy” dataset, the term “going” is notably the most frequent, appearing over 2000 times. “Nihai” follows with approximately 1500 instances (Figure 19), reflecting its significance within the dataset. Additional terms such as “room,” “next,” “back,” “other,” and “side” also show high frequency, each surpassing 1000 mentions. This prevalence underscores their importance in the context of chicken vocalizations under healthy conditions.

**Figure 19.**
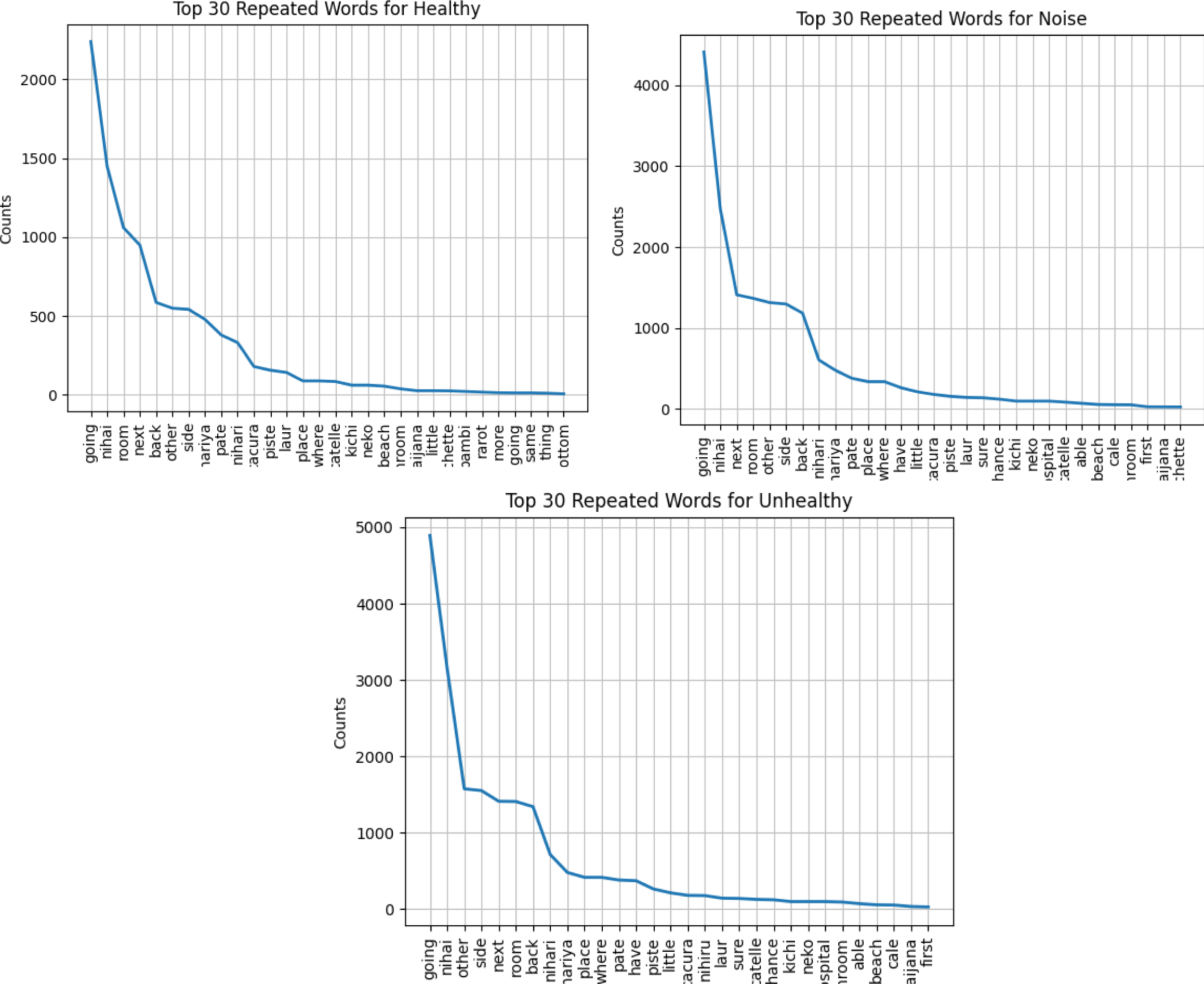
Comparative Analysis of the Top 30 Most Frequently Occurring Words Across Three Phases: Healthy, Noise, and Unhealthy. This figure presents a side-by-side comparison of the counts (Y-axis) of the top 30 words from each phase (X-axis) in chicken vocalizations. It provides a detailed view of lexical trends and consistencies observed through the cycles of health and environmental noise, highlighting how certain words persist or vary across different conditions.

**Figure 20A.**
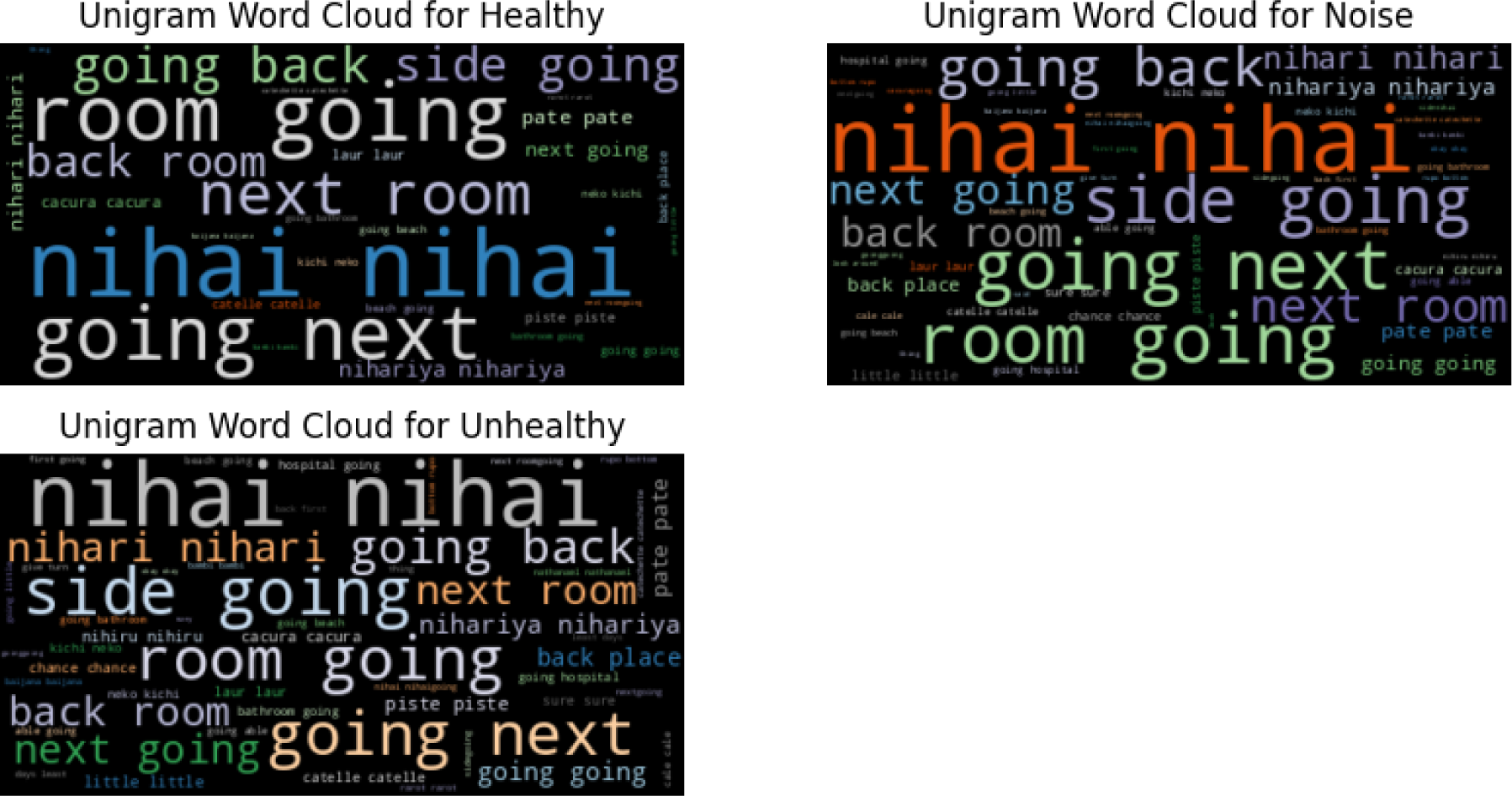
Unigram Word Cloud for Chicken Vocalizations Across Phases – This word cloud illustrate the most common unigrams found in chicken vocalizations during healthy, noise, and unhealthy phases. It highlights key terms like “going” and “nihai” and their consistent presence across various contexts, providing a visual overview of word usage patterns in different environmental conditions.

**Figure 20B.**
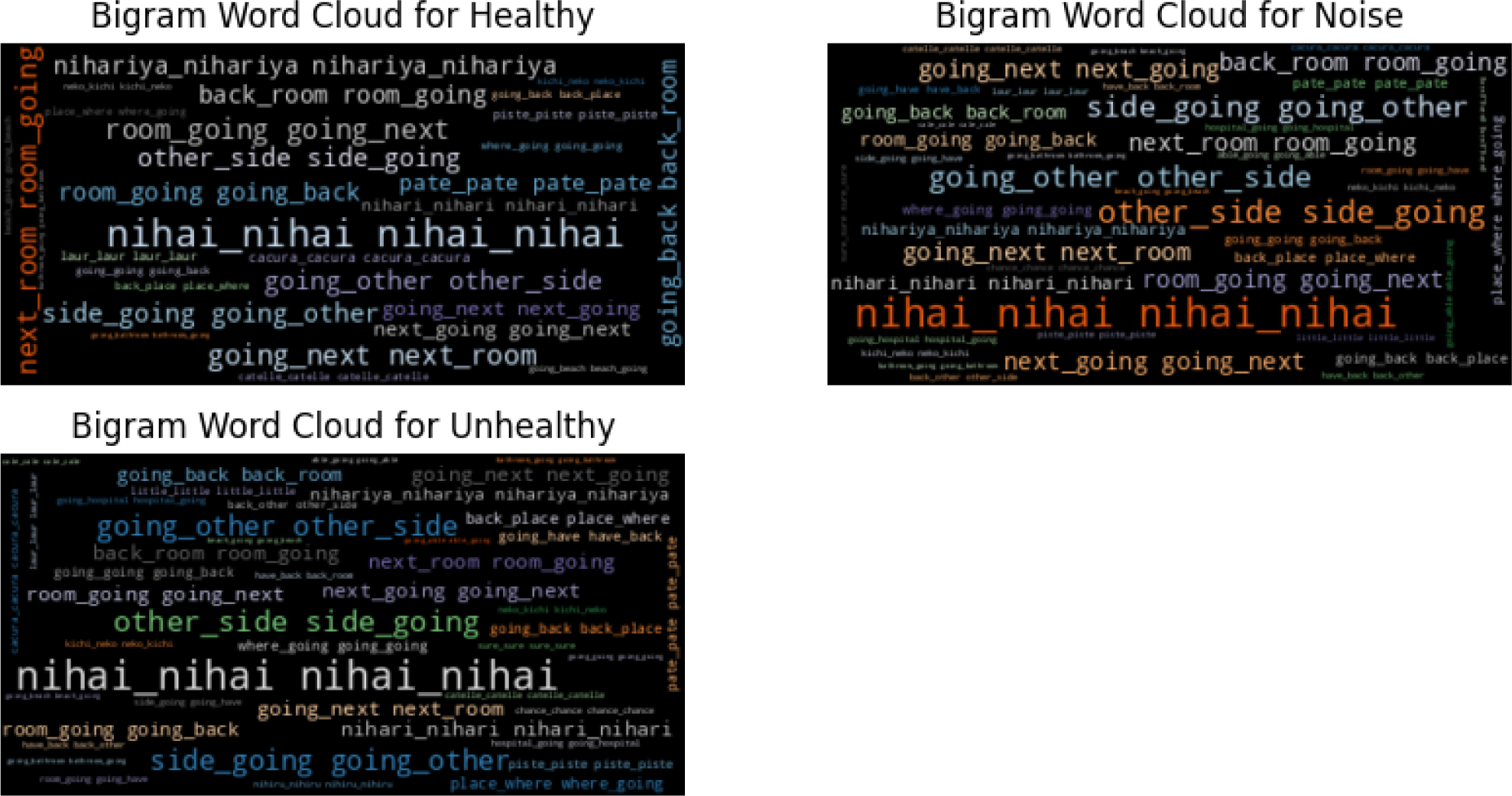
Bigram Word Cloud for Chicken Vocalizations Across Conditions – This word cloud visualizes the analysis of bigram frequency, revealing the main word pairs that dominate discussions within chicken vocalizations across healthy, noise, and unhealthy conditions. It illustrates significant linguistic pairings, showcasing how these terms interconnect to form the core thematic elements within each vocalization scenario.

**Figure 20C.**
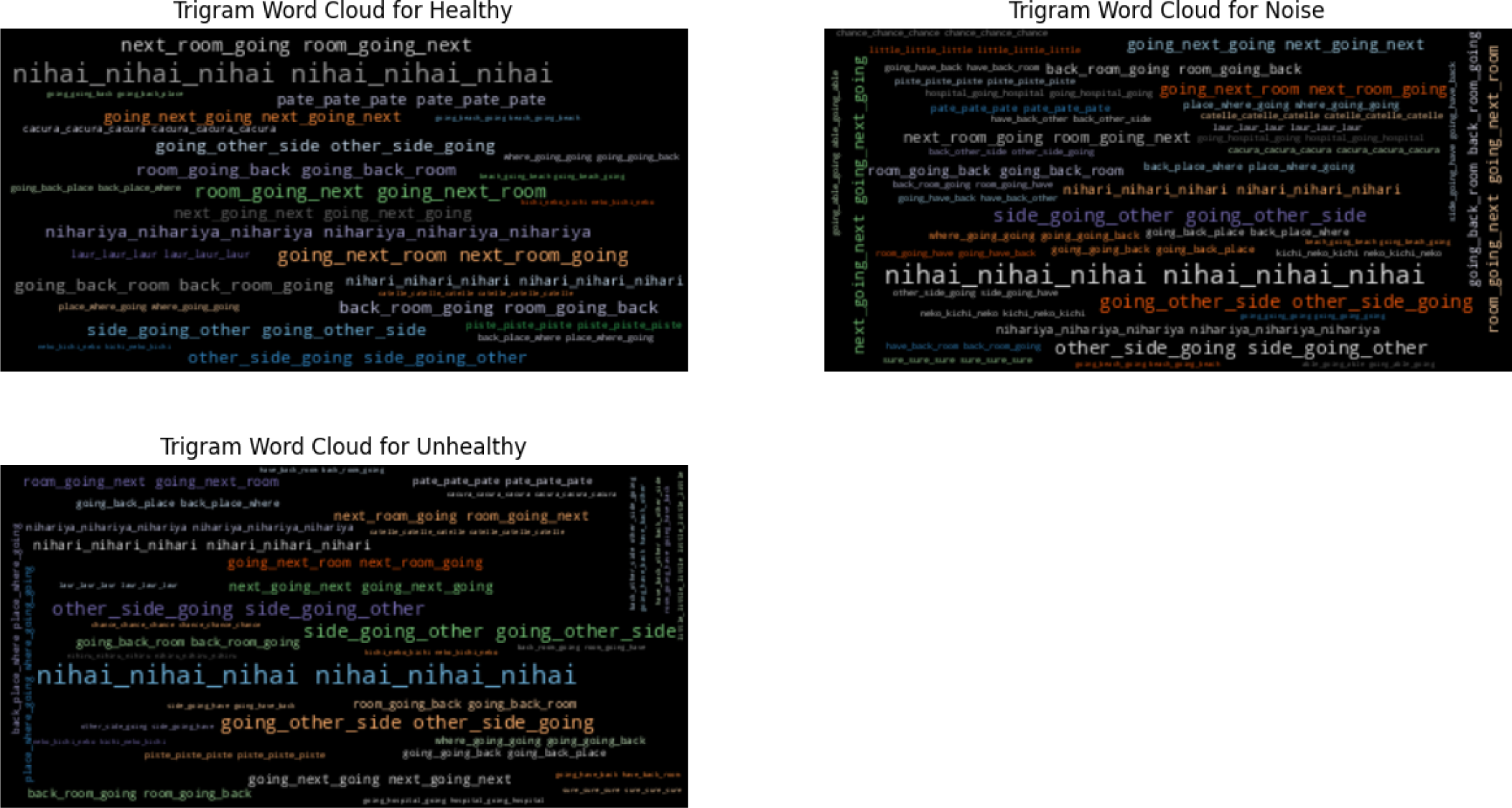
Trigram Word Cloud for Chicken Vocalizations Across Stages – This visualization offers an in-depth look at the trigram frequencies in chicken vocalizations, spanning healthy, noise, and unhealthy stages. It highlights the complex linguistic structures and thematic consistency across the datasets, emphasizing the dominant trigrams that recur and shape the narrative in each phase of vocalization.

**Figures 21 A-C.**
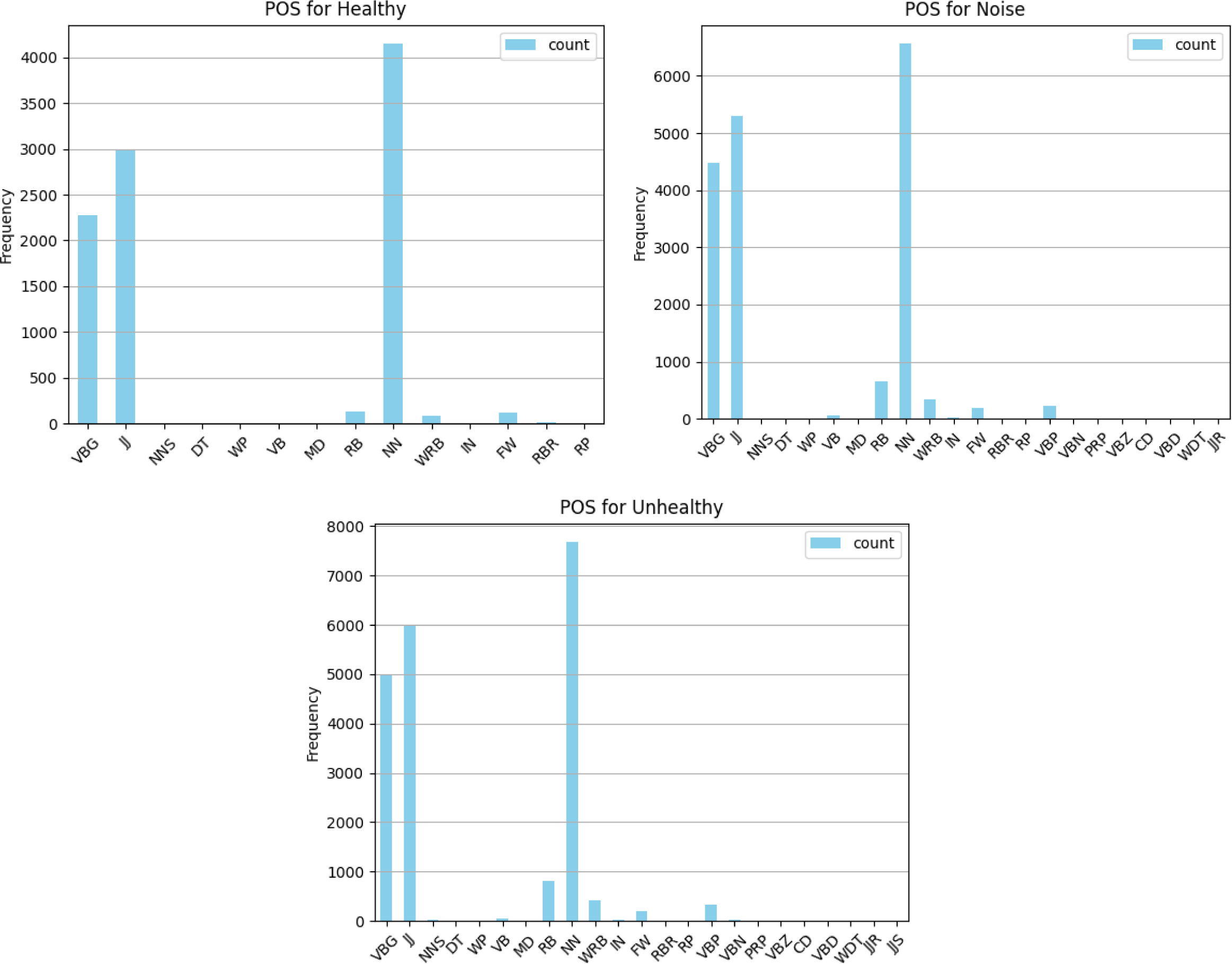
Comparative Distribution of Parts of Speech in Chicken Vocalizations Across Healthy, Noise, and Unhealthy Phases. These graphs demonstrate the frequency distribution of parts of speech for the Healthy (A), Noise (B), and Unhealthy (C) phases of chicken vocalizations, as analyzed through th Whisper model. Each graph distinctly highlights the variations in linguistic composition, showcasing how different health conditions affect the usage of various grammatical categories in chicken vocalizations.

Transitioning to the “noise” dataset, a consistent lexical pattern emerges. The word “going” dominates once more, with over 4000 occurrences, and “nihai” maintains substantial prominence with around 2500 mentions. Other significant terms include “next,” “room,” “other,” “side,” and “back,” each exceeding 1000 occurrences. This consistency in word repetition between the “healthy” and “noise” datasets suggests a uniformity in vocabulary usage despite varied acoustic backgrounds.

Within the “unhealthy” dataset, the persistence of the word “going,” which appears over 5000 times, illustrates its ongoing relevance. “Nihai” also remains highly recurrent, along with “other,” “side,” “next,” “room,” and “back,” all surpassing 1000 instances. This repeated word usage across the datasets underscores a common linguistic framework that spans across different health states of the chickens.

Our comprehensive analysis reveals a notable uniformity in word usage across the “healthy,” “noise,” and “unhealthy” datasets, indicating a shared linguistic pattern that persists irrespective of the health condition or environmental context of the chickens. This suggests that specific terms are central to discussing and analyzing chicken vocalizations, reflecting consistent thematic elements within the datasets.

#### 6.2.9. Unigram Analysis of Chicken Vocalizations

In the “healthy” dataset, the word “going” emerges as the most frequent, appearing over 2000 times. It is closely followed by “nihai,” which occurs around 1500 times. Other frequently occurring words include “room,” “next,” “back,” “other,” and “side,” each surpassing 1000 repetitions. This highlights their significant presence within the text, indicating common themes or activities discussed in this phase.

The “noise” dataset shows a similar pattern to the “healthy” dataset, with “going” leading again, noted over 4000 times. “Nihai” follows with approximately 2500 occurrences. The consistency in word usage with words such as “next,” “room,” “other,” “side,” and “back” being prominent in both datasets suggests a uniformity across different environmental contexts, emphasizing the resilience of certain lexical items in varied settings. In the “unhealthy” dataset, “going” remains the most repeated word, appearing over 5000 times. “Nihai” also maintains a high frequency of occurrences. This dataset continues to share common words with the other two, like “other,” “side,” “next,” “room,” and “back,” each appearing over 1000 times. This repetition across all conditions underlines a shared pattern in textual content, pointing to similar themes or topics of discussion regardless of the health status of the chickens. The analysis across the “healthy,” “noise,” and “unhealthy” files reveals a notable similarity in word usage, suggesting a commonality in language patterns despite differences in the file context. This demonstrates that certain words are central to describing chicken vocalizations, reflecting consistent elements of their behavior or environment.

#### 6.2.10. Bigram Frequency Analysis

The most frequent bigram in the “healthy” dataset is (’nihai’, ‘nihai’), with 1438 instances, indicating significant emphasis on this term. Other notable bigrams include (’room’, ‘going’) and (’going’, ‘next’), which highlight common narrative threads about movement or sequential actions within a room or environment. In the “noise” dataset, the bigram (’nihai’, ‘nihai’) again appears prominently, with repeated occurrences suggesting its pivotal role in discussions within noisy environments. Other frequent bigrams such as (’room’, ‘going’) and (’going’, ‘next’) indicate similar themes of movement and spatial relations as seen in the “healthy” dataset. Th “unhealthy” dataset shows a high recurrence of the bigram (’nihai’, ‘nihai’), indicating its central thematic importance. Spatial and action-oriented bigrams like (’other’, ‘side’) and (’side’, ‘going’) are also significant, reflecting discussions on spatial dynamics or contrasting elements within the discussions of unhealthy conditions.

#### 6.2.11. Trigram Frequency Analysis

The dominant trigram (’nihai’, ‘nihai’, ‘nihai’) in the “healthy” dataset underscores a strong focus on the term “nihai.” Other common trigrams involve sequences of movement or location transitions, such as (’next’, ‘room’, ‘going’), reflecting detailed actions within specific settings. Similar to the “healthy” dataset, the “noise” dataset frequently features the trigram (’nihai’, ‘nihai’,

’nihai’), highlighting its thematic relevance. Trigrams like (’other’, ‘side’, ‘going’) and (’going’, ‘next’, ‘going’) suggest a continuity in discussing movement and alternative options or paths within noisy environments. In the “unhealthy” dataset, the prevalence of (’nihai’, ‘nihai’, ‘nihai’) continues, paired with trigrams that emphasize back-and-forth movements or alternative routes, such as (’back’, ‘room’, ‘going’), which may reflect the complexity of scenarios discussed in the context of illness or stress. Across all three datasets, the consistency of certain trigrams, especially involving the term ‘nihai,’ indicates its significant role in the vocalizations across different health and environmental contexts. This repetition across datasets suggests common underlying themes or discussions prevalent in the vocalization studies of chickens, providing insights into their behavioral expressions.

#### 6.2.12. Parts of Speech Analysis Across Different Health States of Chicken Vocalizations

The linguistic profile of the “healthy” dataset provides a detailed view of the prevalent grammatical structures. Verbs in gerund or present participle forms (VBG) are notably dominant, tallying at 2,278 occurrences, which illustrates a narrative rich in descriptions of ongoing actions. Adjectives (JJ) are also significantly utilized, with 2,978 appearances, emphasizing the descriptive nature of the text. Singular or mass nouns (NN) top the frequency with 4,148 mentions, underscoring the text’s focus on specific entities or ideas. Adverbs (RB) and foreign words (FW) appear 127 and 122 times, respectively, adding depth and diversity to the discourse. Additionally, WH-adverbs (WRB) and comparative adverbs (RBR) surface less frequently, suggesting limited usage of complex comparative or interrogative constructs. The sparse appearance of modal verbs (MD) and base form verbs (VB) further points to a narrative that prefers descriptive and continuous tense forms over expressions of possibility or direct actions.

In the “noise” dataset, adjectives (JJ) prominently appear 5,298 times, indicating a strong focus on descriptive elements within the vocalizations. The count of singular or mass nouns (NN) reaches 6,564, reflecting frequent references to tangible and intangible elements. The verbs in gerund or present participle form (VBG) follow closely with 4,480 instances, suggesting continuous or repeated actions within the dataset’s context. Adverbs (RB) are also prevalent at 652 occurrences, providing detailed modifications to actions and qualities. The presence of WH-adverbs (WRB) and foreign words (FW) at 337 and 193 occurrences respectively, enriches the text with questions and multicultural or technical terms. The lower frequency of modal verbs (MD), determiners (DT), and prepositions (IN) highlights a narrative less concerned with hypotheticals and more focused on describing ongoing states or actions.

The “unhealthy” dataset reveals a complex linguistic structure where adjectives (JJ) are most frequent at 6,005 occurrences, emphasizing the strong descriptive focus of the narrative. This is followed by singular or mass nouns (NN) at 7,675 occurrences, which dominate the discourse, indicating a significant emphasis on various entities and states. Verbs in the gerund or present participle form (VBG) are also highly present at 4,974 instances, portraying ongoing conditions or actions. Adverbs (RB) and WH-adverbs (WRB) add layers of detail and questioning to the text, occurring 809 and 414 times respectively. The analysis shows minimal use of modal verbs (MD), determiners (DT), and prepositions (IN), which suggests a narrative style that leans towards detailed descriptions and continuous actions rather than conditional or relational frameworks. The incorporation of foreign words (FW) highlights the inclusion of diverse linguistic elements, possibly reflecting specific technical or cultural contexts.

#### 6.2.13. Topic Modeling Analysis of Chicken Vocalization Data

In analyzing chicken vocalizations, the topics extracted across different datasets—healthy, noise, and unhealthy—reveal a broad spectrum of settings, activities, and themes, enriching our understanding of vocal behavior in varying environments.

In the “healthy” dataset, the diversity of topics (Figure S6) ranges from outdoor activities such as recreational or leisure pursuits (Topic 0: “back,” “piste,” “bathroom”) to indoor environments indicating domestic spaces (Topic 1: “room,” “bathroom”). Discussions also extend to specific terms or entities frequently mentioned (Topic 2: “nihai”), leisure destinations (Topic 3: “beach,” “room”), and cultural or culinary concepts (Topic 8: “nihari,” “nihariya”). The topics suggest an extensive range of activities, from discussing specific locations or objects to ongoing actions and transitions between activities, showcasing the complexity of the healthy vocalization context.

The “noise” dataset topics delve into diverse themes from various locations or environments (Topic 0: “place,” “piste,” “bathroom”) to attributes of minor matters (Topic 1: “little,” “first”). The discussions also cover physical movements or actions (Topic 2: “back,” “laur”), spatial arrangements (Topic 3: “room,” “sidegoing”), and specific institutions or establishments (Topic 4: “side,” “hospital”). This dataset (Figure S7) illustrates the chickens’ vocal response to potentially disruptive or noisy environments, with a focus on actions or behaviors and certainty or assurances about ongoing activities (Topic 8: “nihari,” “sure”).

The “unhealthy” dataset (Figure S8) provides insights into topics concerning medical settings and physical spaces (Topic 0: “piste,” “bathroom,” “hospital”), specific actions or attributes (Topic 1: “back,” “little”), and discussions about certainty or quantities (Topic 6: “sure,” “days”). These topics highlight the range of discussions from health-related environments to future possibilities or actions, reflecting the conditions and concerns in scenarios of ill health.

#### 6.2.14. Analysis of Inter-Topic Distances Across Datasets

In the healthy dataset, topics 1, 4, and 8 form a distinctive triangular shape, indicating their dissimilarity from the other seventeen topics. These three topics are uniquely positioned in the topic space, suggesting specific themes that do not overlap much with others. In contrast, the remaining seventeen topics are closely clustered, suggesting a high degree of overlap and similarity in the themes they cover.

In the noise dataset, topic 1 is noticeably isolated, standing apart from topics 2 and 3, which are nearer to each other but still distinct. The rest of the topics in this dataset form a dense cluster, indicating a significant overlap in thematic content, which suggests that most topics within the noise dataset are closely related or share similar themes.

For the unhealthy dataset, topics 3, 4, and 8 are distinctly separated from the rest, marking them as unique in the context of this dataset. The other topics, while closer to each other, show a certain level of thematic interdependence yet maintain clear distinctions, illustrating a complex interplay of similar yet distinguishable themes.

Across all datasets—healthy, noise, and unhealthy (Figure S9)—the analysis of inter-topic distances reveals significant insights into how topics are spatially and thematically arranged. In the healthy dataset, the clear distinction among three topics against a backdrop of closely related others suggests specialized discussions that are not as prevalent in the rest of the data. In the noise and unhealthy datasets, the isolation of specific topics indicates unique discussions that are particularly relevant to the conditions simulated in these environments. These spatial relationships highlight the distinct thematic structures within each dataset, enhancing our understanding of the underlying patterns in chicken vocalizations across different phases.

## 7. Discussion

Table 3 presents an exhaustive sentiment analysis of poultry vocalizations across diverse experimental settings, including control and treatment groups under stress (dataset 1), and observations in healthy, unhealthy, and noise conditions (dataset 2). These metrics are pivotal for delineating the psychological and emotional responses of poultry to varied stressors and environmental stimuli, offering critical insights for advancing animal welfare practices. By comparing sentiment distributions and scores across different stimuli, the study elucidates the complex relationship between environmental factors and poultry welfare.

**Table 3.**
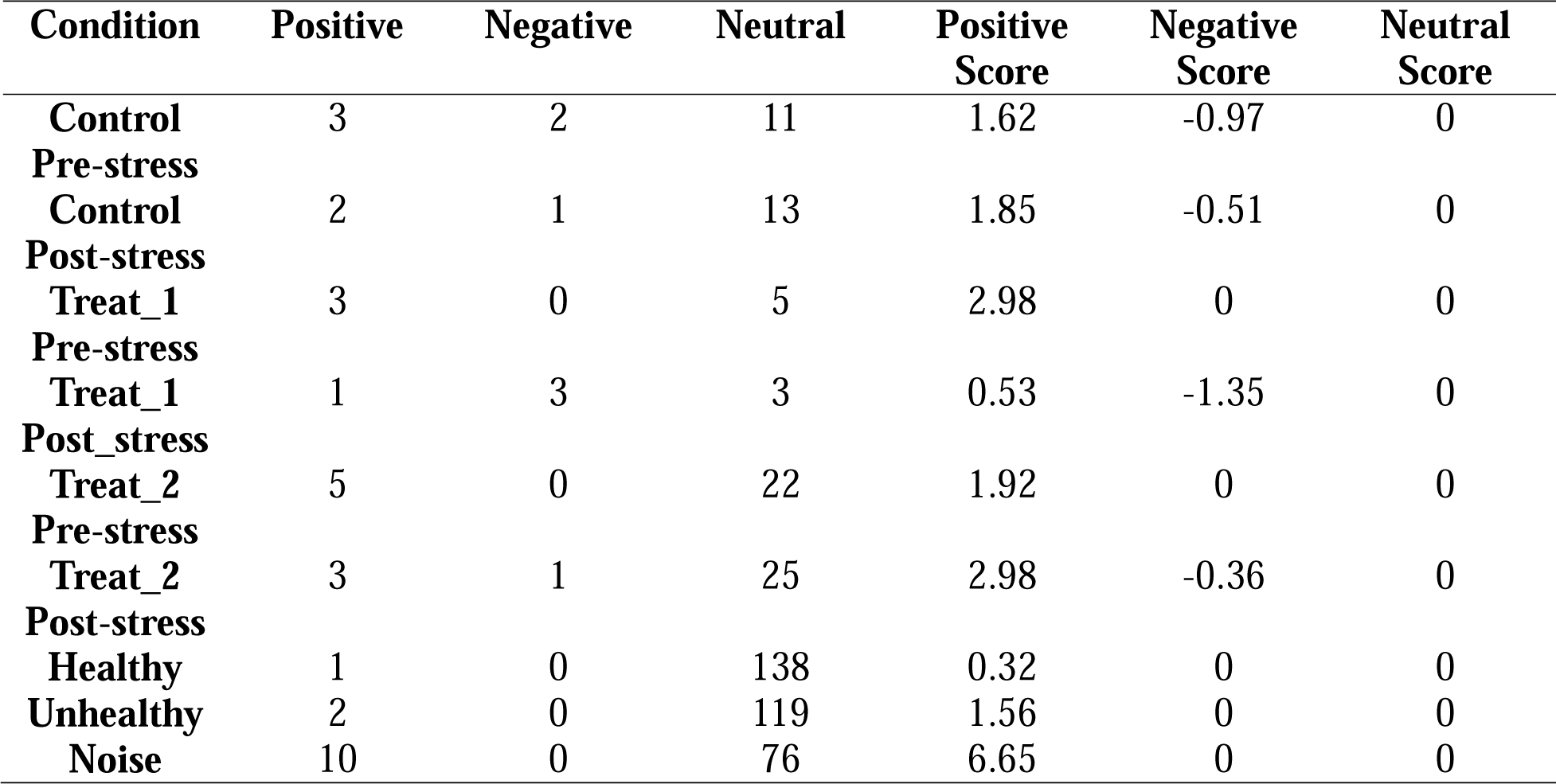
Comparative Sentiment Analysis of Poultry Vocalizations Under Varied Stress and Environmental Conditions.

The sentiment analysis reveals significant affective reactions of chickens to diverse stimuli, employing classifications of positive, negative, and neutral sentiments with corresponding scores, thereby innovating poultry welfare evaluation through vocal expression analysis. A discernible shift in emotional expression was observed within the control group from pre- to post-stress conditions, marked by a decrease in positive sentiments and an increase in neutral sentiments post-stress, indicating stress’s subtle impact on welfare. Notably, the stability of negative sentiments suggests that the applied stressor did not elicit significant distress signals.

Treatment groups responded differently to stress induction. Treatment 1 experienced a decline in positive sentiments and an increase in negative sentiments post-stress, indicating a significant stress impact. In contrast, Treatment 2 saw an increase in positive sentiments post-stress, suggesting habituation or a different stressor perception. Dataset 2’s analysis under healthy, unhealthy, and noise conditions revealed baseline contentment in healthy chickens, dominated by neutral sentiments, and an intriguing increase in positive sentiments under unhealthy conditions, possibly reflecting distress communication. Noise conditions significantly increased positive sentiments, indicating enhanced vocal activity in response to environmental stimuli, potentially for social communication.

The study highlights the complexity of interpreting poultry vocalizations and the varied impacts of environmental and health stressors on welfare. The findings suggest that chickens may increase social vocalizations in response to noise, implying social dynamics within poultry farms and vocalization patterns as potential environmental enrichment or stress indicators. This sentiment analysis underscores the nuanced context required in welfare assessments, emphasizing qualitative stressor impacts on poultry vocal expressions. It showcases the potential of NLP and machine learning in poultry welfare research, offering a novel approach to understanding animal subjective experiences, complementing traditional welfare assessments, and promoting humane poultry management practices.

This research introduces a pioneering method to interpret the emotional states of chickens using advanced acoustic analysis and Natural Language Processing. Unlike traditional approaches that relied on observable behaviors or invasive methods, our study employs the Whisper NLP model to extract nuanced emotional cues from vocal patterns. This approach confirms the initial hypothesis that vocalizations are not just random noises but structured communications that reflect various emotional states such as distress, contentment, and fear. These findings are crucial as they provide a basis for real-time, non-invasive monitoring systems that can significantly enhance animal welfare management.

### 7.1. Comparative Analysis with Previous Studies

Historically, research into poultry vocalizations has often focused on categorizing types of calls without a deeper analysis of their emotional content. For example, earlier studies might categorize a sound as an ‘alarm call’ but could not determine the specific nature of the threat or the level of stress it caused to the animal. In contrast, the current study leverages sophisticated machine learning algorithms to interpret subtle acoustic variations, offering a much deeper understanding of the emotional states behind the sounds. This method represents a significant advancement over studies like those conducted by Marino (2017) and Hernandez et al. (2022), which while foundational, did not utilize the power of AI to link sounds directly to specific emotional states.

### 7.2. Placing New Data within the Broader Scientific Context

This study represents a significant breakthrough in the realm of animal communications, particularly in understanding the nuances of chicken vocalizations. It extends beyond merely expanding our knowledge of poultry behavior, making substantial contributions to the field of precision animal husbandry—a domain where meticulous monitoring and tailored care protocols are paramount. By harnessing cutting-edge real-time data acquisition and analysis, this research dovetails with the innovative trends in smart farming technologies that emphasize the critical relationship between enhanced animal welfare and increased farm productivity.

Smart farming or Digital Livestock Farming, an approach characterized by its integration of advanced technologies and data-driven decisions, is rapidly transforming agricultural practices. In this context, the study’s focus on decoding chicken vocalizations through sophisticated NLP and acoustic analysis tools aligns perfectly with the smart farming ethos. These technologies not only aim to optimize the operations on poultry farms but also seek to elevate the standard of care provided to the animals. This alignment with technological innovation underscores a shift from traditional farming methods to more scientifically informed and humane approaches.

The significance of this study is further underscored by existing research, which consistently shows that improved animal welfare correlates directly with better overall health and increased productivity. For instance, chickens that are less stressed generally exhibit better growth rates, higher egg production, and lower susceptibility to diseases. By enabling farmers to understand and quickly respond to the emotional and physical distress signals of chickens, the technologies developed from this study could facilitate more proactive management of health issues, potentially reducing the incidence of disease outbreaks and lessening the need for medical interventions.

Moreover, the ability to monitor and interpret the emotional cues of chickens offers a pathway to more sustainable and ethical farming practices. Enhanced welfare standards can lead to significant improvements in farm sustainability by reducing resource wastage and minimizing the environmental impact of farming operations. For example, better welfare practices can decrease the turnover rate of livestock through improved health, which in turn reduces the resources spent on raising replacement animals and lowers the overall ecological footprint of the farm.

In practical terms, the integration of this NLP-based monitoring system into daily farming operations could revolutionize the way farmers interact with their flocks. By receiving real-time feedback on the emotional state of their chickens, farmers can make immediate adjustments to their management strategies, such as modifying feeding schedules, altering environmental conditions, or providing health interventions. This level of dynamic response not only enhances the welfare of the animals but also optimizes the productivity of the farm.

The implications of this research extend beyond individual farms, offering potential benefits to the broader agricultural sector by setting new standards for animal welfare and farm management. As these technologies become more widespread, they could help shift industry benchmarks toward more humane and scientifically grounded practices. This shift could also influence public perceptions of poultry farming, potentially increasing consumer confidence in the welfare of the animals and the quality of the products they produce.

Furthermore, the methodologies developed in this study could serve as a model for other areas of livestock management, suggesting that similar approaches could be adapted for use with other farm animals such as pigs, cattle, and sheep. This possibility opens up an exciting avenue for future research and development, where the principles of precision farming and welfare-oriented management could be applied across diverse animal husbandry contexts.

### 7.3. Significance for Practical Applications and Contributions to the Field

From a practical standpoint, the implications of this research are vast. For farmers, the ability to understand and quickly respond to the emotional states of chickens could mean a reduction in mortality rates and an improvement in the overall health of the flock. Implementing these insights into decision support systems could lead to more personalized care for animals, optimizing conditions to meet their specific needs and improving the efficiency of operations. For instance, if a particular area of a farm consistently triggers distress calls, adjustments can be made either to the environment or how chickens are managed in that area.

### 7.4. Vision for a Chicken Translator

Looking to the future, the idea of developing a “chicken translator” is a promising avenue for bridging the communication gap between humans and poultry. Such a tool would not only decode the basic emotional states but could also provide insights into the health, well-being, and possibly even the needs of chickens. This technology, powered by AI and NLP, could transform poultry farming by allowing farmers to understand and cater to the vocal expressions of chickens, leading to a revolution in the way we manage animal welfare.

### 7.5. Limitations and Influencing Factors

While promising, the current study’s methodologies do have limitations. One of the primary challenges is the interference of background noise, which can significantly affect the accuracy of vocalization analysis. Additionally, the emotional range of chickens is complex, and while the study successfully categorizes some basic emotions, there is a spectrum of emotional states that may still be undetected. The quality of the data, particularly the clarity and consistency of the audio recordings, also plays a crucial role in the performance of the NLP model.

### 7.6. Future Research Directions

Future research should aim to refine the accuracy and robustness of emotion detection models by incorporating larger and more varied datasets, possibly collected from different breeds and under various environmental conditions. Integrating multimodal data, such as combining vocal analysis with visual observations or physiological measurements, could enhance the model’s reliability and depth of analysis. Moreover, developing algorithms that can adapt to and filter out background noise would be critical for applying these technologies in typical farm environments, which are often noisy.

## 8. Conclusions

The study offers groundbreaking insights into the field of poultry vocalizations, utilizing a Natural Language Processing approach to assess the welfare of chickens. By decoding the subtleties of chicken communications, the research provides a novel method for monitoring and enhancing animal welfare, aligning with the burgeoning field of precision livestock farming. This innovative approach stands to transform traditional practices, shifting toward non-invasive, real-time monitoring that emphasizes the psychological well-being of animals alongside their physical health. The main findings reveal that chicken vocalizations convey a rich knowledge of emotional and physiological states, from distress and fear to contentment and social engagement. These vocal patterns serve not just as a means of communication among chickens but as indicators of their overall welfare. By applying advanced acoustic analysis and NLP techniques, the study deciphers these vocalizations, offering a window into the emotional states of chickens that was previously obscured by more invasive or interpretatively limited methodologies. The current approach represents a significant stride forward in the field of poultry welfare. It goes beyond merely categorizing the types of vocalizations made by chickens, delving into the subtle emotional nuances inherent in these calls. This holistic approach offers a richer comprehension of chicken welfare, shedding light on how various farm management practices influence animal health.

These findings are placed within the broader context of scientific knowledge, suggesting that enhanced understanding and monitoring of animal vocalizations can lead to better welfare practices. This aligns with the trend towards more humane and scientifically informed approaches in agriculture, particularly under the umbrella of smart farming. Here, technology serves not only to increase efficiency and productivity but also to foster an environment where animal welfare is paramount. The practical applications of this research are vast and varied. For farmers, the ability to interpret the needs and states of their flocks in real-time can lead to more precise management decisions, improving both the welfare of the animals and the efficiency of the farming operations. This could potentially manifest in systems that alert farmers to changes in flock behavior or health, prompting immediate action to mitigate issues before they become severe.

Looking to the future, the notion of developing a “chicken translator” seems increasingly plausible. Such a tool would not only enhance our understanding of poultry communications but could also transform poultry management by providing continuous, real-time monitoring and assessment of chicken welfare. This could eventually lead to systems that interpret a range of animal vocalizations, supporting welfare improvements across various livestock sectors. However, the study is not without its limitations. The potential for background noise interference and the complexity of interpreting the emotional range of chickens suggests that further refinement of the technology is necessary. Future research should focus on expanding the datasets to include a wider variety of chicken breeds and environmental conditions, improving the robustness and accuracy of the emotion detection models.

## Acknowledgments

The author thanks the Agriculture and Agri-Food Canada (AAFC) and the Natural Sciences and Engineering Research Council of Canada (NSERC) for funding our study.

